# Allosteric activation of T-cell antigen receptor signalling by quaternary structure relaxation

**DOI:** 10.1101/2020.12.02.407882

**Authors:** Anna-Lisa Lanz, Giulia Masi, Nicla Porciello, André Cohnen, Deborah Cipria, Dheeraj Prakaash, Štefan Bálint, Roberto Raggiaschi, Donatella Galgano, David K. Cole, Marco Lepore, Omer Dushek, Michael L. Dustin, Mark S. P. Sansom, Antreas C. Kalli, Oreste Acuto

## Abstract

The mechanism of T cell antigen receptor (TCR-CD3) signalling remains elusive. Here, we identified mutations in the transmembrane region of TCRβ or CD3ζ that augmented pMHC-induced signalling, not explicable by enhanced ligand binding, lateral diffusion, clustering or co-receptor function. Using a novel biochemical assay and molecular dynamics simulation, we demonstrated that the gain-of-function mutations loosened interaction between TCRαβ and CD3ζ. We found that, similar to the activating mutations, pMHC binding reduced TCRαβ cohesion with CD3ζ. This event occurred prior to CD3ζ phosphorylation and at 0°C. Moreover, we demonstrated that soluble monovalent pMHC alone induced signalling and reduced TCRαβ cohesion with CD3ζ in membrane-bound or solubilised TCR-CD3. Our data provide compelling evidence that pMHC binding suffices to activate allosteric changes propagating from TCRαβ to the CD3 subunits, reconfiguring interchain transmembrane region interactions. These dynamic modifications could change the arrangement of TCR-CD3 boundary lipids to licence CD3ζ phosphorylation and initiate signal propagation.

## Introduction

Signalling through the TCR-CD3 complex drives thymocyte maturation into immunocompetent T cells and T cell response to foreign antigens (Stritesky et al., 2012). These processes initiate upon TCR-CD3 ligation by highly polymorphic major histocompatibility complex (MHC) proteins carrying short peptides (p) originated from the degradation of self and foreign proteins. TCR-CD3 allows T cells to respond with exceptional specificity and sensitivity (Davis et al., 2007) to membrane-bound pMHC ligands of a virtual continuum of weak *K*_d_ (0.1-100 μM) and t_1/2_ of < 0.5 to several seconds (Aleksic et al., 2012; Cole et al., 2007; Stone et al., 2009) and ligand-receptor interfaces of diverse shape and chemical reactivity. To accomplish this task, TCR-CD3 employs a clonally distributed αβ disulphide-linked dimer (TCR) with Ig-like variable domains, Vα and Vβ. VαVβ contains the pMHC binding site composed of six loops homologous to antibody complementarity determining regions (CDRs) 1, 2 and 3 (Garboczi et al., 1996; Garcia et al., 1996). Germline-encoded CDR1 and CDR2 have limited variability, while CDR3s are hypervariable. VαVβ orientates diagonally relative to the long axis of the peptide-binding groove (Garboczi et al., 1996; Garcia et al., 1996), with CDR3s contacting mainly the peptide and CDR1s, and CDR2s contacting primarily the MHC (Baker et al., 2012; Garcia et al., 2012; Marrack et al., 2008). Vα and Vβ are joined to Ig-like constant domains, Cα and Cβ, that are linked to transmembrane regions (TMRs) via a stalk connecting peptide (CP). pMHC binding is signalled intracellularly by four non-covalently associated subunits (γ, δ, ε. and ζ), called CD3, organised into three dimers, γε, δε and ζζ, the latter disulphide-linked (Call et al., 2002)ε,γ and δ each exhibits an Ig-like extracellular domain (ECD) connected to their TMRs by short CPs, while ζ features a ≈ 10 residue-long ECD. A recent TCR-CD3 cryo-electron microscopy (EM) structure at 3.7 Å (Dong et al., 2019) largely reconciles with mutational and NMR studies (Call et al., 2002; He et al., 2015; Mariuzza et al., 2020; Natarajan et al., 2016) but reveals also unsuspected features. VαVβ projects forward while Cα interfaces with CD3*δ*.ECD, Cβ interfaces with CD3*γε*. and CD3*δ*. ECDs, and CD3*ε*. and CD3*ε*. (of δε) ECDs contact each other. Whilst the TMRs of both ζζ subunits (ζ_1_ζ_2_) and of αβ interact with each other, .δε is contacted by α and ζ_1_, and. is contacted by β and ζ_2_. The CPs of α. δ and the ECD of ζ_1_ are stabilised via polar interactions. This highly interlaced structure suggests a mutualistic contribution of each dimer to TCR-CD3 topology and cohesion. The intrinsically disordered intracellular tails of ε, γ, δ and ζ, invisible in the cryo-EM structure, contain immunoreceptor tyrosine-based activation motifs (ITAMs) that become phosphorylated by constitutively active Lck kinase (Nika et al., 2010) within ≤ 1 sec after pMHC binding (Acuto et al., 2008; Huse et al., 2007). The CD3 tails are anchored to the plasma membrane (PM) via basic amino acid residues and ITAM tyrosines that interact with negatively charged lipids and hydrophobic interactions, respectively (DeFord-Watts et al., 2009; Xu et al., 2008), perhaps preventing ITAM phosphorylation of unliganded receptor by Lck. Early studies indicated that agonist anti-CD3 Ab induces exposure of CD3 cytoplasmic tails, presumably by conformational changes triggered by the Ab binding (Gil et al., 2002). However, crystallographic studies of pMHC bound to isolated TCRαβ ECD found considerable conformation changes in the CDRs (Baker et al., 2012; Garcia et al., 2012) but no unambiguous or consistent changes beyond the TCRαβ binding site. This led to put forward signalling models independent of conformational changes or in which pMHC binding alone was insufficient to induce conformational changes of TCR-CD3. These models have proposed that pMHC-induced TCR-CD3 clustering (Cochran et al., 2001; Yokosuka et al., 2005), co-receptors (CD8/CD4) recruitment (Delon et al., 1998) or segregation of the tyrosine phosphatase CD45 (Davis and van der Merwe, 2006) initiated ITAMs phosphorylation and T cell activation. Alternatively, mechanosensing-based models have suggested that force generated by PM movements acts on pMHC-bound TCR-CD3 to induce conformational changes and signalling (Kim et al., 2009; Liu et al., 2014). Finally, it was proposed that clustering by pre-existing pMHC dimers drives conformational changes in CD3ε, but not.directly in TCRαβ (Gil et al., 2002; Minguet et al., 2007). Nevertheless, one crystal structure (Kjer-Nielsen et al., 2003) and a fluorescence-based study (Beddoe et al., 2009) provided evidence that pMHC binding induced a conformational change in a Cα loop. Moreover, deuterium-exchange (Hawse et al., 2012) and recent NMR investigations (Natarajan et al., 2017; Rangarajan et al., 2018) have inferred changes in conformational dynamics of soluble TCRαβ ECD bound to pMHC. These changes mapped to where Cα and Cβ interface with the ECD of the CD3 subunits (He et al., 2015; Natarajan et al., 2016). Although of great appeal, these studies do not rule in or out models proposed thus far; nor do they prove that allosteric effects propagate from αβ to the CD3 subunits for signalling to occur. To challenge this impasse, we conceived a genetic perturbation analysis that should help discriminate first between models requiring or not molecular flexibility (i.e., conformational changes).Towards this goal, we questioned the functional role of αβ TMR, as a key portion of the entire complex establishing physical connection between the pMHC-binding module and the CD3 subunits governing signal delivery into the intracellular milieu. If TMRs are exclusively required for TCR-CD3 solvation within the lipid bilayer and for quaternary structure topology, mutations should not change TCR-CD3 intrinsic signalling capability. In contrast, this could happen in mechanisms based on allosteric interaction or force. We gathered compelling evidence for TMR mutations in TCRβ and CD3ζ that slightly modified the quaternary structure cohesion and augmented intrinsic signalling output. We also found that cohesion changes in TCR-CD3 quaternary structure and signal transduction were induced by soluble monomeric agonist pMHC, independently of co-receptor, clustering or force. We propose that allosteric activation of the T cell antigen receptor by pMHC binding is the prime mover of T cell activation.

## Results

### Gain-of-function mutations in β TMR

To question whether structural alterations in the TMR of the TCRαβ ligand-binding module affected signalling, we employed 1G4, an HLA-A2-restricted TCR specific for the 157-165 peptide from the NY-ESO-1 tumor antigen (Chen et al., 2005). Most residues of TCRβ TMR were individually replaced by alanine or leucine and the corresponding mutants tested for reconstituting TCR-CD3 surface expression in the TCRβ-deficient 31.13 Jurkat cell line (J31.13) (Fig. **1A**). As reported earlier, β K287 mutation substantially reduced TCR-CD3 surface expression (Alcover et al., 1990). However, alanine substitution at βY281, βL285, βG286, βT289, βL290, βY291, βS296 and Leu at βA292 showed only a ≈ 20 % to 40 % decrease of surface expression. Next, the majority of mutants showing 0 % to 40 % reduction of surface expression were co-expressed together with WT 1G4 TCRα in J31.13 and Erk activation (pErk) was monitored after stimulation with 6V-HLA-A2 tetramer (6V-A2)_4_ (Fig. **1B**). While no mutation significantly reduced Erk activation, both βA290 and βA291 significantly increased pErk. A gain-of-function was unexpected, even more so as βA290 and βA291 reduced TCR-CD3 surface expression (the data in Fig. **1B** are not normalised for TCR-CD3 surface expression).

**Figure 1.**
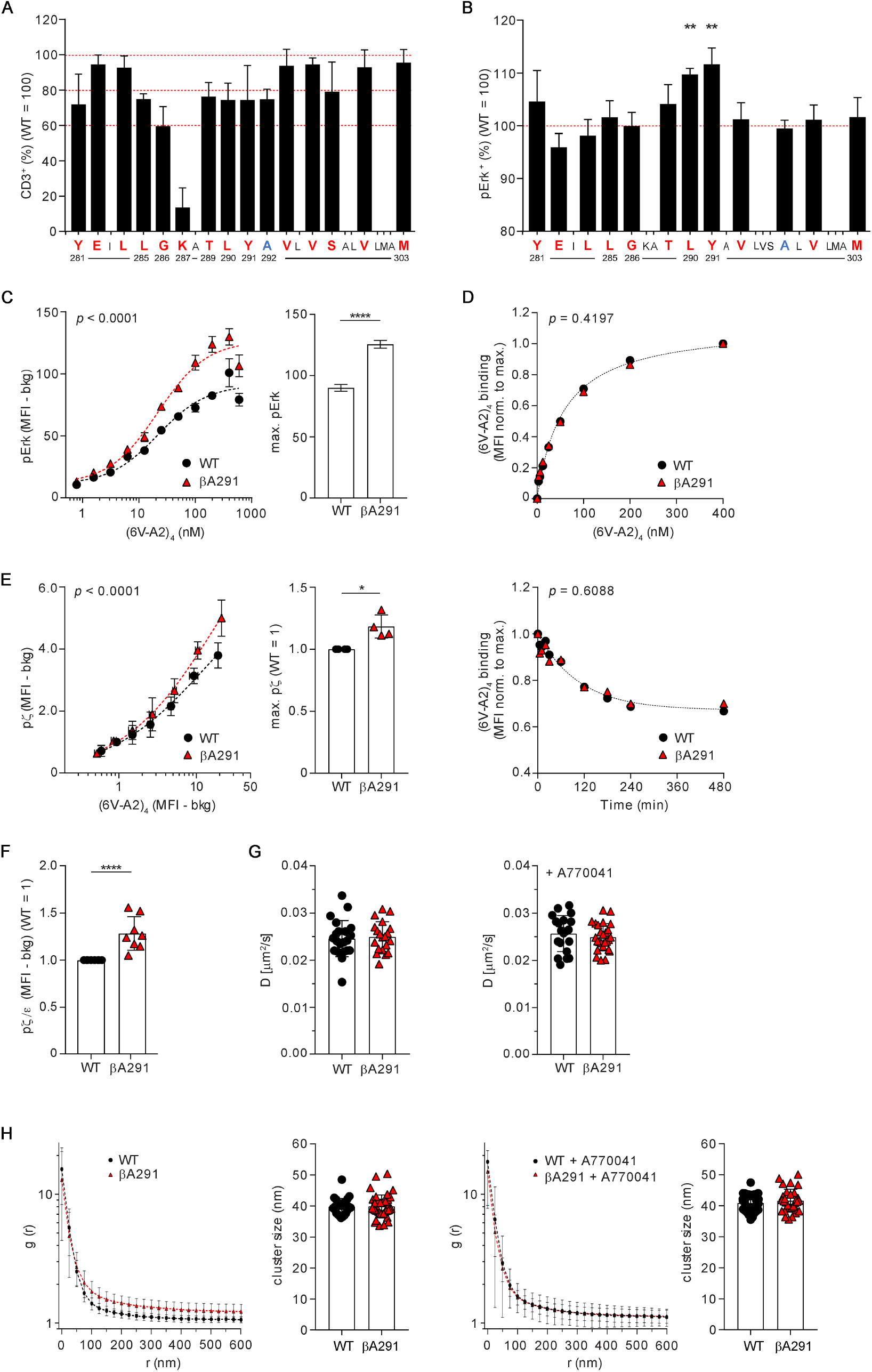
Gain-of-function mutations in β TMR. **A** CD3 surface expression of 1G4-WT and mutants. 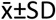 of CD3^+^ cells, n=3-8. Ala (red), Leu (blue) substitution. **B** pErk response of 1G4-WT and β mutants stimulated with (6V-A2)_4_. 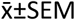 of pErk^+^ cells, n=3-6, unpaired *t*-test *p*=0.0011 (βA290), *p*=0.0092 (βA291). **C** pErk response of CD8-deficient J76 1G4-WT and 1G4-βA291 stimulated with the indicated concentrations of (6V-A2)_4_. **Left**, non-linear regression fit of (6V-A2)_4_ nM vs. pErk MFI, n=3, R^2^=0.82 (WT), 0.89 (βA291); EC_50_=19.5±5.5 (WT), 19.0±3.1 (βA291). **Right**, 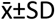 of max. pErk, n=3, F-test *p*<0.0001. See also Fig. **S1E. D** (6V-A2)_4_ binding to 1G4-WT or 1G4-βA291. **Top**, (6V-A2)_4_ dose-dependent association, n=3, non-linear regression fit, R^2^=0.98 (WT), 0.97 (βA291), F-test (ns). **Bottom**, (6V-A2)_4_ dissociation rate, n=5, non-linear regression fit, R^2^=0.84 (WT), 0.72 (βA291), F-test (ns). **E** pζ response of J76 1G4-WT or 1G4-βA291 stimulated with (6V-A2)_4_. **Left**, non-linear regression fit of (6V-A2)_4_ MFI vs. pζ MFI, n=3, R^2^=0.95 (WT), 0.96 (βA291). **Right**, 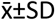. pζ, n=4, unpaired *t*-test *p*=0.0078. See also Fig. **S1F. F** Basal pζ in J76 1G4-WT or 1G4-βA291. pζ MFI normalised to surface CD3 MFI, n=8, unpaired *t*-test *p*<0.0001. See also Figs. **S1H** and **S1I. G** FRAP of 1G4-WT or 1G4-βA291, treated (**right**) or not (**left**) with A770041. 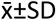 of diffusion coefficient, D (μm^2^/s), n≥20 cells, *t*-test (ns). **H** Lateral distribution by dSTORM of 1G4-WT or 1G4-βA291, treated (**right**) or not (**left**) with A770041. Plots represent pair auto-correlation analysis (g), 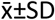 of ≥25 cells. Histograms show DBSCAN cluster analysis, 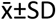 of cluster size per cell, *t*-test (ns).

### βA291 heightens basal and ligand-induced signalling

To validate this apparently paradoxical observation, we focused on βA291 and modified the experimental set up to improve data robustness. Thus, α and β of 1G4 were expressed as a single self-cleavable polypeptide (Fig. **S1A**) from a doxycycline (dox)-inducible promoter in J76, a TCRαβ-deficient Jurkat cell line (STAR Methods). J76 expressed maximum levels of surface TCR-CD3 after 16-18 h of dox treatment and were tested soon after to reduce potential risk of phenotypic drift of cells expressing 1G4 carrying βA291 (hereafter, referred to as 1G4-βA291). As in 31.13 cells, 1G4-βA291 expressed in J76 showed reduced surface expression (≈ 30 %) (cf. Fig. **1A** with Fig. **S1B**). However, in most experiments we lowered the dox concentration when inducing 1G4-WT in order to reduce the difference in surface expression with 1G4-βA291 (to < 5 %) (Fig. **S1C**). Moreover, in most flow-cytometry analyses, J76 expressing 1G4-WT or mutant were bar-coded by labelling with CellTrace™ violet, mixed before stimulation and analysed simultaneously. These stratagems considerably simplified and made more robust the computation of differences in signalling output between WT and mutant. Erk activation was retained as a sensitive and reliable read-out of TCR-CD3 signal transduction and propagation as it reports the occurrence of a cascade of early signalling steps, including ITAM phosphorylation, ZAP-70 activation, LAT signalosome assembly and PLCγ1 activation that generates IP_3_ (for intracellular [Ca^2+^] increase) and DAG required for Ras activation by Ras-GRP (Acuto et al., 2008). Titration of (6V-A2)_4_ showed a shift in pErk response by 1G4-βA291 towards higher sensitivity, but also revealed a significant higher Erk activation (Fig. **1C**). This was not due to a higher Erk activation ceiling in 1G4-βA291-expressing cells (Fig. **S1D**) nor to augmented binding of (6V-A2)_4_ to 1G4-βA291 (Fig. **1D**, upper panel and lower panels), but it was consistent with the dose-response plot showing unchanged EC_50_ between 1G4-βA291 and 1G4-WT (Fig. **1C** and see Method for computation). The higher maximal response of 1G4-βA291 was compatible with a faster proofreading rate (*k*_p_) for a receptor operating in a kinetic proofreading regimen (McKeithan, 1995). Indeed, fitting the data of Fig. **1C** into a minimal model of kinetic proofreading (Dushek et al., 2011) showed the *k*_p_ for 1G4-βA291 was considerably higher than 1G4-βWT (Fig. **S1E** and STAR Methods), consistent with βA291 enhancing TCR-CD3 intrinsic signalling capability (i.e., enhancing ligand potency). Note that the gain-of-function was observed in CD8-deficient J76 compared to CD8-deficient J76 expressing WT (Fig. **1C**), ruling out that the βA291 mutation enhanced TCR-CD3 interaction with co-receptor. Augmented signalling was also evident for ζ phosphorylation (pζ) (Figs. **1E** and **S1F**), the earliest intracellular signalling event. Remarkably, anti-CD3ε (UCHT1) Ab stimulation of 1G4-βA291 also heightened pζ (Fig. **S1G**), a triggering modality that by-passes pMHC binding, further supporting that βA291 enhanced TCR-CD3 signalling output. These data suggested that βA291 might increase constitutive TCR-CD3 signalling that can be detected by measuring pζ in non-stimulated cells. Indeed, pζ was significantly higher basally in cells expressing 1G4-βA291 as compared to 1G4-WT (Figs. **1F** and **S1H**) and it was TCR-signal specific as it disappeared after treatment by A770041 (Stachlewitz et al., 2005), a potent and highly specific inhibitor of Lck (Fig. **S1I**). We then asked if βA291 increased signalling by influencing TCR-CD3 lateral diffusion and/or distribution. However, fluorescence recovery after photo-bleaching (FRAP) found no significant difference in the diffusion coefficient (D) between 1G4-βA291 and 1G4-WT (Fig. **1G**, left panel), which remained unchanged after A770041 treatment (Fig. **1G**, right panel). dSTORM super-resolution microscopy found no statistically significant difference in the cluster size distribution formed by 1G4-βA291 and 1G4-WT (histograms in Fig. **1H**). Although not statistically significant, the reproducible small increase of larger cluster frequency for 1G4-βA291 disappeared after A770041 treatment (cf. auto-correlation function plots in left and right panels of Fig. **1H**), indicating it to be secondary to 1G4-βA291 heightened basal signalling (Fig. **1F**), rather than βA291 causing it. Finally, we questioned the potential cause(s) of mildly reduced 1G4-βA291 surface expression. We excluded that βA291 reduced β protein expression (Fig. **S1J**) and considered that heightened basal signalling might decrease receptor surface expression by increasing its down-regulation rate. However, exposure to A770041 for several hours increased surface expression of both 1G4-βA291 and 1G4-WT in similar proportion (≈ 20 %) but did not significantly reduce their difference (Fig. **S1K**). These data led us to consider if βA291 modified the stability of TCR-CD3 quaternary structure that could mildly reduce export to the PM due to increased negative triage of mutant vs. WT by protein quality control systems (Feige and Hendershot, 2013).

### βY291 contribution to TCR-CD3 quaternary structure cohesion

Non-ionic detergents used at high concentration to quantitatively extract TCR-CD3 can dissociate TCRαβ from the CD3 modules (Testi et al., 1989). Presumably, this can be attributed to the substitution of natural boundary lipids by the detergent, with possible interference with TMR inter-helical interactions that are critical for TCR-CD3 quaternary structure cohesion (Alcover et al., 1990; Call et al., 2002; Dong et al., 2019). However, 0.5 % of the non-ionic detergent n-Dodecyl-β-D-Maltopyranoside (DDM) allows quantitative extraction of stoichiometrically intact TCR-CD3 (Swamy et al., 2008) (Fig. **S2A**). Thus, if βA291 altered TCRαβ cohesion with CD3 by unsettling TMR inter-helical interactions, 0.5 % DDM extraction may show lower recovery of intact 1G4-βA291 with respect to 1G4-WT. Figure **S2B** illustrates the conceptual and experimental set up of chemically probing TCR-CD3 cohesion by DDM that we named DDM Stability Assay (DSA) (see STAR Methods for further details). Membrane solubilisation by 0.5 % DDM and pull-down (PD) of total β (mostly associated with α (Alcover et al., 2018) by the β-HA tag was followed by quantitative immunoblot (IB) for β (with anti-HA Ab) and for each CD3 subunit. Anti-HA IB identified three β isoforms (named β_1_, β_2_ and β_3_, Fig. **2A**). β_3_ was the endo-H-sensitive ER-resident β isoform (Fig. **S2C**) that is assembled with α, γε, δ ε but not with ζζ (Alcover et al., 2018), as confirmed by β_3_ being undetected in CD3ζ PD (Fig. **S2D**). β_1_ and β_2_ were both endo-H-resistant (Fig. **S2C**), though β_2_ was the only β isoform associated with ζζ (Fig. **S2D**). Thus, to evaluate the effect of βA291 on TCR-CD3 complex cohesion we used the IB signals for the β isoforms, ζζ and ε (including γε and δε). When ζ/β_2_ was set equal to 1 for 1G4-WT (i.e., 100 % recovery of intact TCR-CD3), reduced cohesion between ζζ and αβ in 1G4-βA291 should result in ζ/β_2_ < 1 (Fig. **S2B**). The DSA showed that ζ/β_2_ for 1G4-βA291 was 0.2, indicating only 20 % recovery of intact TCR-CD3 (or 80 % loss of ζζ recovery) after DDM solubilisation (Fig. **2A**). To determine the effect of βA291 on γε and δ ε cohesion with αβ. we used instead the sum of β_1_, β_2_ and β_3_ (or total β .β_T_) IB signals that represented cytoplasmic and PM αβ,most of which is associated with ε (Alcover et al., 2018). ε /β_T_ for 1G4-βA291 was ≈ 0.5 indicating ≈ 50 % reduced recovery of ε (Fig. **2A**). These ratios did not change after A770041 treatment during dox induction of αβ expression (Fig. **S2E**), excluding that reduced recovery concerned the pool of 1G4-βA291 with increased ζ basal phosphorylation (Fig. **1F**). The considerable reduction of ζ (80%) and ε(50%) recovery for 1G4-βA291 could not be the consequence of severance of the same magnitude of αβ from ζζ (or from δε and γε dimers) before export to the PM and/or at the PM, when considering a mere 20-30% reduction of TCR-CD3 expression observed by flow cytometry. And, even more so compatible with increased pMHC-induced signalling. Indeed, when TCRαβ is no longer in contact ζζ, TCRαβεγε alone cannot be exported to the T cell surface (Fig. **S2F** and (Alcover et al., 2018). Therefore, in the PM natural lipid environment, βA291 only slightly perturbed TCR-CD3 quaternary structure cohesion (as the MDS shows, see below), minimally reducing surface expression. However, substitution of natural boundary lipids by DDM severely corroded TCR-CD3 cohesion in 1G4-βA29 and provoked partial physical detachment of ζ and ε from αβ during the solubilisation. IB for γ and δ revealed that βA291 affected both γε and δεcohesion with the rest of the complex though asymmetrically, as it reduced γε and δε recovery of 40 % and 10 %, respectively (Figs. **2B** and **2C**). In the cryo-EM structure, βY291 (note that Dong *et al*. (Dong et al., 2019) refer to βY291 as βY292) contacts mostly γε and therefore βA291 can be expected to affect primarily the interaction between αβ and γε in accordance with the DSA. However, βY291 makes no contacts with ζζ and δε (Dong et al., 2019) (and see also MDS below). Therefore, the DSA revealed a more complex picture, with βA291 presumably affecting indirectly the interaction of both ζζ and δε with the rest of the complex. To further understand the structural role of βA291, we used the TMRs’ atomic coordinates of the cryo-EM structure of the TCR-CD3 octamer (PDB: 6JXR) (Dong et al., 2019) to carry out all-atom molecular dynamics simulations (MDS) with βWT and βA291 in an asymmetric lipid bilayer, mimicking the lipid environment of TCR-CD3 (see STAR Methods for details) and adding dynamical insight into TCR-CD3 cohesion. Simulations for 1250 ns confirmed considerable contacts of β WT with ε (of γε), γα, and ζζ but not with δε (Figs. **2D** and **S2G**) and revealed one new contact of β with ζζ as well as significant reduction in six β contacts with ε (γε), five with γ and four with α (Fig. **S2G**). Specifically, during the simulations, significant contacts of βY291 with αN263, αT267, γL129, γG132, and εL145 were observed (Figs. **2D** right panel and **S2H**) and also with γV133 and γI136, though not considered significant on the normalised scale (Fig. **S2H**). No contacts of βY291 with ζζ were seen (Fig. **S2G**). Simulations of the TMR octamer carrying βA291 indicated new and augmented contacts of β with ε (γε) and γ (Fig. **S2I**). In addition, while βA291 still contacted γL129, it completely lost interaction with γG132, γV133 and γI136 (Fig. **S2J**, middle panel). Likely, these changes were secondary to spatial re-adjustments due to the loss of the bulky tyrosine side chain. No contacts of βA291 with ζζ were observed. Overall, the simulations suggested that βA291 reshuffled contacts with γε, with the net effect of increasing local compaction (Fig. **S2K**), as also indicated by a stabilisation of their α-helices crossing angle (Fig. **S2L**). This result seemed to contradict the DSA data of βA291 severely affecting ζζ interaction with the rest of the complex. Although 1250 ns time-scale is relatively long for all-atom simulations of membrane proteins, it might be insufficient to capture re-adjustments of interchain contacts that possibly occur at larger time-scales. βA291 might affect the role of interfacial lipids in cementing α-helices interactions (Gupta et al., 2017) that when challenged with DDM could cause crumbling of TMRs’ cohesion in the mutant, despite augmented compaction by βA291 elsewhere. However, reduced export to the PM was a good indicator that βA291 (and other β and ζTMR mutants, see below) promoted some instability of the complex, causing dynamical exposure of hydrophobic site and/or retention signals, detected and negatively triaged by protein quality control systems (Feige and Hendershot, 2013). Comprehensively, these data suggested a positive correlation between reduced quaternary structure cohesion of TCR-CD3 and signal transduction activation.

**Figure 2.**
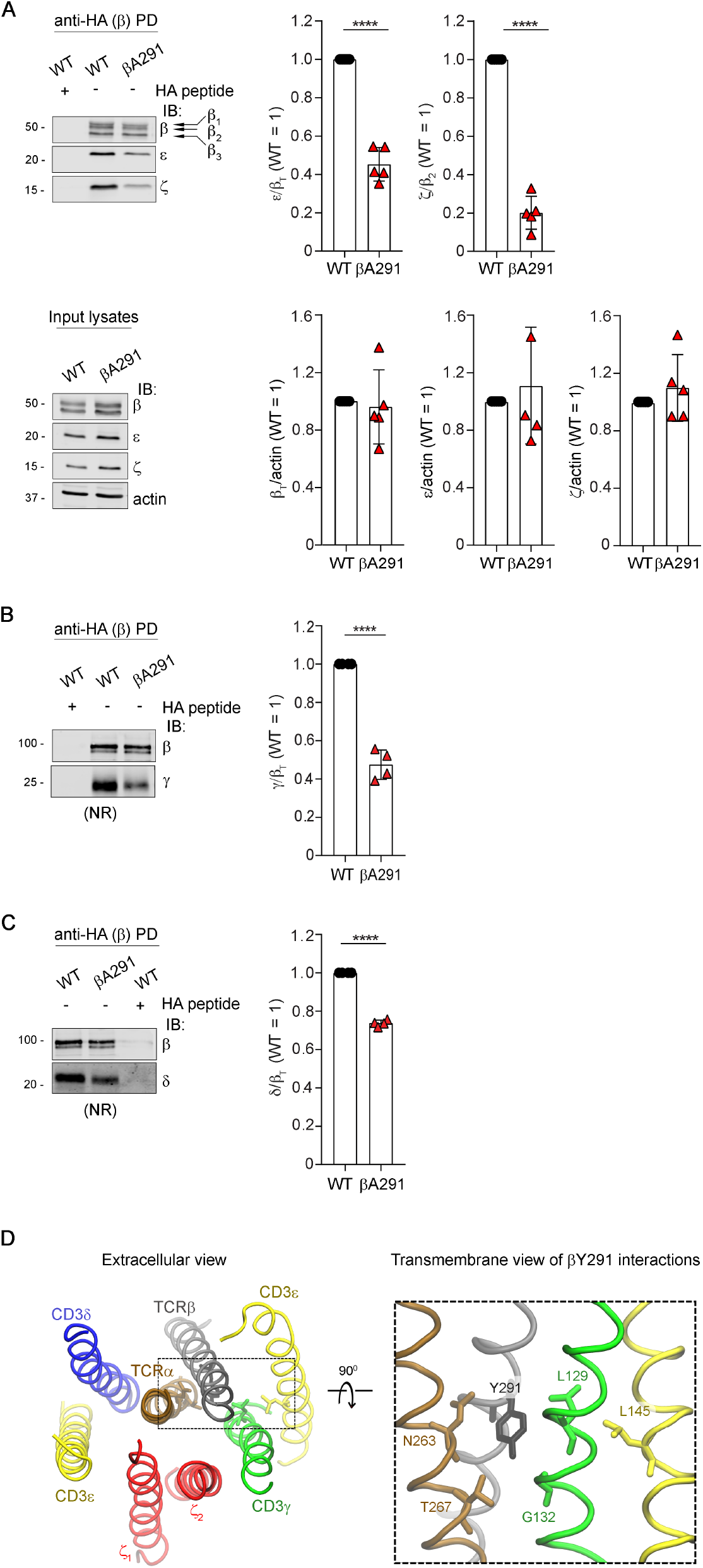
βY291 contribution to TCR-CD3 quaternary structure cohesion. **A** Anti-HA (β-HA) Pull Down (PD) and IB of 1G4-WT or 1G4-βA291. **Upper panels: left**, IB: 1 of 5 experiments, arrows indicate β-isoforms; **right**, 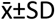 of ε/β_T_ and ζ/β_2_, n=5, unpaired *t*-test *p*<0.0001. **Bottom panels:** input lysates, **left**, IB: 1 of 5 experiments; **right**, 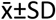 of β_T_/actin, ε/actin and ζ/actin, n=5, unpaired *t*-test (ns). **B** β-HA PD and IB of 1G4-WT or 1G4-βA291. (NR) non-reducing conditions. **Left**, IB: 1 of 4 experiments. **Right**, 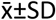 of γ/β_T_, n=4, unpaired *t*-test *p*<0.0001. **C** β-HA PD and IB of 1G4-WT or 1G4-βA291. (NR) non-reducing conditions. **Left**, IB: 1 of 4 experiments. **Right**, 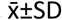 of δ/β_T_, n=4, unpaired *t*-test *p*<0.0001. **D** All-atom MDS of TCR-CD3 TMRs. TCRα (ochre), TCRβ (grey), CD3δ (blue), CD3ε (yellow), CD3γ (green), ζ (red). **Left**, extracellular view of a snapshot of TCR-CD3 TMRs. **Right**, βY291 interactions with TCR-CD3 TMRs. Significant contacts of βY291 with TCRα, CD3γ and CD3ε are shown as liquorice sticks. See also Fig. **S2H**.

### Loosening ζ association enhances signalling

To corroborate this hypothesis, we investigated the phenotype of additional mutations in β and ζ TMRs. We found that similar to βA291, also βF291 and βL291 mildly reduced TCR-CD3 surface expression, despite no decrease in β expression (Fig. **3A**). Both mutations reduced interaction of β with ζ and ε (Fig. **3B**) and augmented pErk maximal response to (6V-A2)_4_ (Fig. **3C** and **3D**), whose binding remained unchanged (Fig. **S3A**). These three readouts ranked according to: βL291 ≥ βA291 > βF291 > WT, presumably reflecting conservative or non-conservative replacements, hence indicating a direct correlation between increased quaternary structure loosening and heightened signalling. We then tested the effect of βA291 in 2H5, an HLA-A2-restricted TCR specific for the MART-1 tumour antigen (MART-1 (Circosta et al., 2009). Similar to 1G4-βA291, 2H5-βA291 showed reduced surface expression (Fig. **3E**) and TCR-CD3 cohesion (Fig. **3F**) and augmented pErk for equal (MART-1-A2)_4_ binding (Figs. **3G** and **S3B**). Conversely, mutation of β TMR residues not involved in critical contacts (Dong et al., 2019), such as βA293 and βA303, showed no significant change of 1G4 surface expression (Fig. **S3C**), TCR-CD3 cohesion (Figs. **S3D** and **S3E**) and no increase in (6V-A2)_4_-induced pErk (Figs. **S3F** and **S3G**).

**Figure 3.**
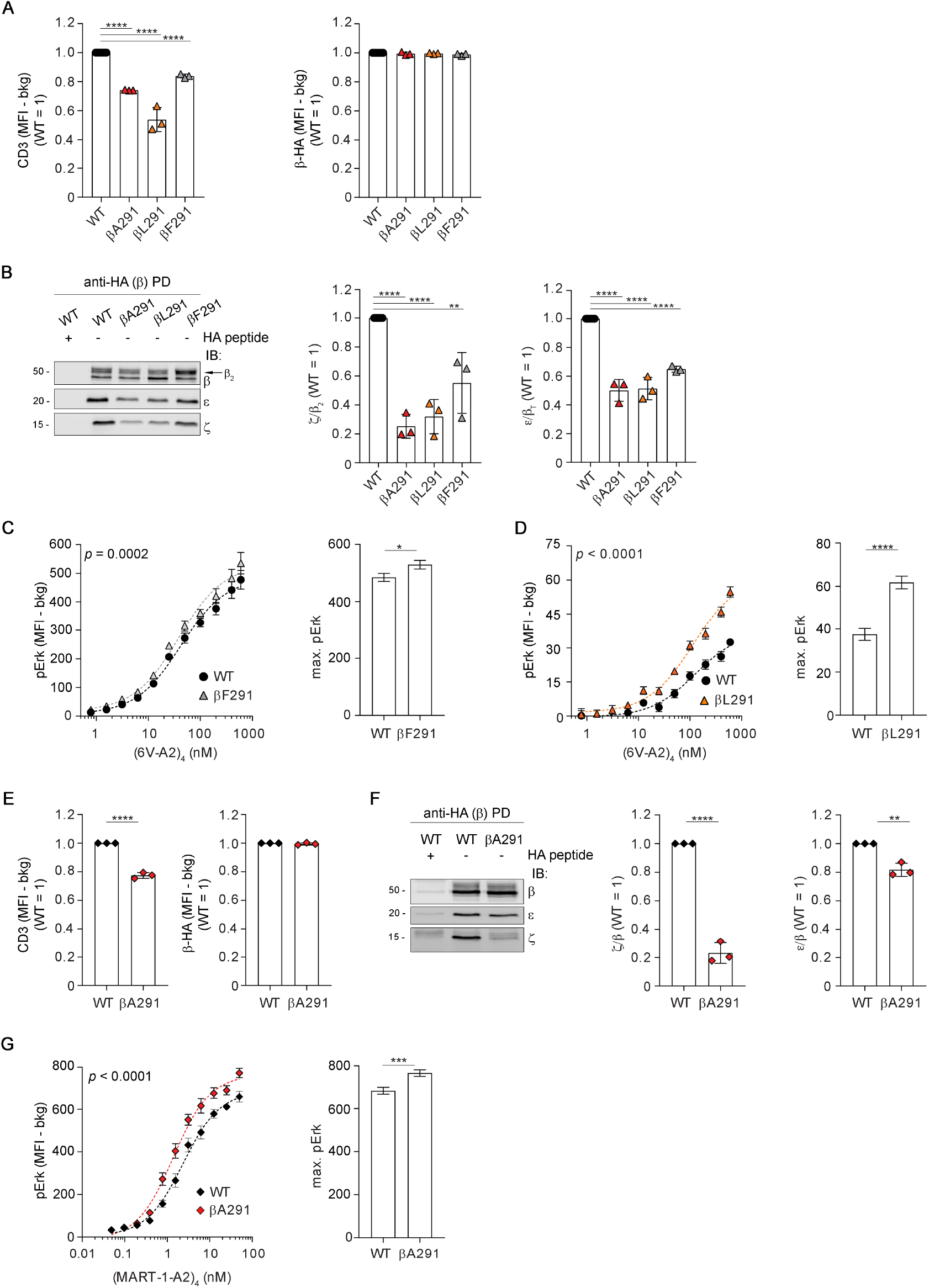
Loosening ζ association enhances signalling. **A** TCR-CD3 expression of 1G4-WT, 1G4-βA291, 1G4-βL291, 1G4-βF291 in CD8-deficient J76. **Left**, 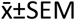 of CD3 MFI in HA^low^gate, n=3, unpaired *t*-test *p*<0.0001. **Right**, 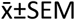 of β-HA MFI in HA^low^gate, n=3, *t*-test (ns). **B** β-HA PD and IB of 1G4-WT or the indicated 1G4-β mutants. **Left**, IB: 1 of 3 experiments. The arrow indicates β_2_ isoform. **Middle**, 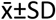 of ζ/β_2_, n=3, unpaired *t*-test WT vs. βA291, WT vs. βL291 *p*<0.0001, WT vs. βF291 *p*<0.01. **Right**, 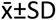 of ε/β_T_, n=3, unpaired *t*-test *p*<0.0001. **C** pErk response of CD8-deficient J76 1G4-WT or 1G4-βF291 stimulated with the indicated concentrations of (6V-A2)_4_. **Left**, non-linear regression fit of (6V-A2)_4_ nM vs. pErk MFI, n=3, R^2^=0.915 (WT), 0.910 (βF291). **Right**, 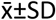 of max. pErk, n=3, F-test *p*<0.05. See also Fig. **S3A** (**left**). **D** pErk response of CD8-deficient J76 1G4-WT or 1G4-βL291 stimulated with the indicated concentrations of (6V-A2)_4_. **Left**, non-linear regression fit of (6V-A2)_4_ nM vs. pErk MFI, n=3, R^2^=0.827 (WT), 0.910 (βL291). **Right**, 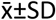 of max. pErk, n=3, F-test *p*<0.0001. See also Fig. **S3A** (**right**). **E** TCR-CD3 expression in CD8-deficient J76 2H5-WT or 2H5-βA291. **Left**, 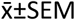 of CD3 MFI in HA^low^gate, n=3, unpaired *t*-test *p*<0.0001. **Right**, 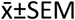 of β-HA MFI in HA^low^gate, n=3, *t*-test (ns). **F** β-HA PD and IB of 2H5-WT or 2H5-βA291. **Left**, IB: 1 of 3 experiments. **Middle**,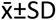 of ζ/β, n=3, unpaired *t*-test *p*<0.0001. **Right**, 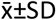 of ε/β, n=3, unpaired *t*-test *p*<0.01. **G** pErk response of CD8-deficient J76 2H5-WT or 2H5-βA291 stimulated with the indicated concentrations of (MART-1-A2)_4_. **Left**, non-linear regression fit of (MART-1-A2)_4_ nM vs. pErk MFI, n=3, R^2^=0.94 (WT), 0.94 (βA291). **Right**, 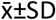 of max. pErk, n=3, F-test *p*<0.001. See also Fig. **S3B**.

βY291 did not contact ζ. but its mutation augmented basal (Fig. **1F**) and ligand-induced ζ phosphorylation (Fig. **1E**) and signal propagation. This was reminiscent of allosteric interaction revealed by mutations (Changeux and Christopoulos, 2016; Volkman et al., 2001) -e.g., mutations at βY291 induced local re-adjustments but also distal functional effects, such as favouring exposure of ζ cytosolic tail to active-Lck. To investigate this possibility, we tested whether mutations in ζ TMR residues susceptible to loosen ζζ contacts with subunits other than αβ phenocopied mutations at βY291. TCR-CD3 cryo-EM structure and MDS indicated that ζ_1_ and ζ_2_ TMRs contacted only the N-terminal moiety of β TMR (Figs. **4A** and **S4A**) and α TMR throughout (Figs. **4B** and **S4B**). However, ζ_2_ and ζ_1_ contacted also γ(Figs. **4C** and **S4C**) and ε(of the δε) (Figs. **4D** and **S4D**), respectively. Specifically, MDS revealed that ζ_1_I38 contacted two residues of ε (of ε) (Figs. **4D** and **S4E**, left panel) and ζ_2_I41 contacted two residues of γ (Figs. **4C** and **S4F**, left panel), whereas ζ_2_I38 and ζ_1_I41 bulged toward the membrane lipids and made no contact with the complex. Thus, ζ_1_I38 and ζ_2_I41 were deemed capable of partially disturbing ζ_1_ and ζ_2_ interactions with ε (of δ ε) and γ, but perhaps not with αβ. To verify this prediction, 1250 ns all-atom simulations of TCR-CD3 octamer TMRs composed of ζ WT, ζA41 and ζA38 mutants were carried out. At the end of the simulations, alignment of snapshots of the mutated and WT TMRs showed distortion in the contacts of ζ_1_ with. (Fig. **4E**) and ζ_2_ with. (Fig. **4G**). As a consequence, ζζ containing ζ_1_A38 (3 out of 3 simulations) or ζ_2_A41 (2 out of 3 simulations) increased fluctuation relative to αβ as compared to ζζ WT. This can be appreciated from the average spatial distribution plots of the Cα atoms of ζζ relative to the Cα atoms of αβ that showed broader density for both mutants (Figs. **4F** and **4H**), though more pronounced for ζ_1_A38. These results were indicative of ζA38 and ζA41 increasing ζζ flexibility relative to αβ. Both mutants maintained some ζζ contacts with the rest of the complex (Figs. **S4G** -**S4N**). These results prompted us to test if, similar to the βA291 mutations, also these ζ mutations showed reduced surface expression, complex cohesion by DSA and enhanced signalling. The data showed that ζA38 or ζA41 reduced 1G4 surface expression by ≈ 30 %, for similar ζ expression (Fig. **5A**). The DSA showed that ζA38 and ζA41 reduced ζ/β_2_ ratio to ≈ 0.05 and ≈ 0.25 (95 % and 75 % loss of ζ recovery), respectively, without apparently affecting ε cohesion with αβ. (Figs. **5B** and **S5A**). Thus, the DSA agreed with the loosening of ζζ interaction with αβ. as predicted by the atomistic simulations. We surmise that by weakening direct interactions with CD3 TMRs and causing higher ζζ mobility, ζA38 and ζA41 indirectly loosened ζζ interaction also with αβ, as revealed by the DDM extraction. Simulations of larger time-scales may provide clearer insights into dynamic alterations of ζζ by βA291 that presumably first produced a direct effect on γε and later on ζζ. Importantly, similar to βA291, both ζ mutations conferred to 1G4 heightened pErk response to (6V-A2)_4_, with a higher maximum compared to 1G4-WT (Figs. **5C** and **5D**) for equal (6V-A2)_4_ binding (Figs. **S5B** and **S5C**). We concluded that reduced cohesion between αβ and ζζ caused heightened signalling, rather than the mutations of βY291 *per se*. This strengthened the idea that reducing TCR-CD3 cohesion populated the active signalling state of TCR-CD3 -i.e., it lowered the activation energy between two presumed functional states: inactive and active, the latter initiating transmembrane signalling. These data made unlikely that TCR-CD3 TMRs are just structural elements required for TCR-CD3 membrane solvation and architecture, as conformational change-independent models would imply. Rather, by analogy with allosterically regulated proteins that can be switched on or off by mutations distal from their active site(s) (Changeux and Christopoulos, 2016; Volkman et al., 2001) and considering recent NMR studies (He et al., 2020; Natarajan et al., 2016; Natarajan et al., 2017; Rangarajan et al., 2018), our data suggested that pMHC binding could activate an allosteric cascade that loosened TCR-CD3 cohesion including TMR interactions with ζζ TMRs serving as a second-to-last relay before licencing ζ ITAM phosphorylation. These considerations prompted us to investigate this possibility.

**Figure 4.**
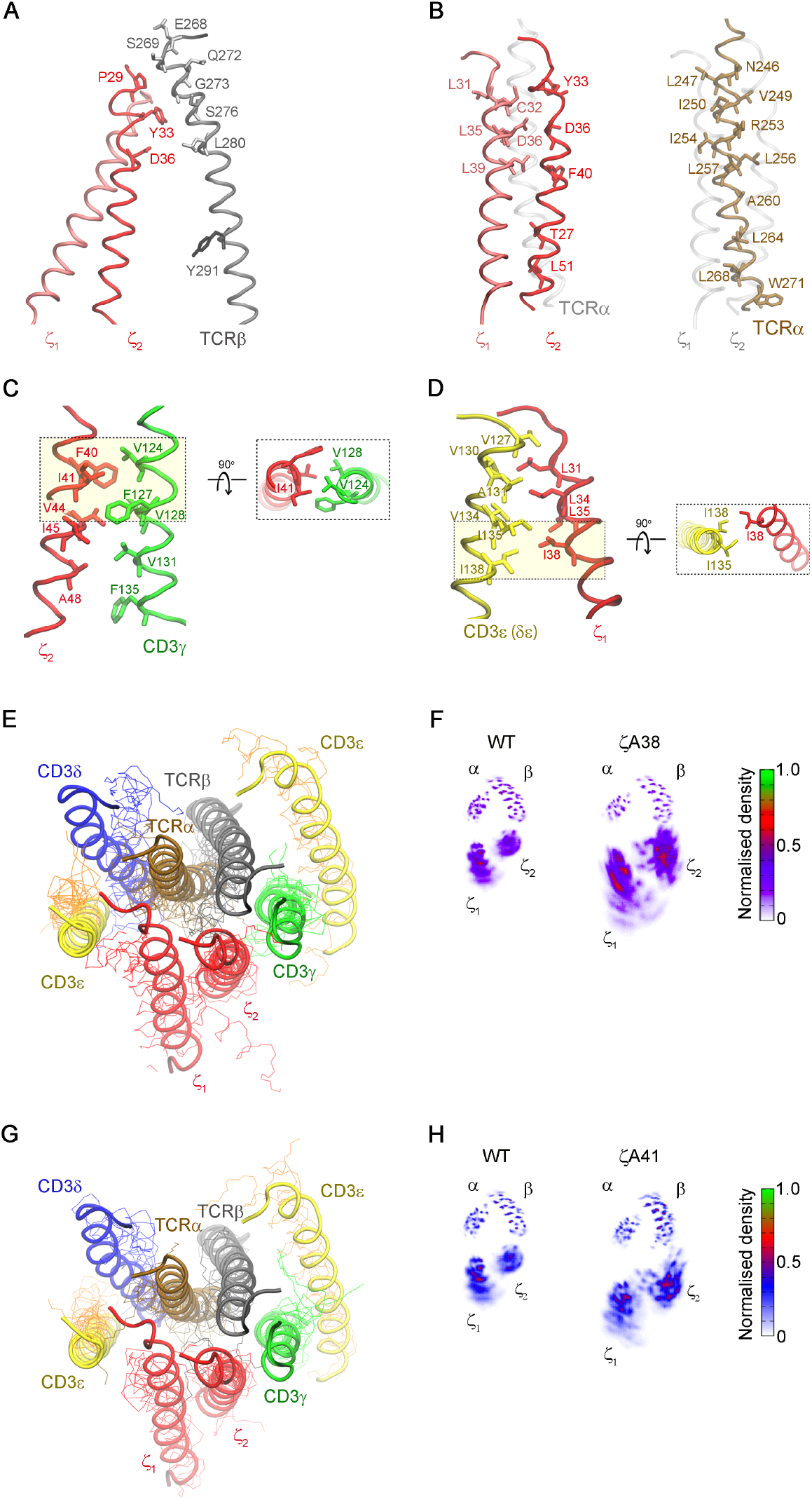
Loosening ζ association enhances signalling. **A** Snapshot from all-atom MDS of TCR-CD3 TMRs. Contacts between TCRβ (grey), ζ_1_ (light red) and ζ_2_ (dark red) TMRs. See also Fig. **S4A**. βY291 is represented as liquorice stick for reference and does not contact ζζ. **B** Snapshot from all-atom MDS of TCR-CD3 TMRs. **Left**, ζ_1_ (light red) and ζ_2_ (dark red) residues contacting TCRα TMR (in transparency). **Right**, TCRα (ochre) residues contacting ζ_1_ and ζ_2_ TMRs (in transparency). See also Fig. **S4B. C** Snapshot from all-atom MDS of TCR-CD3 TMRs. **Left**, contacts between ζ_2_ (red) and CD3γ (green) TMRs. **Right**, Top view of ζ_2_I41 (red) contacts with CD3γ (green). See also Figs. **S4C** and **S4F. D** Snapshot from all-atom MDS of TCR-CD3 TMRs. **Left**, contacts between ζ_1_ (red) and CD3ε (δε) (yellow) TMRs. **Right**, Top view of ζ_1_I38 (red) contacts with CD3ε (δε) (yellow). See also Figs. **S4D** and **S4E. E** Snapshot from all-atom MDS of TCR-CD3 TMRs carrying ζA38 (lines) aligned to ζWT (cartoon) at the end of 1250ns MDS. TCRα (ochre), TCRβ (grey), CD3δ (blue), CD3ε (yellow), CD3γ (green), ζ (red). See also Figs. **S4G**-**S4J. F** Normalised spatial distributions of the Cα atoms of ζζ relative to the Cα atoms of TCRαβ in ζWT and ζA38. **G** Snapshot from all-atom MDS of TCR-CD3 TMRs carrying ζA41 (lines) aligned to ζWT (cartoon) at the end of 1250ns MDS. TCRα (ochre), TCRβ (grey), CD3δ (blue), CD3ε (yellow), CD3γ (green), ζ (red). See also Figs. **S4K**-**S4N. H** Normalised spatial distributions of the Cα atoms of ζζ relative to the Cα atoms of TCRαβ in ζWT and ζA41.

**Figure 5.**
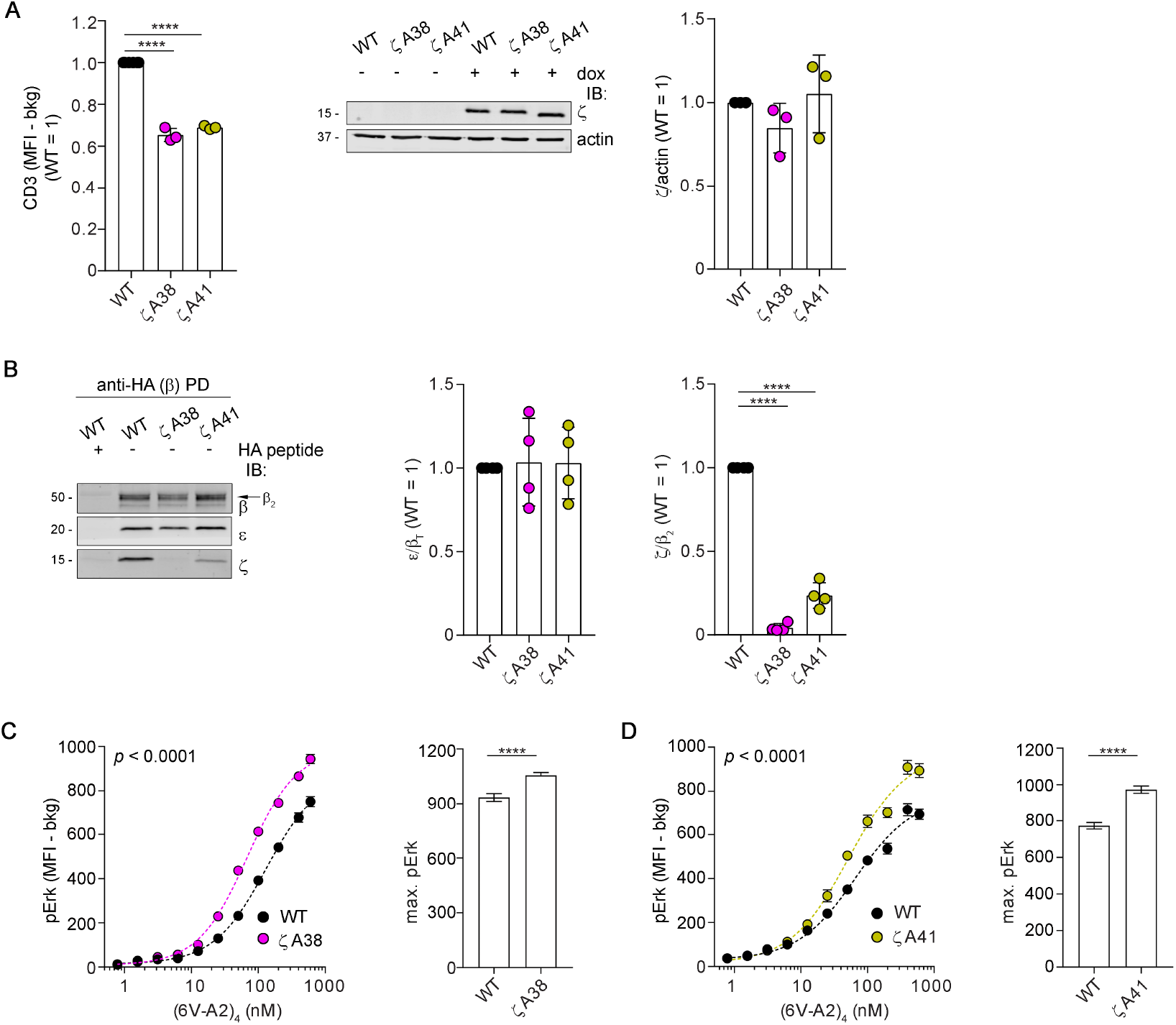
Loosening ζ association enhances signalling. **A** TCR-CD3 expression in J76-1G4WT-ζKO expressing ζWT or ζA38 or ζA41. **Left**, 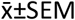 of CD3 MFI in HA^low^ gate, n=3, unpaired *t*-test *p*<0.0001. **Middle**, IB: 1 of 3 experiments. **Right**, 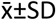 of ζ*/*actin, n=3, unpaired *t*-test (ns). **B** β-HA PD and IB of 1G4-WT carrying ζWT or ζA38 or ζA41. **Left**, IB: 1 of 4 PD. The arrow indicates β_2_ isoform. **Middle**, 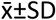 of ε/β_T_, n=4, unpaired *t*-test (ns). **Right**, 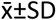 of ζ/β_2_, n=4, unpaired *t*-test *p*<0.0001. See also Fig. **S5A. C** pErk response of J76-1G4WT-ζKO expressing ζWT or ζA38 stimulated with the indicated concentrations of (6V-A2)_4_. **Left**, non-linear regression fit of (6V-A2)_4_ nM vs. pErk MFI, n=3, R^2^=0.98 (WT), 0.99 (ζA38). **Right**, 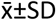 of max. pErk, n=3, F-test *p*<0.0001. See also Fig. **S5B. D** pErk response of J76-1G4WT-ζKO expressing ζWT or ζA41 stimulated with (6V-A2)_4_. **Left**, non-linear regression fit of (6V-A2)_4_ nM vs. pErk MFI, n=3, R^2^=0.96 (WT), 0.97 (ζA41). **Right**, 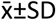 of max. pErk, n=3, F-test *p*<0.0001. See also Fig. **S5C**.

### pMHC tetramer binding loosens αβ association with ζ

Conformational changes produced by pMHC binding may have distal effects and reduce contacts of CαCβ with δε and/or γ ε ECDs, eventually propagating to TMR contacts, including ζζ TMR and ultimately showing reduced TCR-CD3 complex cohesion. If correct, the DSA should show reduced recovery of ζζ in ligand-engaged TCR-CD3 vs. unengaged TCR-CD3. To test this idea, we aimed to capture DDM-solubilised pMHC-engaged TCR-CD3 and compare subunits’ recovery with unengaged TCR-CD3. Thus, 1G4-WT-expressing J76 were briefly stimulated with (9V-A2)_4_, tetramerised with His-tagged streptavidin [(9V-A2)_4_-His] (Fig. **S6A** and STAR Methods), ligand excess was removed and cells rapidly solubilised with 0.5 % DDM. Post-nuclear lysates were incubated with His-Cobalt beads for a quick capture of (9V-A2)_4_-His-bound 1G4. This experimental setting failed to capture sufficient (9V-A2)_4_-His-bound TCR-CD3, likely because detergent solubilisation interfered with the avidity gain due to tetramer-induced clustering in the membrane milieu, by making the dissociation rate of individual 9V-A2 in the tetramer closer to that of a 9V-A2 monomer alone (i.e., a solution *k*_off_ of 0.33 s^-1^ at 25 °C (Aleksic et al., 2010). To overcome this limitation, we initially employed wtc51 (Irving et al., 2012), a 1G4 variant harbouring four mutations in βCDR2 (Fig. **S6B**) that confer a 15 nM *K*_d_ for NY-ESO-1_157-165_-HLA-A2 (*k*_off_ of 0.0013 s^-1^ at 25 °C). Computational modelling showed that NY-ESO-1_157-165_-HLA-A2 adopts a canonical orientation onto wtc51 VαVβ. almost indistinguishable from 1G4-WT (Fig. **S6B**). (9V-A2)_4_-His induced robust wtc51-mediated Erk activation (Fig. **6A**, left panel) and allowed specific capture of engaged wtc51 (Fig. **6A**, middle panel, lanes 2 and 4) to be compared with unliganded wtc51 isolated by anti-HA β pull-down (Fig. **6A**, middle panel, lanes 1 and 3). β_2_ was the only isoform bound to (9V-A2)_4_-His (Fig. **6A**, middle panel, lanes 2 and 4), consistent with it being the only one associated to ζ and present at the cell surface (Fig. **S2D**). Therefore, ζ/β_2_ ratio was used to assess if pMHC binding had reduced cohesion of ζ within TCR-CD3 (Fig. **6A**, right panel). The data showed that ζ/β_2_ in liganded wtc51 was 0.5 (Fig. **6A**, right panel), in agreement with pMHC binding causing TCR-CD3 relaxation. pMHC-induced reduced cohesion of TCR-CD3 was independent of ζ phosphorylation, as identical results were obtained after A770041 treatment (Fig. **6A** middle and right panels). Allosteric interaction typically occurs in the µs to few ms time-scale (Volkman et al., 2001), similar to the time required by pMHC binding to induce conformational changes in Cβ loops (Natarajan et al., 2017). Because pMHC binding dwell-times are of a much longer time-scale (e.g., hundreds of ms to min), allosterically-induced conformational changes should be observable at non-physiological lower temperatures. Consistently, almost identical reduction of ζ/β_2_ ratio was observed when (9V-A2)_4_ was reacted with cells at 0 °C (Fig. **6B** middle and right panels). To exclude that our observations were biased by the particular mutations introduced in βCDR2 and/or by the non-physiological affinity of wtc51, we used QM-α TCR, a 1G4 variant carrying mutations in αCDR2, βCDR2 and βCDR3 (Fig. **S6B**) (Irving et al., 2012), which confer a 140 nM *K*_d_ (*k*_off_, 0.015 sec^-1^ at 25°C) for NY-ESO-1_157-165_-HLA-A2 (Irving et al., 2012), which is within the physiological range of TCR-pMHC binding affinity (Aleksic et al., 2012; Cole et al., 2017; Cole et al., 2007; Stone et al., 2009). Molecular modelling showed that that QM-α and 1G4-WT have superimposable canonical orientation when bound to NY-ESO-1_157-165_-HLA-A2 (Fig. **S6B**). Figure **6C** showed that binding of (9V-A2)_4_-His to QM-α induced strong Erk activation (Fig. **6C**, left panel) and reduced ζ/β_2_ ratio (40 %), which remained unchanged after A770041 treatment (Fig. **S6C**). These observations were extended to 868, a TCR isolated from an HIV elite controller (Varela-Rohena et al., 2008). 868 recognises a spontaneously mutated HIV p17 Gag-derived peptide SLYNTIATL (6I) presented by HLA-A2, with a *K*_d_ of 53 nM at (*k*_off_, 0.0013 at 4°C (Cole et al., 2017). Being a natural TCR directed at a viral antigen, 868 was ideal to validate the data obtained with *in vitro*-modified TCRs against a tumour antigen. Binding of tetramerised ligand (6I-A2)_4_-His stimulated strong Erk activation (Fig. **6D**, left panel) and weakened 868 quaternary structure cohesion, as shown by the reduced ζ/β_2_ ratio (Fig. **6D**, middle and right panels). The occurrence of the same effect (i.e., structural changes) in three different TCRs by mere coincidence is highly unlikely but it is rather the consequence of the same cause: ligand-induced conformational changes that modify critical contacts maintaining TCR-CD3 complex cohesion (Alcover et al., 1990; Call et al., 2002; Dong et al., 2019). Reduced ζ cohesion was also observed in wtc51 when expressed in primary human T cells stimulated with (9V-A2)_4_-His (Fig. **6E**), excluding non-physiological behaviour of TCR-CD3 in the PM of Jurkat cells. To date, it remains unclear whether anti-CD3ε Abs used in clinical settings activate TCR-CD3 by mechanisms distinct from that of pMHC. To address this question, we slightly modified the DSA (STAR Methods). We employed mono-biotinylated Fab’ of UCHT1 anti-CD3ε as a proxy for minimally- or non-stimulated receptor and mono-biotinylated UCHT1 Ab to stimulate and capture TCR-CD3 with streptavidin for IB analysis. Since TCR-CD3 was captured via CD3ε, β/ ε and ζ/ ε ratios were used to assess TCR-CD3 cohesion. We found that UCHT1 Ab binding reduced β/ ε and ζ/ εratios, hence the cohesion of ε with β but less so with ζ (Fig. **6F**). Similar observations were made if cells were pre-treated with A770041 (Figs. **6G** and **S6D**) or reacted with UCHT1 at 0°C (Figs. **6H** and **S6E**). Taken together, these observations and the TMR mutants’ phenotype strongly suggested that TCR-CD3 signals intracellularly by an allosteric interaction propagating from the αβ binding site to the CD3 subunits and modifying critical contacts within the TMRs.

**Figure 6.**
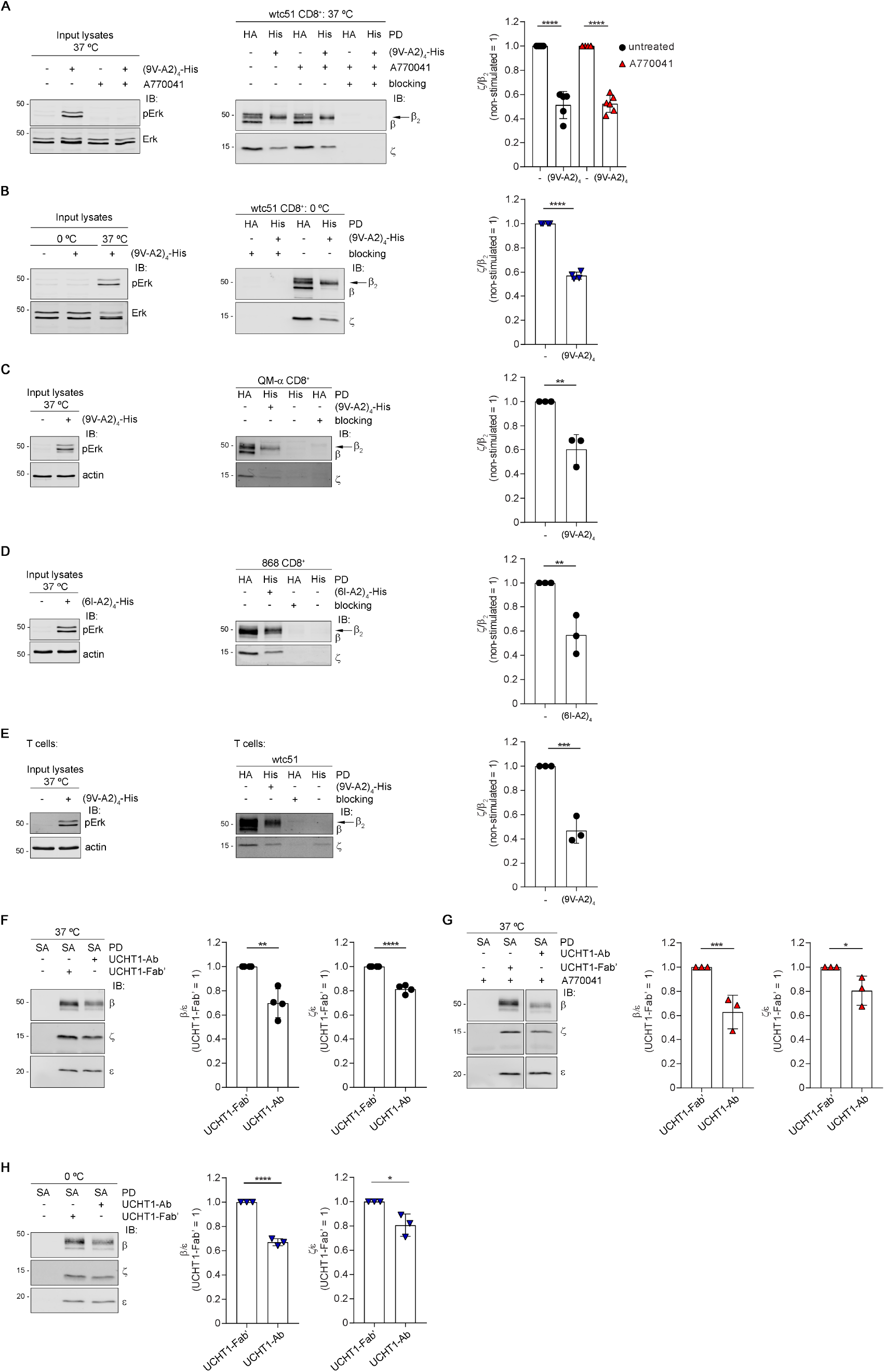
pMHC tetramer binding loosens αβ association with ζ. **A** J76 wtc51 stimulated with (9V-A2)_4_-His ±A770041. **Left**, pErk response (1 of 4 experiments). **Middle**, PD with anti-HA Ab (HA) (lanes 1, 3, 5) or His-Cobalt beads (His) (lanes 2, 4, 6) followed by IB for β and ζ. Blocking was with HA peptide or imidazole. 1 of 6 experiments. The arrow indicates β_2_ isoform. **Right**, 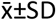 of ζ/β_2_, n=5-6, unpaired *t*-test *p*<0.0001. **B** J76 wtc51 stimulated with (9V-A2)_4_-His at 0°C. **Left**, pErk response (1 of 4 experiments). **Middle**, β-HA (lanes 1, 3) or His (lanes 2, 4) PD and IB for β and ζ (1 experiment of 4). The arrow indicates β_2_ isoform. **Right**, 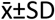 of ζ/β_2_, n=4, unpaired *t*-test *p*<0.0001. **C** J76 QM-α stimulated with (9V-A2)_4_-His. **Left**, pErk response (1 of 3 experiments). **Middle**, β-HA (lanes 1, 4) or His (lanes 2, 3) PD and IB for β and ζ (1 experiment of 3). The arrow indicates β_2_ isoform. **Right**, 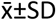 of ζ/β_2_, n=3, unpaired *t*-test *p*<0.01. **D** J76 868 stimulated with (6I-A2)_4_-His. **Left**, pErk response (1 of 3 experiments). **Middle**, β-HA (lanes 1, 3) or His (lanes 2, 4) PD and IB for β and ζ (1 experiment of 3). The arrow indicates β_2_ isoform. **Right**, 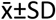 of ζ/β_2_, n=3, unpaired *t*-test *p*<0.01. **E** Primary T cells expressing wtc51 stimulated with (9V-A2)_4_-His. **Left**, pErk response (1 of 3 experiments). **Middle**, β-HA (lanes 1, 3) or His (lanes 2, 4) PD and IB for β and ζ (1 experiment of 3). The arrow indicates β_2_ isoform. **Right**, 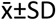 of ζ/β_2_, n=3, unpaired *t*-test *p*<0.001. **F** J76 1G4 ±UCHT1-Fab’ or UCHT1-Ab. **Left**, streptavidin (SA)-mediated PD and IB for β, ζ and ε (1 of 4 experiments). **Right**, 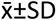 of β/ε and ζ/ε, n=4, unpaired *t*-test *p*=0.0023, *p*<0.0001. **G** J76 1G4 ±A770041 incubated or not with UCHT1-Fab’ or UCHT1-Ab. **Left**, SA-mediated PD and IB for β, ζ and ε (1 of 3 experiments). See also Fig. **S6D. Right**, 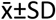 of β/ε and ζ/ε, n=3, unpaired *t*-test *p*=0.0007, *p*=0.0494. **H** J76 1G4 ±UCHT1-Fab’ or UCHT1-Ab at 0°C. **Left**, SA-mediated PD and IB for β, ζ and ε (1 of 3 experiments). **Right**, 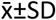 of β/ε and ζ/ε, n=3, unpaired *t*-test *p*<0.0001, *p*=0.0218. See also Fig. **S6E**.

### Monovalent pMHC in solution triggers TCR-CD3 untying and intracellular signalling

pMHC tetramers induced conformational change and signalling without applying force. However, pMHC tetramers necessarily induced fast TCR-CD3 clustering and therefore cannot allow to discern if receptor aggregation was responsible for allosteric activation, as previously suggested (Minguet et al., 2007). We therefore reacted wtc51-expressing J76 with biotinylated soluble monovalent (sm)-9V-A2 (Fig. **7A**). Mono-dispersion of (sm)-9V-A2 was controlled by size-exclusion chromatography-multi-angle-light scattering (SEC-MALS), just before use (Fig. **S7B**, note that fractions within the sm-9V-A2 peak were used). Following DDM solubilisation, sm-9V-A2-bound TCR-CD3 was captured by His-Streptavidin/His-Cobalt beads (Fig. **S7A**) and ζ recovery examined. The data showed that ζ/β_2_ ratio was considerably reduced in sm-9V-A2-bound vs. unbound wtc51 and was unaffected by A770041 treatment (Fig. **7A**, IBs and histograms) or by the absence of CD8 co-receptor (Fig. **7B**). To definitively exclude potential sm-pMHC cross-linking after solubilisation by streptavidin used for capturing ligand-bound TCR-CD3, we used instead an Avidin monomer (mAv). However, this condition did not change the result (Fig. **S7D**). Similar ζ/β_2_ reduction was observed for 868 TCR reacted with monodispersed sm-6I-A2 (Figs. **7C** for DSA and **S7C** for SEC). Figure **7E** (Ibs and histograms) shows that sm-9V-A2 reduced ζ recovery also in wtc51 expressed in primary T cells, excluding a bias of Jurkat cell PM. A more stringent test for TCR-CD3 allosteric regulation was to assess whether sm-pMHC promotes quaternary structure untying after solubilisation. Sm-9V-A2 bound to wtc51 TCR at 0 °C in post-nuclear lysates with 0.5 % DDM, as revealed by streptavidin IB (Fig. **S7E**), considerably reduced ζ/β_2_ ratio (Fig. **7F**). This indicated that TCR-CD3 complex loosening by pMHC binding did not require intact PM and therefore, as it should be expected for an allosteric change, it relied essentially on protein-protein interactions. Moreover, because it occurred in isolated TCR-CD3, these data further corroborated the idea that the allosteric change was independent of force, clustering and co-receptor. If the conformational change induced by sm-pMHC was functionally relevant, it should also induce intracellular signalling. Previous work could not demonstrate that binding of sm-pMHC in solution elicited [Ca^2+^]_i_increase unless co-receptor was expressed (Delon et al., 1998). However, we found that sm-9V-A2, controlled by SEC-MALS for being mono-dispersed (Fig. **S7B**), did induce robust pErk in both CD8-efficient (Fig. **7A**) and CD8-deficient (Fig. **7B**) J76 cells expressing wtc51, that was abolished by A770041 (Fig. **7A**). Erk activation by sm-9V-A2 was dose-dependent (Fig. **S7F**), with as little as 3 nM inducing 50 % of the maximum and occurred at 2 min after sm-9V-A2 addition (Fig. **S7G**), similar to (9V-A2)_4_ stimulation of 1G4-WT (Paster et al., 2015), though (9V-A2)_4_ generally induced stronger pErk response. We obtained similar data with 868 in presence or absence of CD8 co-receptor (Figs. **7C, 7D** and **S7C**) and with QM-αwithout CD8 (Fig. **S7H**). Non-specific adsorption of sm-pMHC onto J76 cell membrane during the stimulation assay was negligible even at the highest sm-9V-A2 concentration (Fig. **S7I**). This made unlikely that signalling by sm-9V-A2 was the consequence of non-specific adsorption to the cells surface resulting in cell-to-cell ligand cross-presentation rather than direct stimulation by soluble sm-9V-A2. Moreover, we experimentally tested whether even this negligible amount of non-specifically bound sm-9V-A2 on J76 cells could be stimulatory. However, we did not detect any Erk activation (Fig. **S7J**). Multiple reasons can explain why our data apparently contradict previous observations. First and foremost, we used TCRs of reduced *k*_off_ (higher-affinity range) for pMHC, including a natural one (868). sm-pMHC ligands of low-medium affinity range (µM) can be expected to induce low/non-sustained [Ca^2+^]_i_ increase, whose ramp-up requires a more robust and complex cascade of additional events (Irvine et al., 2002; Lewis, 2019), including co-receptor implication (Delon et al., 1998; Minguet et al., 2007). Also, sm-pMHC engages TCR-CD3 without immediately clustering it, contrary to pMHC tetramers that provide this critical signalling-reinforcing effect (see Discussion). Moreover, membrane-tethered pMHC has lower degree of freedom than soluble pMHC, a property that sensibly increases pMHC on-rate (Huppa et al., 2010; O’Donoghue et al., 2013). Comprehensively, our genetic, biochemical, MDS and functional data constitute substantial evidence that TCR-CD3 is a genuine allosteric device. We name this model “TCR-CD3 allosteric relaxation” (Fig. **S7K**) as a mechanism sufficient to incite initial T cell activation solely by pMHC binding.

**Figure 7.**
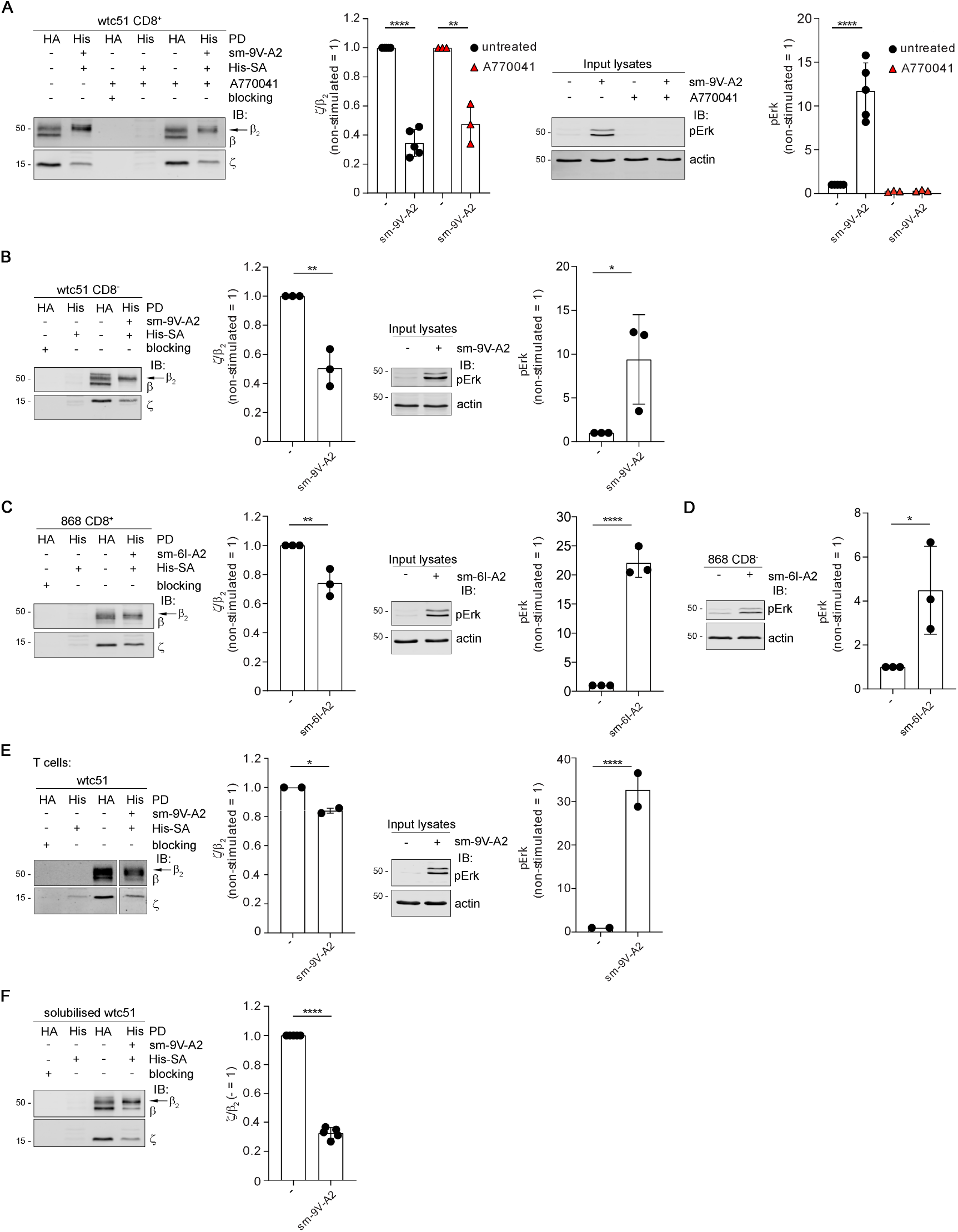
Monovalent pMHC in solution triggers TCR-CD3 untying and intracellular signalling. **A** J76 wtc51 ±A770041 stimulated or not with sm-9V-A2 were lysed and subjected to PD with anti-HA Ab or Talon beads. **First panel**, anti-HA (β-HA) (lanes 1, 3, 5) or Talon beads (His) (lanes 2, 4, 6) PD and IB for β and ζ (1 experiment of 3). The arrow indicates β_2_ isoform. **Second panel**, 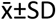 of ζ/β_2_, n≥3, unpaired *t*-test *p*<0.0001 and *p*<0.01. **Third panel**, pErk IB: 1 of 3 experiments. **Fourth panel**, 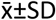 of pErk, n≥3, unpaired *t*-test *p*<0.0001. **B** CD8-deficient J76 wtc51 stimulated or not with sm-9V-A2 were processes as in **A. First panel**, β-HA (lanes 1, 3) or His (lanes 2, 4) PD and IB for β and ζ (1 of 3 experiments). The arrow indicates β_2_ isoform. **Second panel**, 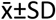 of ζ/β_2_, n=3, unpaired *t*-test *p*<0.01. **Third panel**, pErk IB (1 of 3 experiments). **Fourth panel**, 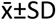 of pErk, n=3, unpaired *t*-test *p*<0.05. **C** J76 868 stimulated or not with sm-6I-A2 were processes as in **A. First panel**, β-HA (lanes 1, 3) or His (lanes 2, 4) PD and IB for β and ζ (1 of 3 experiments). The arrow indicates β_2_ isoform. **Second panel**, 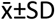 of ζ/β_2_, n=3, unpaired *t*-test *p*<0.01. **Third panel**, pErk IB: 1 of 3 experiments. **Fourth panel**, 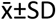 of pErk, n=3, unpaired *t*-test *p*<0.0001. **D** CD8-deficient J76 868 stimulated or not with sm-6I-A2. **Left**, pErk IB (1 of 3 experiments). **Right**, 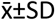 of pErk, n=3, unpaired *t*-test *p*<0.05. **E** Primary T cells expressing wtc51 stimulated or not with sm-9V-A2 were processes as in **A. First panel**, β-HA (lanes 1, 3) or His (lanes 2, 4) PD and IB for β and ζ (1 of 2 experiments). The arrow indicates β_2_ isoform. **Second panel**, 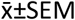 of ζ/β_2_, n=2, unpaired *t*-test *p*<0.05. **Third panel**, pErk IB: 1 of 2 experiments. **Fourth panel**, 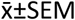 of pErk, n=2, unpaired *t*-test *p*<0.0001. **F** J76 wtc51 were lysed, incubated or not with sm-9V-A2 and subjected to PD by anti-HA or Talon beads. **Left**, β-HA (lanes 1, 3) or His (lanes 2, 4) PD and IB for β and ζ (1 of 5 experiments). The arrow indicates β_2_ isoform. **Right**, 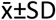 of ζ/β_2_, n=5, unpaired *t*-test *p*<0.0001.

## Discussion

Allostery governs signal transduction of many membrane receptors (Changeux and Christopoulos, 2016). However, this hard-wired, tuneable and fast interaction mode exploiting protein conformational flexibility has not gained sufficient traction for the elucidation of TCR-CD3 signalling mechanism (Mariuzza et al., 2020). To gather insight into TCR-CD3 signalling mode, we used a genetic perturbation approach and uncovered a previously unnoticed allosteric property of the entire TCR-CD3 complex. We found that TMR mutations that loosened cohesion between TCRαβ and CD3ζ populated TCR-CD3 activated state and increased agonist’s potency. This gain-of-function phenotype mimicked pMHC agonist binding that also reduced cohesion between TCRαβ and CD3ζ, independently of CD3 ITAM phosphorylation and at 0 °C. These convergent results suggested that weakening of TCR-CD3 TMR contacts is a key step in an allosteric mechanism initiated by pMHC binding and culminating in ITAM phosphorylation. We favour the idea that conformational changes occurring at the pMHC binding site propagate to CαCβ ECDs, where they contact the CD3 subunits as indicated by several investigations (Beddoe et al., 2009; He et al., 2020; Natarajan et al., 2017; Rangarajan et al., 2018). The ECDs and TMRs of TCRαβ, CD3δε and CD3γε show extended intra-dimer interface, yet less so between dimers (Dong et al., 2019), suggesting that each dimer retains flexibility vis-à-vis the other dimers. Moreover, CD3δε and CD3γε interact much more with TCRαβ than with each other. It is therefore conceivable that pMHC binding can induce reshuffling of contacts between ECD dimers, making the CD3δε and CD3γε acquiring a higher degree of freedom vis-à-vis TCRαβ. This may result in slight rotation and/or translation of CD3δε and/or CD3γε vis-à-vis TCRαβ. The mechanical rigidity conferred to the CPs of ε, δ and γ by the Cys-Cys loop (Alcover et al., 2018) could transmit these movements to the respective TMRs, resulting in local rearrangements of helix-helix packing, similar to those caused by the activating TMR mutations studied here and perhaps of interfacial lipids (Gupta et al., 2017) (Fig. **S7K**). Consistently, mutations of ε Cys-Cys loop affect TCR-CD3 signalling (Wang et al., 2009). The relaxed quaternary structure of ligand-activated TCR-CD3 could reduce contacts between the TMRs of αβ with ζζ (the most loosely attached dimer), making ζζ prone to detach from the rest of the complex, due to further erosion of TMR contacts by DDM during membrane extraction. As our data suggest, TMR quaternary structure relaxation activated by pMHC (or anti-CD3 Ab) or by TMR mutations promotes ITAM accessibility by active-Lck (Nika et al., 2010), which would require conformational changes of membrane-tethered CD3 intracellular tails (Xu et al., 2008). However, the membrane-juxtaposed segments of all CD3 subunits are intrinsically disordered, hence they may lack mechanical rigidity required to respond to TMR movements. We suggest instead that subtle untying of TMRs may indirectly reduce the grip of CD3 intracellular tails to the membrane and favour ITAM tyrosine exposure (Fig. **S7K**). Phosphatidylserine (PS) (Xu et al., 2008) and PIP2 (Chouaki-Benmansour et al., 2018) are thought to keep the CD3 tails retracted onto the PM. An attractive possibility is that local TMR octamer rearrangement permits fast exchange of PIP2 and PS with neutral lipids that may reduce CD3ζ and ε tails interaction with the lipid bilayer (Fig. **S7K**), gradually augmenting the exposure of ITAM tyrosines to active-Lck. Changes in cholesterol interacting with TMR helices (Swamy et al., 2016; Wang et al., 2016) and/or with the tyrosines of the ITAMs might be part of this mechanism. Agonist anti-CD3ε mAb produced similar gain-of-function in TMR mutants as well as quaternary structure untying, in agreement with CD3ε ECD lying on the conformational trajectory activated by pMHC binding.

A key finding of our investigation is the formal evidence that binding of soluble monovalent and mono-dispersed pMHC (sm-pMHC) alone to membrane-bound or detergent-solubilised TCR-CD3 suffices to induce TCR-CD3 quaternary structure relaxation and signal transduction. Stimulation of TCR-CD3 by sm-pMHC alone agrees with a genuine allosteric mechanism, as hinted by our genetic perturbation analysis, much like membrane receptors activated by soluble ligands. Allosteric activation occurred without co-receptor or TCR-CD3 clustering or force, making extrinsic energy source, such as actomyosin-induced membrane movements required for mechanosensing (Das et al., 2015; Kim et al., 2009; Liu et al., 2014) dispensable for igniting TCR-CD3 signalling. A *K*_d_ of 7 µM was insufficient to analyse biochemically quaternary structure cohesion of 9V-A2-bound vs. free 1G4 TCR. However, we succeeded by using 1G4 variants wtc51 and QM-α capable of binding 9V-A2 with a *K*_d_ of 15 nM and 140 nM, respectively, the latter within the physiological *K*_d_ range for pMHC agonists (Aleksic et al., 2012; Cole et al., 2007; Stone et al., 2009). A third example was 868, a CTL-derived anti-HIV TCR, whose *K*_d_ was 53 nM, slightly higher than the lower limits of ≈ 100 nM observed for naturally occurring pMHC-TCRs binding. This was due to a single natural mutation in the antigen peptide (6V → 6I) that raised the *K*_d_ towards the unmutated epitope of 170 nM (Aleksic et al., 2012; Cole et al., 2017). All these TCRs showed ligand-induced quaternary structure loosening and intracellular signalling upon pMHC engagement. There is no evidence for high range affinity TCRs to induce T cell signalling and activation different from ligands of one-two-digit µM affinities and proven valid for T-cell adoptive immunotherapy. Thus, it is unlikely that higher-than-normal affinity for pMHC confers to TCR-CD3 an allosterically regulated signalling while lower affinities do not. Indeed, allosteric changes in TCRαβ have been also demonstrated with pMHC binding with µM *K*_d_ (Beddoe et al., 2009; He et al., 2020; Natarajan et al., 2017; Rangarajan et al., 2018).

Early work could not detect [Ca^2+^]_i_ increase with sm-pMHC, unless CD8 was co-engaged (Delon et al., 1998). Others also did not observe TCR-CD3 signalling using pMHC monomer (Boniface et al., 1998). This apparent conflict with our data can be reconciled by considering differences in sensitivity of the signalling outputs measured (i.e., [Ca^2+^]_i_ vs. pErk) and increased ligand dwell-time in our experiments (in absence of co-receptor). Moreover, membrane-tethered pMHC considerably increases *k*_on_ (and little or no increase of the *k*_off_) as compared to in solution measures (Huppa et al., 2010; O’Donoghue et al., 2013), suggesting more effective entropically-driven signalling by the former. Moreover, [Ca^2+^]_i_ rise of high amplitude and duration requires sustained TCRαβ engagement (Irvine et al., 2002; Lewis, 2019), likely achieved by higher lateral ordering of cell-surface and signalling complexes in micro-clusters and immunological synapse (IS) (Varma et al., 2006). The combination of these conditions (e.g., co-receptors and clustering), that sets in motion robust intracellular signalling mechanism (i.e. sustained signalling for full T-cell activation), may not be required for just igniting TCR-CD3 signal transduction as sm-pMHC alone does.

Mechanical forces play multiple roles in T cell activation at the molecular and cellular levels (Zhu et al., 2019). Reduced *k*_off_ of TCR-pMHC interaction is observed when subjected to ≈ 10 -20 pN pulling force, which means that “catch-bond” can be formed (Liu et al., 2014). However, mechanical force (pulling/pushing) generated by membrane fluctuations and/or dedicated actin protrusions are observed in the time-scale of seconds, much slower (at least three order of magnitude) than allosteric changes propagating occurring in µs to ms (Kern and Zuiderweg, 2003; Natarajan et al., 2017; Rangarajan et al., 2018). Thus, pMHC-induced signal transduction and initial intracellular signalling could occur without the need of active mechanical force. It is also interesting that recent biophysical data suggest that force developed between membrane-tethered TCR-CD3 and pMHC is of fairly low magnitude (∼ 2 pN)(Göhring et al., 2020). Perhaps, this low-amplitude force may play a role in extending the purely allosterically-induced interactions supported by our study.

Changes in conformational dynamics can have long-range consequences of functional relevance, a mechanism known as dynamic allostery (Tzeng and Kalodimos, 2012), which relies on changes in conformational entropy only. Conformational entropy cannot be frequently observed in a protein’s crystalline state, unlikely to capture a protein higher-energy (activated) state. However, NMR can correlate very fast local conformational changes (ps, ns) occurring at distant sites over time-scales of µs to ms, compatible with allosteric regulation (Kern and Zuiderweg, 2003; Natarajan et al., 2017). Dynamic allostery may therefore apply to TCRαβ, as pMHC binding induces changes in conformational dynamics at distal H3, H4 helices and FG loop of Cβ and Cα AB loop (Beddoe et al., 2009; He et al., 2020; Natarajan et al., 2017; Rangarajan et al., 2018), the latter having been captured only in a single crystal structure (Kjer-Nielsen et al., 2003) of many solved to date (http://atlas.wenglab.org/web/index.php. High affinity TCR-CD3-pMHC interacting by non-canonical orientation cannot induce signalling (Adams et al., 2011). In the light of our data, allosteric activation of TCR-CD3 occurs perhaps optimally only with ligands interacting in a diagonal canonical orientation.

Our data should help reconcile controversies about TCR-CD3 signalling mechanism. Thus, pMHC co-engagement by TCR and co-receptor has been found to be conditional on initial TCR-CD3 signalling (Casas et al., 2014; Jiang et al., 2011) and catch-bonding was contrasted by inhibiting Lck (Hong et al., 2018). We found that enhanced basal signalling by 1G4-βA291 induced weak but detectable clustering that was erased by Lck inhibition. These data suggest that all these events are instigated by an initial mechanism of allosteric nature induced solely by pMHC binding. Thus, co-receptor engagement, receptor clustering, shielding from PTPs and actomyosin-driven mechanical force (facilitating clustering and catch-bonding) may stabilise and potentiate initial allosterically-induced signalling. They may help reduce physical and chemical noise during receptor signalling, augmenting and stabilising signals of narrow amplitude and duration initiated by sparse engagement of pMHC monomers (Brameshuber et al., 2018).

The fast time-scale by which allosteric interaction propagates should ensure that ITAMs’ exposure to active-Lck relies on pMHC binding dwell-time compatible with both very weak (self) and agonist ligands (e.g., hundreds of ms to sec). However, allosteric activation for receptors that recognise multiple ligands raises the possibility of “biased agonism” (Freed et al., 2017; Furness et al., 2016; Lane et al., 2017) whereby different ligand-induced conformational changes and/or ligand-binding kinetics correlate with distinct functional outputs. In principle, allosteric interaction activated by different TCRαβ CDR loop conformational changes upon canonical orientations over pMHC might follow different conformational trajectories propagating along TCR-CD3. In this case, ligand potency may result from both binding kinetics and variable conformational trajectories. Alternatively, binding of diverse pMHC-TCRαβ interfaces in canonical orientation may generate equivalent allosteric changes making ligand binding kinetics the unique factor governing ligand potency. Future studies on TCR-CD3 using native nanodiscs, cryo-EM, MDS and genetic perturbation should help to further clarify these questions. We anticipate that the novel data for TCR-CD3 signalling reported here should spark interest for innovative strategies to harness TCR-CD3 signalling for immunotherapy.

## Methods

Please refer to Supplementary Information for a detailed description of all experimental procedures used in this investigation.

## Acknowledgements

Financial support: Wellcome Trust WT200844/Z/16/Z to O.A.; WT092970MA and WT208361/Z/17/Z to M.S.P.S.; WT100262Z/12/Z to M.L.D.; 207537/Z/17/Z to O.D.; SBF002\1031 to A.C.K. Springboard Award and WT. ERC-2014-AdG 670930 to S.B. Care-for-Rare Postgraduate Fellowship to A-L.L. For MDS, High-Performance Computing facilities, University of Leeds. We thank Juliane Cohen, Christine Ralf and Salvatore Valvo for technical support and John Orban, Brian Baker, Ian Wilson, Gerhard Schütz, Pedro Carvalho, Marco Fritzsche, Hai-Tao He, Immanuel Luescher, Cheng Zhu, Andrew Sewell for discussions, suggestions and for reading the manuscript; C.R. and J.C. and Ana Maria Vallés for manuscript editing. We apologise to colleagues whose work could not be adequately cited and commented herein because of space limitation.

## Author contributions

Conceptualization, O.A.

Methodology, A-L.L.; G.M.; N.P.; A.C.; D.C.; R.R.; D.K.C.; M.L.; S.B.; A.C.K.; D.P.; O.D.; M.L.D.; M.S.P.S.; O.A.

Software, A.C.K.; O.D.

Formal Data Analysis, A-L.L.; G.M.; N.P.; A.C.; D.C.; D.P.; S.B.; O.D.; M.S.P.S.; O.A.

Investigation, A-L.L.; G.M.; N.P.; A.C.; D.C.; A.C.K.; S.B.; D.G.; D.K.C.; D.P.; M.S.P.S.; O.A.

Resources, S.B.

Writing - Original Draft, O.A.

Writing - Review and Editing, O.A.; A-L.L.; G.M.; N.P.; A.C.; D.P.; A.C.K.; M.S.P.S.; D.K.C.; M.L.;

M.L.D. and D.C.

Visualization, G.M.; A-L.L.; N.P.; A.C.K.; A.C.; D.C.; D.P.; D.K.C.; S.B.; O.A. Supervision, A.C.K.; M.S.P.S.; M.L.D.; O.A.

Funding acquisition, A.C.K.; M.S.P.S.; M.L.D.; O.A.

