## Supplementary Figures and Methods for "Allosteric activation of T-cell antigen receptor signalling by quaternary structure relaxation": Supplementary Figures and Methods.pdf

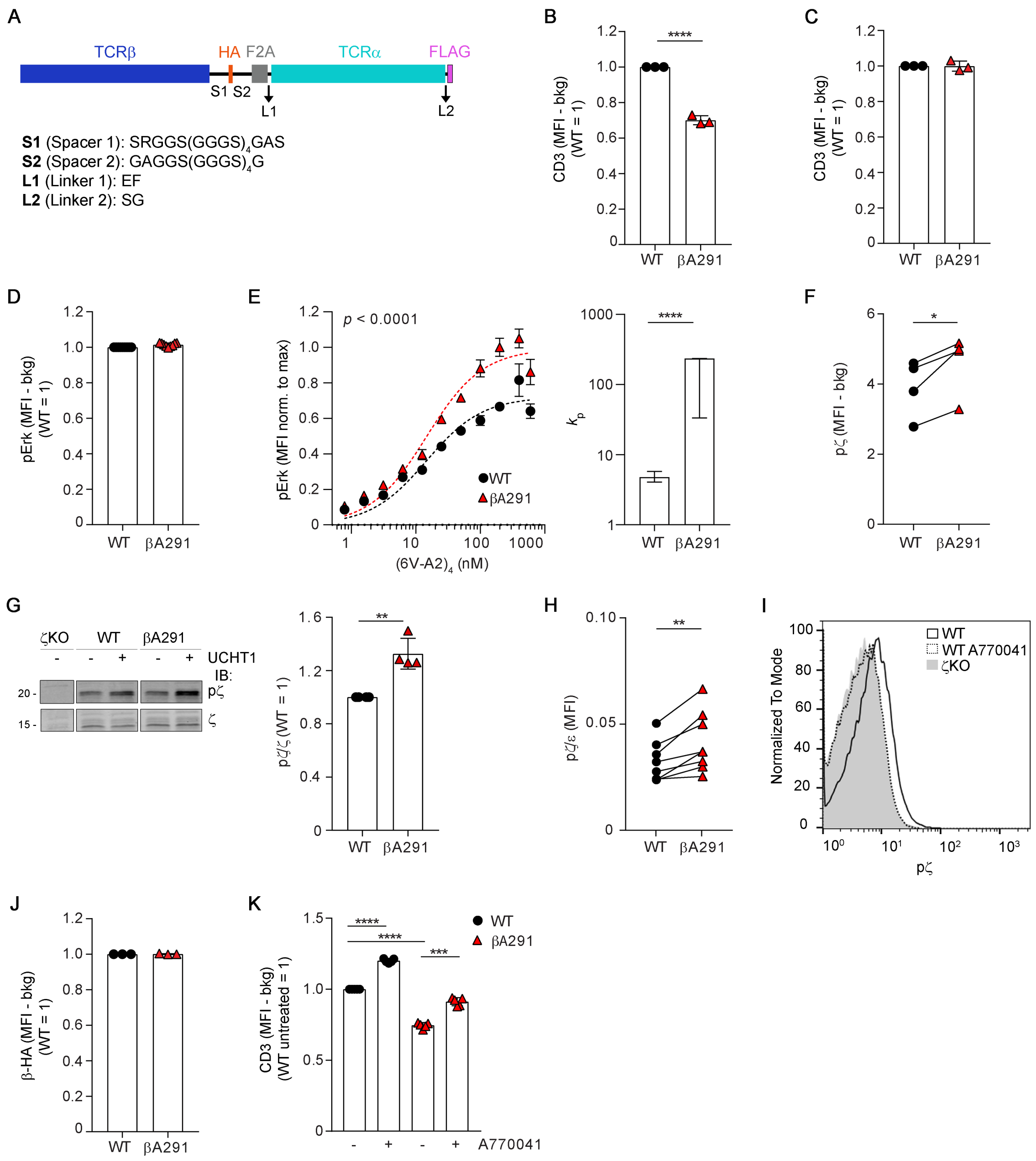

**Figure S1. Gain-of-function mutations in the βTMR**

**A** Scheme of the single polypeptide self-splicing 1G4 αβ expressed in J76 and CD8-deficient J76 cells. β chain (blue), α chain (cyan), HA-tag (orange), FLAG-tag (magenta), F2A sequence (grey). The entire aa sequence can be found in **Table 1** of STAR Methods. Spacer and short linker aa sequences are indicated below the scheme. **B** CD3 surface expression of CD8-deficient J76 1G4-WT or 1G4-βA291. Cells were labelled or not with CellTrace violet, mixed 1:1 and analysed for CD3 surface expression and total TCRβ-HA by FACS. Gates for equal HA-tag expression level were applied and the CD3 MFI of 1G4-WT and 1G4-βA291 was extracted from the TCRβ-HA$^{low}$ gate, see STAR Methods. Mean ± SEM of background subtracted CD3 MFI from the indicated gate normalised to the corresponding WT in the same gate, n = 3 experiments in duplicates, unpaired *t*-test $p < 0.0001$. **C** Similar CD3 surface expression of CD8-deficient J76 1G4-WT and 1G4-βA291. Cells were induced with different doses of doxycycline, labelled or not with CellTrace violet, mixed 1:1 and analysed for CD3 surface expression by FACS, see STAR Methods. Mean ± SD of $CD3^+$ MFI normalised to WT, n = 3 experiments in triplicates, unpaired *t*-test (ns). **D** PMA/Iono induced pErk response of CD8-deficient J76 1G4-WT or 1G4-βA291. Cells were labelled or not with CellTrace violet, mixed 1:1, treated with PMA/Iono (5') and analysed for pErk by FACS. Mean ± SD of pErk MFI normalised to WT, n = 3 experiments in triplicates, unpaired *t*-test (ns). **E** pErk response of CD8-deficient J76 1G4-WT or 1G4-βA291 stimulated with $(6V\text{-}A2)_4$. Cells were induced with different doses of doxycycline, labelled or not with CellTrace violet, mixed 1:1 and stimulated for 3 min with different doses (0.78 – 600 nM) of PE-conjugated $(6V\text{-}A2)_4$ and analysed for pErk and $(6V\text{-}A2)_4$ binding by FACS. **Left**, $(6V\text{-}A2)_4$ nM vs. pErk MFI normalised to max (showed in Fig. **1C**) fitted to a minimal model of kinetic proofreading, n = 3 experiments in triplicates, $R^2$ = 0.84 (WT), 0.90 (βA291), F-test $p < 0.0001$. **Right**, mean ± SD of proofreading rate ($k_p$), n = 3 experiments in triplicates, F-test $p < 0.0001$. **F** Paired max. pζ values related to dose response reported in Fig. **1E**. J76 1G4-WT or 1G4-βA291 were labelled or not with CellTrace violet, mixed 1:1, stimulated for 1 min with increasing doses (3.125 - 200 nM) of PE-conjugated $(6V\text{-}A2)_4$ and analysed for pζ and $(6V\text{-}A2)_4$ binding by FACS. Background subtracted MFI of $(6V\text{-}A2)_4$ was plotted vs. pζ MFI and fitted by non-linear regression, n = 4, ratio paired *t*-test $p = 0.023$. **G** pζ response of J76 1G4-WT or 1G4-βA291 stimulated or not with UCHT1 Ab. **Left**, immunoblot (IB) representative of 4 experiments. **Right**, mean ± SD of pζ/ζ, n = 4, unpaired *t*-test $p = 0.0013$. **H** Paired mean values of the basal pζ in J76 1G4-WT or 1G4-βA291 reported in Fig. **1F**. Resting cells were analysed for pζ or CD3 surface expression by FACS. pζ MFI was normalised to surface CD3 MFI, n = 8, paired *t*-test $p = 0.0037$. **I** Representative FACS histograms of pζ staining related to Fig **1F**. J76 1G4-WT cells were treated or not with A770041 (5 μM) for 15 min at 37 $^oC$ and analysed for pζ by FACS. J76-1G4WT-ζKO served as a negative control. **J** β-HA total expression of CD8-deficient J76 1G4-WT or 1G4-βA291. Cells were labelled or not with CellTrace violet, mixed 1:1 and analysed for CD3 surface expression and total TCRβ-HA by FACS in a single HA bin, see STAR Methods. Mean ± SEM of background subtracted HA MFI in $HA^{low}$ gate, n = 3, unpaired *t*-test (ns). **K** TCR-CD3 surface expression in J76 1G4-WT or 1G4-βA291 treated or not with A770041, n = 5, unpaired *t*-test $p < 0.0001$ (βA291 vs. WT), $p < 0.0001$ (WT ± A770041), $p = 0.0003$ (βA291 ± A770041).

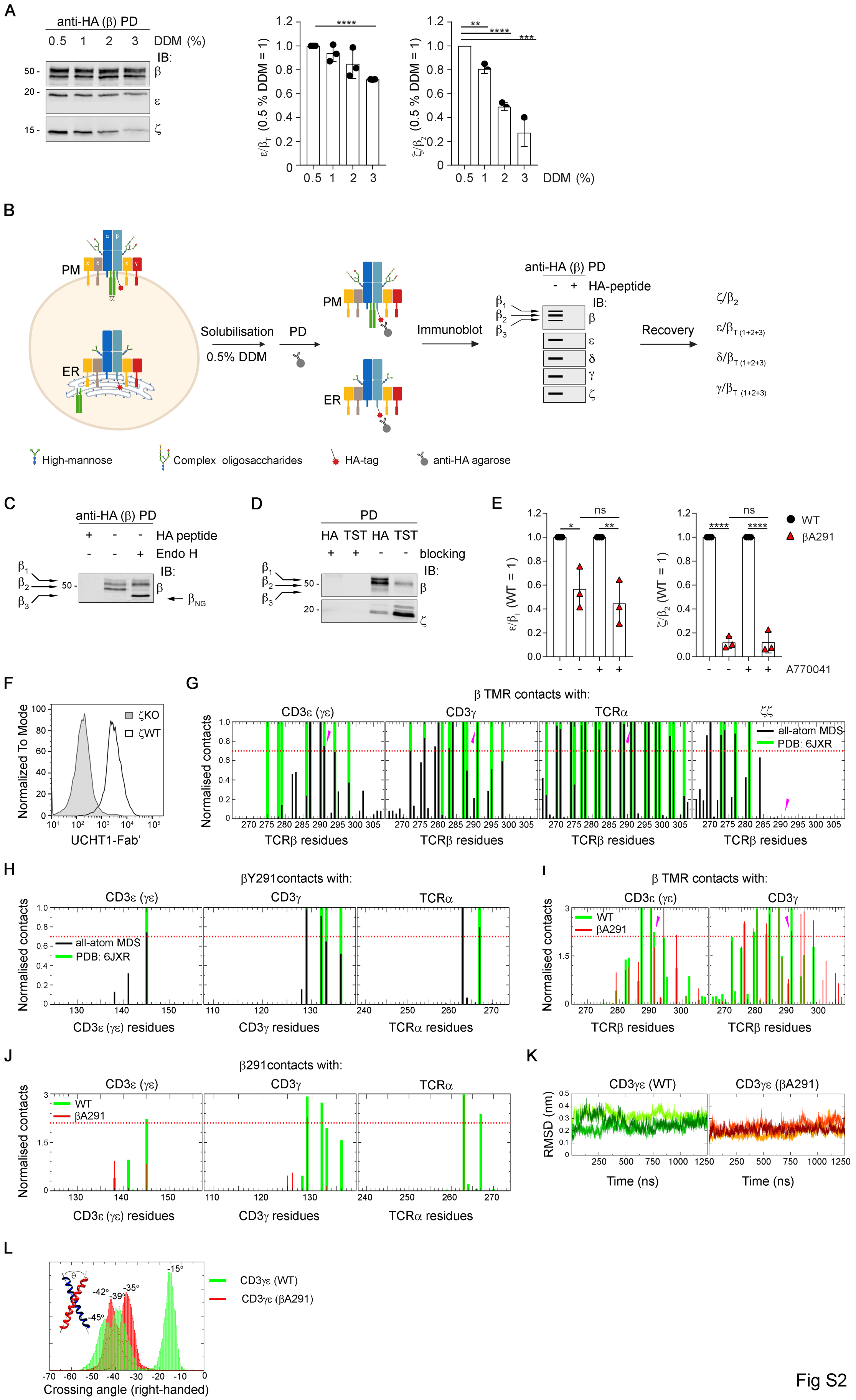
A
anti-HA (β) PD
0.5 1 2 3 DDM (%)
IB:
β
ε
ζ
ε/β_T (0.5 % DDM = 1)
ζ/β_2 (0.5 % DDM = 1)
DDM (%)
B
PM
ER
Solubilisation
0.5% DDM
PD
Immunoblot
anti-HA (β) PD
HA-peptide
Recovery
ζ/β_2
ε/β_T (1+2+3)
δ/β_T (1+2+3)
γ/β_T (1+2+3)
High-mannose
Complex oligosaccharides
HA-tag
anti-HA agarose
C
anti-HA (β) PD
HA peptide
Endo H
β_NG
D
PD
HA TST HA TST
blocking
E
WT
βA291
ε/β_T (WT = 1)
ζ/β_2 (WT = 1)
A770041
F
Normalized To Mode
ζKO
ζWT
UCHT1-Fab'
G
β TMR contacts with:
CD3ε (γε)
CD3γ
TCRα
ζζ
all-atom MDS
PDB: 6JXR
Normalised contacts
TCRβ residues
H
βY291contacts with:
CD3ε (γε) residues
CD3γ residues
TCRα residues
I
β TMR contacts with:
WT
βA291
J
β291contacts with:
K
CD3γε (WT)
CD3γε (βA291)
RMSD (nm)
Time (ns)
L
CD3γε (WT)
CD3γε (βA291)
Crossing angle (right-handed)
Fig S2

**Figure S2. βY291 contribution to TCR-CD3 quaternary structure cohesion**

**A** Sensitivity of TCR-CD3 quaternary structure cohesion to DDM concentrations. CD8-deficient J76 1G4-WT were solubilised with increasing concentrations of DDM and analysed by β-HA PD and IB for β, ε and ζ (**left**). Mean ± SD of $\varepsilon/\beta_T$ (**middle**) and $\zeta/\beta_2$ (**right**), n = 3, unpaired *t*-test ($\varepsilon/\beta_T$): $p < 0.0001$; ($\zeta/\beta_2$): 0.5 vs. 1 % $p < 0.01$, 0.5 vs. 2 % $p < 0.0001$, 0.5 vs. 3 % $p < 0.001$. Note that at ≤ 0.5 % DDM there is no detectable change in cohesion of CD3 subunits with TCRαβ, further supporting that in TCR-CD3 extracted at 0.5 % DDM the subunits' stoichiometry in the octamer remains intact (Swamy et al., 2008). At ≥ 1 % DDM there is progressive loss of ζ followed by γε and δε. **B** Graphic scheme describing the DSA. The entire pool of the TCRαβ is captured by anti-HA (β-HA) PD which includes partial αβγεδε complex assembled in the endoplasmic reticulum (ER) (Alcover et al., 2018), αβγεδεζζ complex resident in the *trans*-Golgi (Alcover et al., 2018) or recycling at steady state in the ER (Alcover et al., 2018) and the largest fraction of αβγεδεζζ present at the plasma membrane (PM). The IB scheme on the right shows expected band patterns for β, ε, δ, γ and ζ derived from the ER and PM. Arrows indicate the different isoforms of TCRβ ($\beta_1$, $\beta_2$, $\beta_3$). To evaluate ζ recovery, $\zeta/\beta_2$ ratio was calculated and the value for $\zeta/\beta_2$ ratio from WT was set equal to one. To evaluate ε, δ, γ recovery, $\varepsilon/\beta_T$, $\delta/\beta_T$ and $\gamma/\beta_T$ ratios were calculated and the ratios for WT were set equal to one. These represented the recovery of intact TCR-CD3 complex. Ratios < 1 indicate a lower recovery of ζ, ε, δ and γ hence a reduced cohesion of TCR-CD3 quaternary structure. See STAR Methods for a detailed description of the experimental procedure. **C** J76 wtc51 treated or not with endo H and analysed by β-HA PD and IB for β. Arrows indicate β isoforms, $\beta_{NG}$ indicates non-glycosylated β isoform after removal of high-mannose carbohydrates by endo-H treatment. Data representative of 2 experiments. **D** J76-1G4-ζKO expressing inducible ζTST were solubilised and analysed by β-HA (lanes 1, 3) or ζTST (lanes 2, 4) PD and IB for β and ζ. Arrows indicate β isoforms. IB representative of 2 experiments. **E** CD8-deficient J76 1G4-WT or 1G4-βA291 ± A770041 were solubilised and analysed by β-HA PD and IB for β, ε and ζ. Mean ± SD of $\varepsilon/\beta_T$ (**left**) and $\zeta/\beta_2$ (**right**), n = 3, unpaired *t*-test ($\varepsilon/\beta_T$): 1G4-WT vs. 1G4-βA291 $p < 0.05$, 1G4-WT+A770041 vs. 1G4-βA291+A770041 $p < 0.01$; ($\zeta/\beta_2$): 1G4-WT vs. 1G4-βA291 $p < 0.0001$, 1G4-WT+A770041 vs. 1G4-βA291+A770041 $p < 0.0001$; ns = non-significant. **F** Representative FACS histograms of UCHT1-Fab' staining of J76-1G4WT-ζKO reconstituted with doxycycline inducible ζWT. Cells were induced or not for ζWT expression, labelled or not with CellTrace violet, mixed 1:1 and analysed for TCR-CD3 surface expression by FACS. **G** Normalised number of contacts of the WT β TMR with the rest of the TCR-CD3 TMRs in our all-atom molecular dynamics simulations (MDS) (related to Fig. **2D**). The contacts in the cryo-EM structure (PDB: 6JXR) are shown in green for comparison with the WT all-atom MDS (black). Magenta arrow indicates β291. Normalisation was done by dividing the number of contacts of each residue by the highest number of contacts. For all contacts analyses in

Figs. **S2G** - **S2J**, a cut-off distance of 4 Å was used to define a contact and the red dotted line represents 70 % of the normalised contacts, a threshold used to measure the significance of contacts. **H** Normalised number of contacts of βY291 with the rest of the TCR-CD3 TMRs (related to Fig. **2D**). Comparison of protein-protein interactions between our WT all-atom MDS (black) and the cryo-EM structure (PDB: 6JXR) (green). Normalisation was done by dividing the number of contacts of each residue by the highest number of contacts. **I** Normalised number of contacts of βWT (green) and βA291 (red) with CD3γε TMR in our all-atom MDS. Magenta arrow indicates β291. Normalisation was done by dividing the number of contacts of each residue by the number of simulation frames. **J** Normalised number of contacts of βY291 WT (green) and βA291 mutant (red) in our all-atom MDS. Normalisation was done by dividing the number of contacts of each residue by the number of simulation frames. **K** Root mean square deviations (RMSD) of the Cα atoms of CD3γε relative to their initial configuration during the WT and βA291 all-atom MDS. Each line represents RMSD obtained from one simulation (n = 3). **L** Comparison of crossing angle distribution of CD3γε TMR between WT and βA291, in 3 simulations. The most observed crossing angle values are labelled.

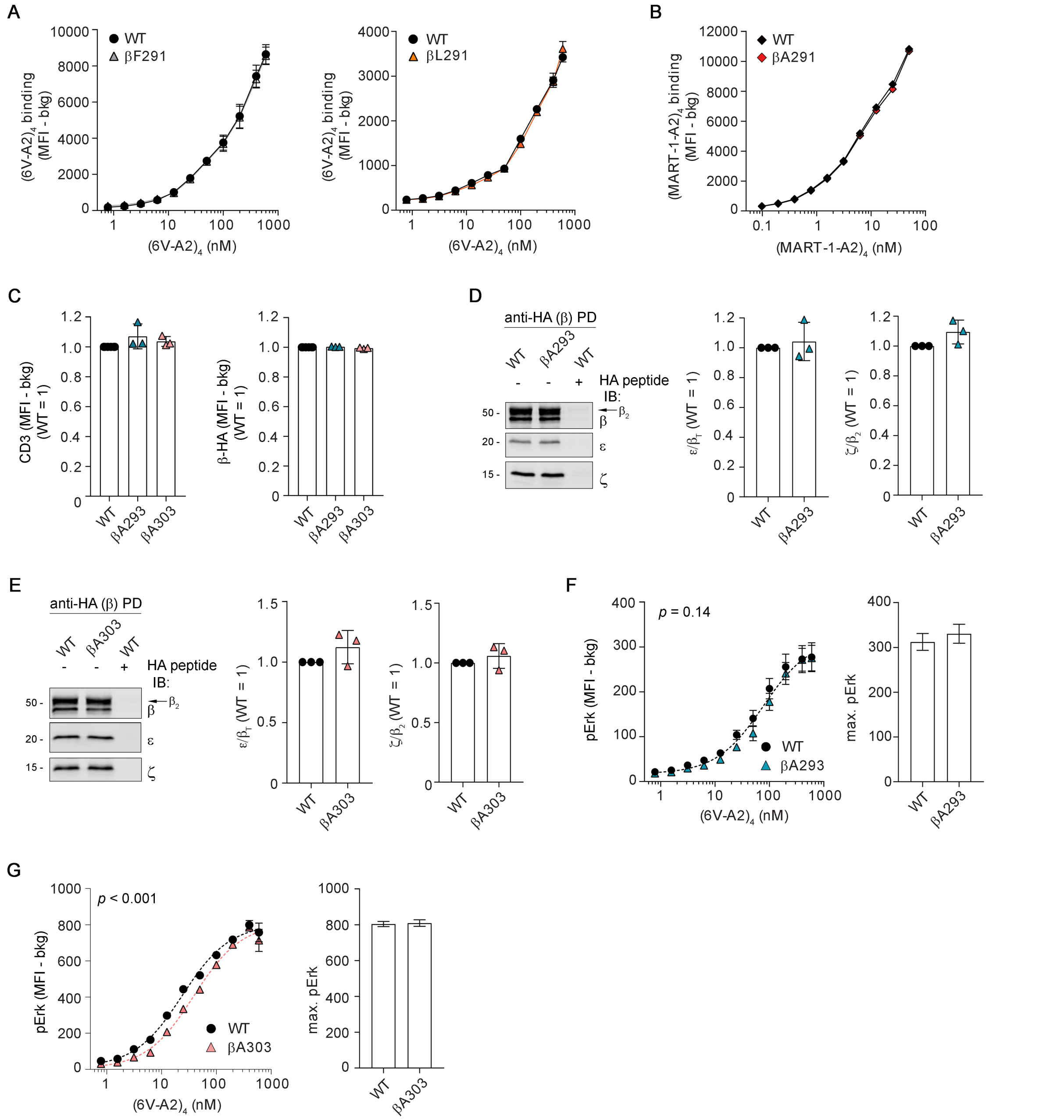


**Figure S3. Loosening ζ association enhances signalling**

**A** $(6V\text{-}A2)_4$ binding to CD8-deficient J76 1G4-WT or 1G4-βF291 (**left**) or 1G4-βL291 (**right**) related to Figs. **3C** and **3D**. Cells were induced with different doses of doxycycline, labelled or not with CellTrace violet, mixed 1:1 and stimulated for 3 min with different doses (0.78 – 600 nM) of PE-conjugated $(6V\text{-}A2)_4$ and analysed by FACS. Plots show mean ± SD of 3 experiments measured in triplicates. **B** $(MART\text{-}1\text{-}A2)_4$ binding to CD8-deficient J76 2H5-WT or 2H5-βA291 related to Fig. **3G**. Cells were induced with different doses of doxycycline, labelled or not with CellTrace violet, mixed 1:1 and stimulated for 3 min with different doses (0.05 – 50 nM) of PE-conjugated $(MART\text{-}1\text{-}A2)_4$ and analysed by FACS. Plot shows mean ± SD of 3 experiments measured in triplicates. **C** TCR-CD3 surface expression in CD8-deficient J76 of 1G4-WT, 1G4-βA293 or 1G4-βA303. Cells were labelled or not with CellTrace violet, mixed 1:1 and analysed for CD3 surface expression and total TCRβ-HA by FACS. Gates for equal HA-tag expression level were applied and the CD3 and β-HA MFI were extracted from the TCRβ-$HA^{low}$ gate, see STAR Methods. **Left**, mean ± SEM of CD3 MFI in $HA^{low}$ gate, 3 experiments measured in duplicates, *t*-test (ns). **Right**, mean ± SEM of β-HA MFI in $HA^{low}$ gate, 3 experiments measured in duplicates, *t*-test (ns). **D** CD8-deficient J76 1G4-WT or 1G4-βA293 were solubilised and analysed by β-HA PD and IB for β, ε and ζ. **Left**, IB: 1 of 3 experiments. The arrow indicates $\beta_2$ isoform. **Middle**, mean ± SD of $\varepsilon/\beta_T$, n = 3, *t*-test (ns). **Right**, mean ± SD of $\zeta/\beta_2$, n = 3, unpaired *t*-test (ns). **E** CD8-deficient J76 1G4-WT or 1G4-βA303 were solubilised and analysed by β-HA PD and IB for β, ε and ζ. **Left**, IB: 1 of 3 experiments. The arrow indicates $\beta_2$ isoform. **Middle**, mean ± SD of $\varepsilon/\beta_T$, n = 3, *t*-test (ns). **Right**, mean ± SD of $\zeta/\beta_2$, n = 3, unpaired *t*-test (ns). **F** pErk response of CD8-deficient J76 1G4-WT or 1G4-βA293. Cells were induced with different doses of doxycycline, labelled or not with CellTrace violet, mixed 1:1 and stimulated for 3 min with different doses (0.78 – 600 nM) of PE-conjugated $(6V\text{-}A2)_4$ and analysed by FACS. **Left**, non-linear regression fit of $(6V\text{-}A2)_4$ nM vs. pErk MFI, n = 3, $R^2$ = 0.52 (WT), 0.60 (βA293). **Right**, mean ± SD of max. pErk, n = 3, F-test (ns). **G** pErk response of CD8-deficient J76 1G4-WT or 1G4-βA303. Cells were induced with different doses of doxycycline, labelled or not with CellTrace violet, mixed 1:1 and stimulated for 3 min with different doses (0.78 – 600 nM) of PE-conjugated $(6V\text{-}A2)_4$ and analysed by FACS. **Left**, non-linear regression fit of $(6V\text{-}A2)_4$ nM vs. pErk MFI, n = 3, $R^2$ = 0.96 (WT), 0.95 (βA293). **Right**, mean ± SD of max. pErk, n = 3, F-test (ns).

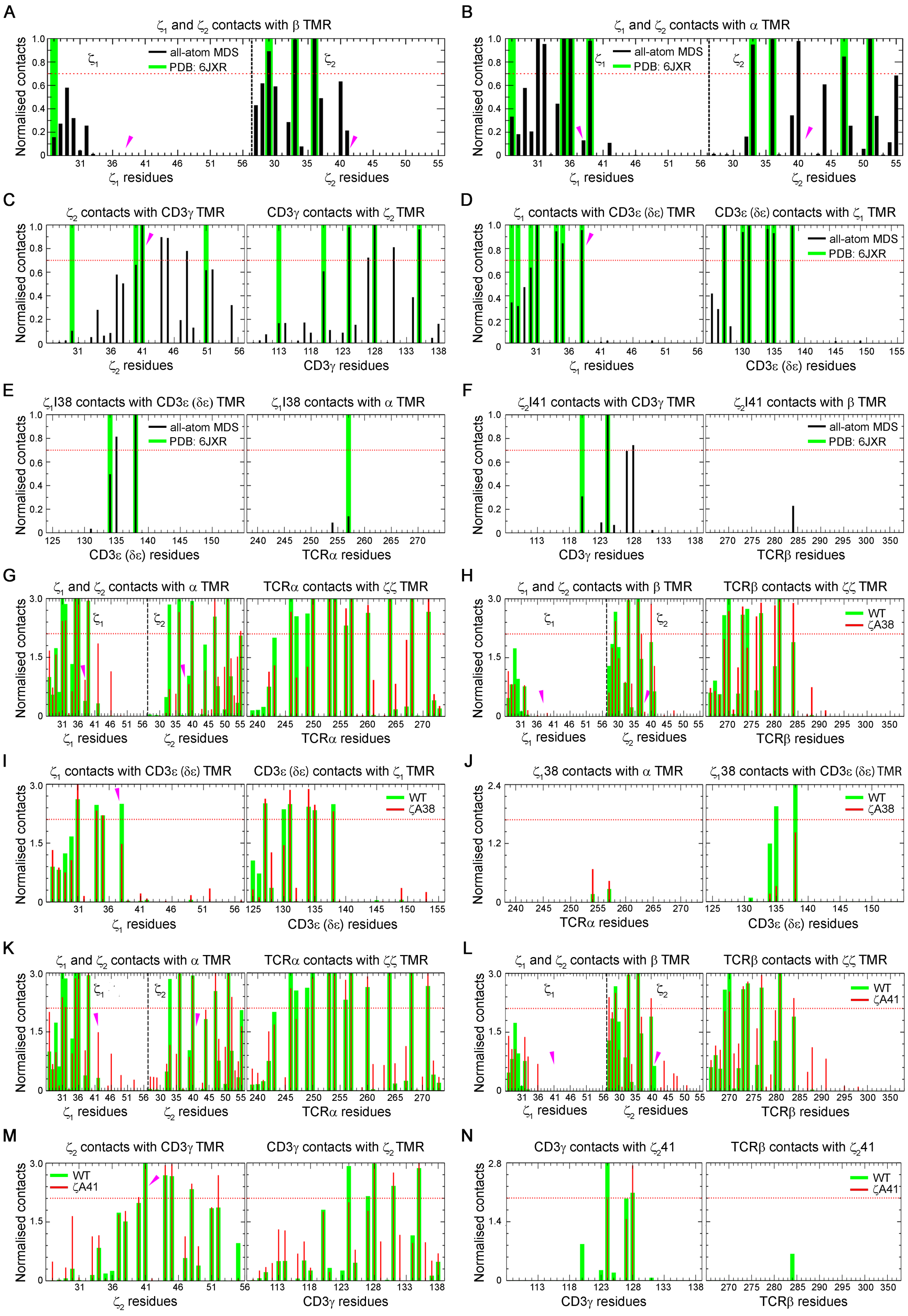


**Figure S4. Loosening ζ association enhances signalling**

**A** Normalised number of contacts of $\zeta_1$ and $\zeta_2$ with β TMR, in our WT all-atom MDS (black) and in the cryo-EM structure (PDB: 6JXR) (green), related to Fig. **4A**. Magenta arrow indicates $\zeta_1$I38 (**left**) or

$\zeta_2$I41 (**right**). In Figs. **S4A** - **S4F**, when comparing the contacts in the cryo-EM structure to our WT all-atom MDS, normalisation was done by dividing the number of contacts of each residue by the highest number of contacts. For all contacts analyses in Figs. **S4A** - **S4N**, a cut-off distance of 4 Å was used to define a contact and the red dotted line represents 70 % of the normalised contacts, a threshold used to measure the significance of contacts. **B** Normalised number of contacts of $\zeta_1$ and $\zeta_2$ with $\alpha$ TMR, in our WT all-atom MDS (black) and in the cryo-EM structure (PDB: 6JXR) (green), related to Fig. **4B**. Magenta arrow indicates $\zeta_1$I38 (**left**) or $\zeta_2$I41 (**right**). **C** Normalised number of contacts of $\zeta_2$ TMR with CD3$\gamma$ TMR (**left**) and of CD3$\gamma$ with $\zeta_2$ TMR (**right**), related to Fig. **4C**. Comparison between protein-protein interactions resulted from our WT all-atom MDS (black) and the cryo-EM structure (PDB: 6JXR) (green). Magenta arrow indicates $\zeta_2$I41. **D** Normalised number of contacts of $\zeta_1$ with CD3$\varepsilon$ ($\delta\varepsilon$) TMR (**left**) and of CD3$\varepsilon$ ($\delta\varepsilon$) with $\zeta_1$ TMR (**right**), related to Fig. **4D**. Comparison between protein-protein interactions resulted from our WT all-atom MDS (black) and the cryo-EM structure (PDB: 6JXR) (green). Magenta arrow indicates $\zeta_1$I38. **E** Normalised number of interactions of $\zeta_1$I38 with the rest of TCR-CD3 TMRs. Comparison between protein-protein interactions resulted from our WT all-atom MDS (black) and the cryo-EM structure (PDB: 6JXR) (green). **F** Normalised number of interactions of $\zeta_2$I41 with the rest of TCR-CD3 TMRs. Comparison between protein-protein interactions resulted from our WT all-atom MDS (black) and the cryo-EM structure (PDB: 6JXR) (green). **G** Normalised number of contacts of $\zeta_1$ and $\zeta_2$ with $\alpha$ TMR (**left**) and of $\alpha$ TMR with $\zeta\zeta$ TMR (**right**) in the WT (green) and $\zeta$A38 (red) all-atom MDS. Magenta arrow indicates $\zeta$38 showing no significant contacts with $\alpha$ TMR in both WT and $\zeta$A38 simulations. However, one contact of $\zeta_2$ ($\zeta_2$Y33) with $\alpha$ TMR and two contacts of $\alpha$ TMR ($\alpha$L247, $\alpha$V249) with $\zeta\zeta$ TMRs are reduced in the simulations carrying $\zeta$A38 substitution. For Figs. **S4G** - **S4N**, when comparing the contacts in the WT simulations to the contacts of the mutants, normalisation was done by dividing the number of contacts of each residue by the number of simulation frames. **H** Normalised number of contacts of $\zeta_1$ and $\zeta_2$ with $\beta$ TMR (**left**) and of $\beta$ TMR with $\zeta\zeta$ TMR (**right**) in the WT (green) and $\zeta$A38 mutant (red) all-atom MDS. Magenta arrow indicates $\zeta$38 showing no significant contacts with $\beta$ TMR in both WT and $\zeta$A38 simulations. However, an increase in the interactions between $\zeta_2$ ($\zeta_2$F40) and $\beta$ ($\beta$S276, $\beta$L280, $\beta$L284) were observed in the simulations carrying $\zeta$A38 substitution. This is likely to be the consequence of the increased $\zeta\zeta$ loosening that allows $\zeta_2$ to wobble and to come in contact with $\beta$ TMR. **I** Normalised number of contacts of $\zeta_1$ with CD3$\varepsilon$ ($\delta\varepsilon$) TMR (**left**) and of CD3$\varepsilon$ ($\delta\varepsilon$) with $\zeta_1$ TMR (**right**) in the WT (green) and $\zeta$A38 mutant (red) all-atom MDS. Magenta arrow indicates $\zeta_1$38 showing that this residue reduced its interaction with CD3$\varepsilon$ ($\delta\varepsilon$) TMR when mutated to alanine ($\zeta$A38, in red), with a net effect of increasing flexibility of both $\zeta_1$ and $\zeta_2$ subunits ($\zeta_1$>$\zeta_2$) relative to TCR$\alpha\beta$ (see also Fig. **4F**). **J** Normalised number of contacts of $\alpha$ TMR (**left**) and of

CD3ε (δε) TMR (**right**) with $\zeta_1$38 in the WT (green) and ζA38 mutant (red) all-atom MDS. A small number of contacts between $\zeta_1$38 and α TMR were observed in both WT and ζA38 simulations while all three residues of CD3ε (δε) TMR (εV134, εI135 and εI138) that interacted in the WT simulations reduced their contacts with $\zeta_1$A38 (red). Normalisation is performed such that the number of contacts of TCRα TMR is compared to that of CD3ε (δε) residues. **K** Normalised number of contacts of $\zeta_1$ and $\zeta_2$ with α TMR (**left**) and of α TMR with ζζ TMR (**right**) in the WT (green) and ζA41 mutant (red) all-atom MDS. Magenta arrow indicates ζ41 showing no significant contacts with α TMR in both WT and ζA41 simulations. However, one contact of $\zeta_2$ ($\zeta_2$Y33) with α TMR and two contacts of α TMR (αL247, αV249) with ζζ TMRs were reduced in the simulations carrying ζA41 substitution. However, one contact of $\zeta_2$ ($\zeta_2$R52) with α TMR increased on ζA41 substitution. **L** Normalised number of contacts of $\zeta_1$ and $\zeta_2$ with β TMR (**left**) and of β TMR with ζζ TMR (**right**) in the WT (green) and ζA41 mutant (red) all-atom MDS. Magenta arrow indicates ζ41 showing no significant contacts of $\zeta_1$41 and $\zeta_2$41 with β TMR in both WT and ζA41 simulations. However, two contacts of $\zeta_2$ with β TMR ($\zeta_2$L27 and $\zeta_2$F40) and one of β TMR with ζζ TMRs (βL284) were increased in the simulations carrying ζA41 substitution. However, one residue of the β TMR (βS269) reduced contact with ζζ TMR during the ζA41 simulations. **M** Normalised number of contacts of $\zeta_2$ with CD3γ TMR (**left**) and of CD3γ with $\zeta_2$ TMR (**right**) in the WT (green) and ζA41 mutant (red) all-atom MDS. Magenta arrow indicates ζ41 showing that the mutated residue $\zeta_2$A41 still maintained contact with CD3γ during the simulations compared to the WT. However, γV124, which strongly interacted with $\zeta_2$I41 in the WT, reduced its interaction with $\zeta_2$A41. One contact of $\zeta_2$ ($\zeta_2$R52) with CD3γ TMR increased in the simulations carrying the ζA41 substitution. **N** Normalised number of contacts of CD3γ TMR (**left**) and of β TMR (**right**) with $\zeta_2$41 in the WT (green) and ζA41 mutant (red) all-atom MDS. Two contacts of CD3γ (γV124 and γF127) with $\zeta_2$41 were reduced and one contact (γV128) was increased during ζA41 simulation. No significant contacts of β TMR with $\zeta_2$41 were observed in both WT and ζA41 simulations.

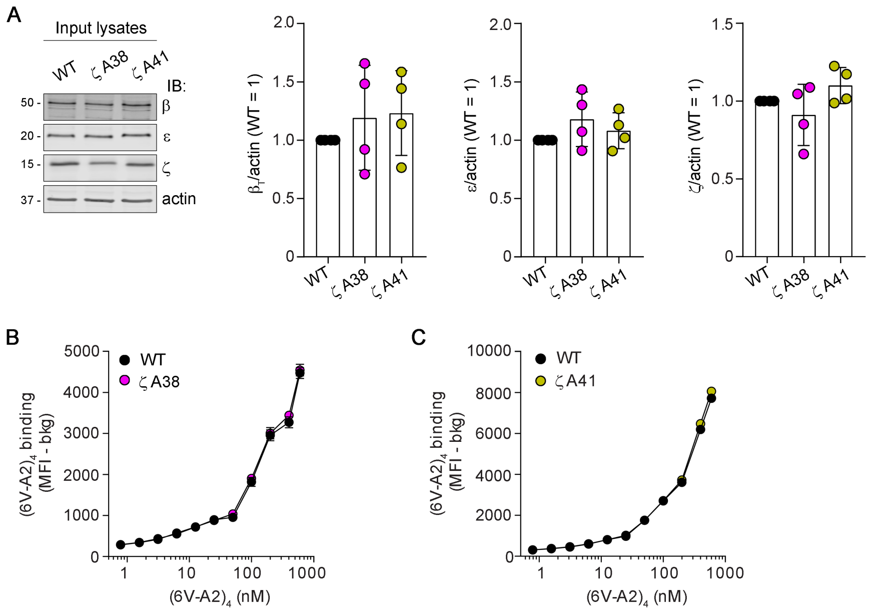


**Figure S5. Loosening ζ association enhances signalling**

**A** J76-1G4WT-ζKO expressing 1G4 ζWT or ζA38 or ζA41 were lysed with 0.5 % DDM and analysed by IB for β, ε, ζ and actin. **Left**, IB of input lysates of the experiment shown in Fig. **5B** (1 of 4 experiments). **Right**, mean ± SD of $\beta_T$/actin, ε/actin and ζ/actin normalized to WT, n = 4, unpaired *t*-test (ns). **B** (6V-A2)$_4$ binding to J76-1G4WT-ζKO expressing ζWT or ζA38 related to Fig. **5C**. Cells were induced with different doses of doxycycline, labelled or not with CellTrace violet, mixed 1:1 and stimulated for 3 min with different doses (0.78 – 600 nM) of PE-conjugated (6V-A2)$_4$ and analysed by FACS. Plot shows mean ± SD of 3 experiments measured in triplicates. **C** (6V-A2)$_4$ binding to J76-1G4WT-ζKO expressing ζWT or ζA41 related to Fig. **5D**. Cells were induced with different doses of doxycycline, labelled or not with CellTrace violet, mixed 1:1 and stimulated for 3 min with different doses (0.78 – 600 nM) of PE-conjugated (6V-A2)$_4$ and analysed by FACS. Plot shows mean ± SD of 3 experiments measured in triplicates.

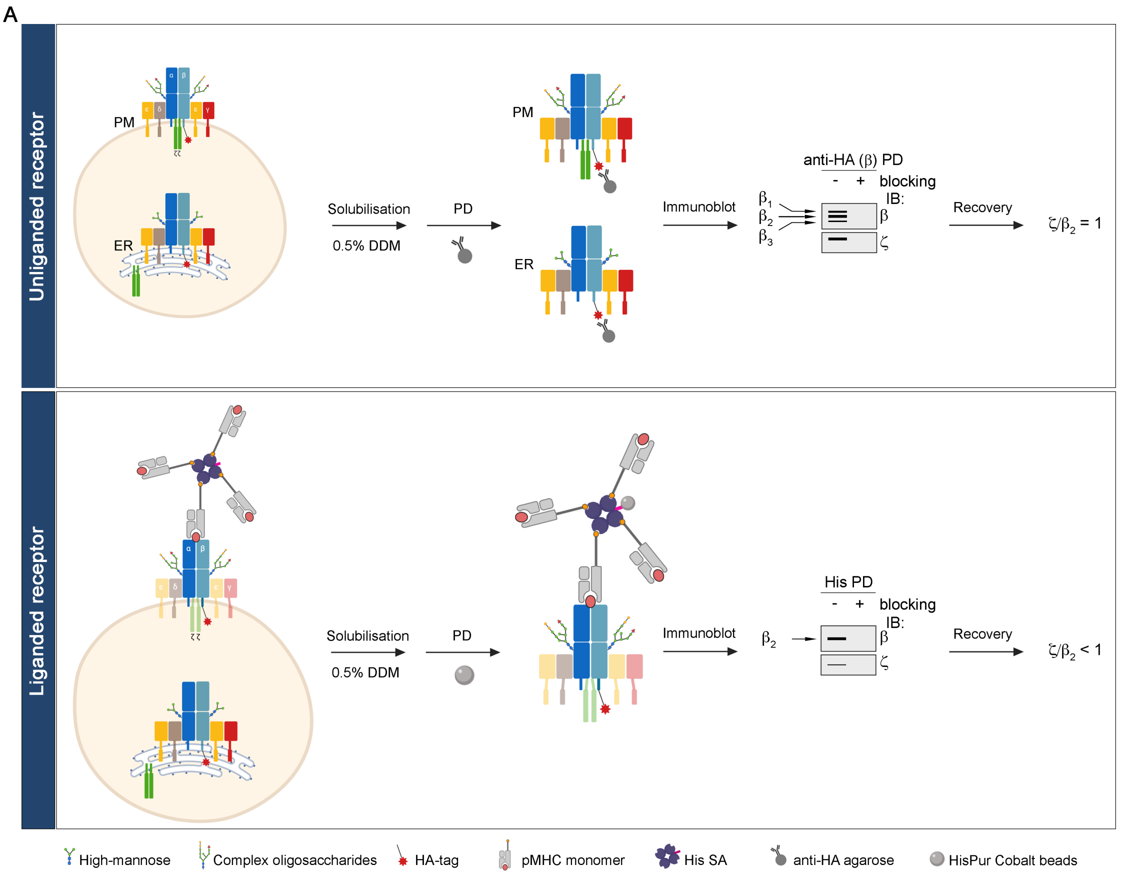


B

| | $K_d$(nM) | CDR1α | CDR2α | CDR3α | CDR1β | CDR2β | CDR3β |
|---|---|---|---|---|---|---|---|
| 1G4 | 1300 | DRGSQS | IQSSQRE | CAVRPTSGGSYIPTFG | MNHEY | SVGAGI | CASSYVGNTGELFFG |
| QM-α | 140 | DRGSQS | IQS**W**QRE | CAVRPTSGGSYIPTFG | MNHEY | SV**AE**GI | CASSYVGNTG**LA**FFG |
| wtc51 | 15 | DRGSQS | IQSSQRE | CAVRPTSGGSYIPTFG | MNHEY | SV**AIQT** | CASSYVGNTGELFFG |

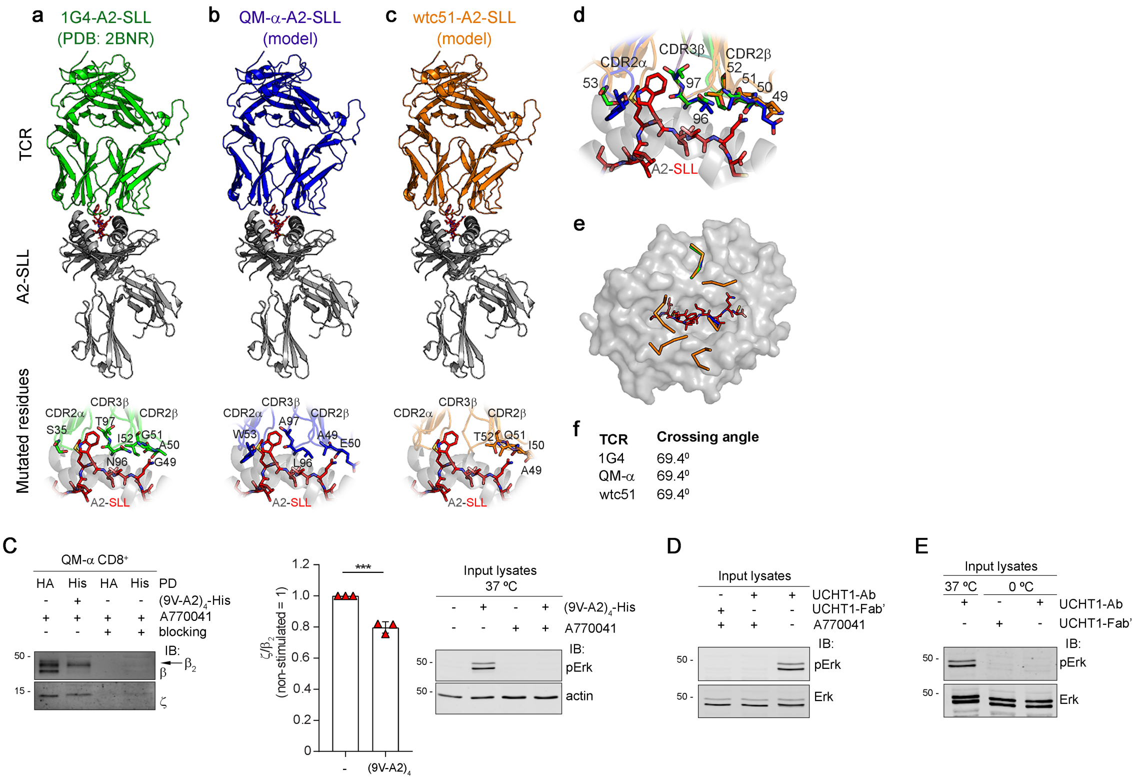


Fig S6

**Figure S6. pMHC tetramer binding loosens αβ association with ζ**

**A** Graphical scheme describing the experimental procedure used to compare the cohesion of unliganded receptor (**top**) and $(pMHC)_4$ ligated TCR-CD3 (liganded receptor, **bottom**). The unliganded receptor was captured by anti-HA (β-HA) PD described in Fig. **S2B**. To pull down the liganded receptor, cells were stimulated with tetramerised His-tagged streptavidin (His SA) and ligand excess was removed. Cells were solubilised with 0.5 % DDM and post-nuclear lysate was incubated with HisPur Cobalt beads to pull down $pMHC_4$-engaged TCR-CD3. The IB schemes on the right show expected bands pattern for β and ζ. Arrows indicate the different isoforms of TCRβ ($\beta_1$, $\beta_2$, $\beta_3$). To evaluate ζ recovery, $\zeta/\beta_2$ ratio was calculated and the value for $\zeta/\beta_2$ ratio from non-stimulated samples was set equal to one. This value represented the recovery of intact TCR-CD3 complex and was compared to $\zeta/\beta_2$ ratio from $(pMHC)_4$ stimulated samples. A ratio < 1 indicates a lower recovery of ζ revealing a reduced cohesion of TCR-CD3 quaternary structure. See STAR Methods for a detailed description of the experimental procedure. **B** Structure and modelling of 1G4-WT and affinity enhanced 1G4 mutants (QM-α and wtc51) used in this study. **Top**, table shows binding affinities of the 1G4 and affinity enhanced TCR mutants (QM-α and wtc51) with sequence alignment highlighting the CDR loops with mutated residues (bold and underlined). **a**, structural overview of the 1G4 TCR (green)-A2 (grey)-SLL (red sticks) tri-molecular complex structure (PDB: 2BNR). Positions of the TCR residues (green sticks) in relation to A2-SLL that are mutated in the affinity enhanced TCRs are shown below. **b**, structural overview of the QM-α TCR (blue)-A2-SLL tri-molecular complex structure (mutations modelled using PDB: 2BNR). Positions of the TCR residues (blue sticks) in relation to A2-SLL that are mutated in the affinity enhanced TCR are shown below. **c**, structural overview of the wtc51 TCR (orange)-A2-SLL tri-molecular complex structure (mutations modelled using PDB: 2BNR). Positions of the TCR residues (orange sticks) in relation to A2-SLL that are mutated in the affinity enhanced TCRs are shown below. **d**, overlay of the mutated residues in the CDR loops comparing 1G4 (green sticks), QM-α (blue sticks) and wtc51 (orange sticks) TCRs. **e**, overlay of the positions of the CDR loops comparing 1G4 (green ribbon), QM-α (blue ribbon) and wtc51 (orange ribbon). **f**, analysis of the TCR crossing angles for each TCR. **C** J76 QM-α treated with A770041 and stimulated or not with $(9V\text{-}A2)_4$-His. **Left**, β-HA (lanes 1, 3) or His (lanes 2, 4) PD and IB for β and ζ (1 of 3 experiments). The arrow indicates $\beta_2$ isoform. **Middle**, mean ± SD of $\zeta/\beta_2$, n = 3, unpaired *t*-test $p < 0.001$. **Right**, pErk IB: 1 of 3 experiments. **D** J76 1G4 ± A770041 were incubated with or w/o UCHT1-Fab' or UCHT1-Ab. pErk IB of the experiment shown in Fig. **6G** (1 of 3 experiments). **E** J76 1G4 were cooled on ice for 20 min and incubated for 5 min on ice with UCHT1-Fab' or UCHT1-Ab. pErk IB of the experiment shown in Fig. **6H** (1 of 3 experiments).

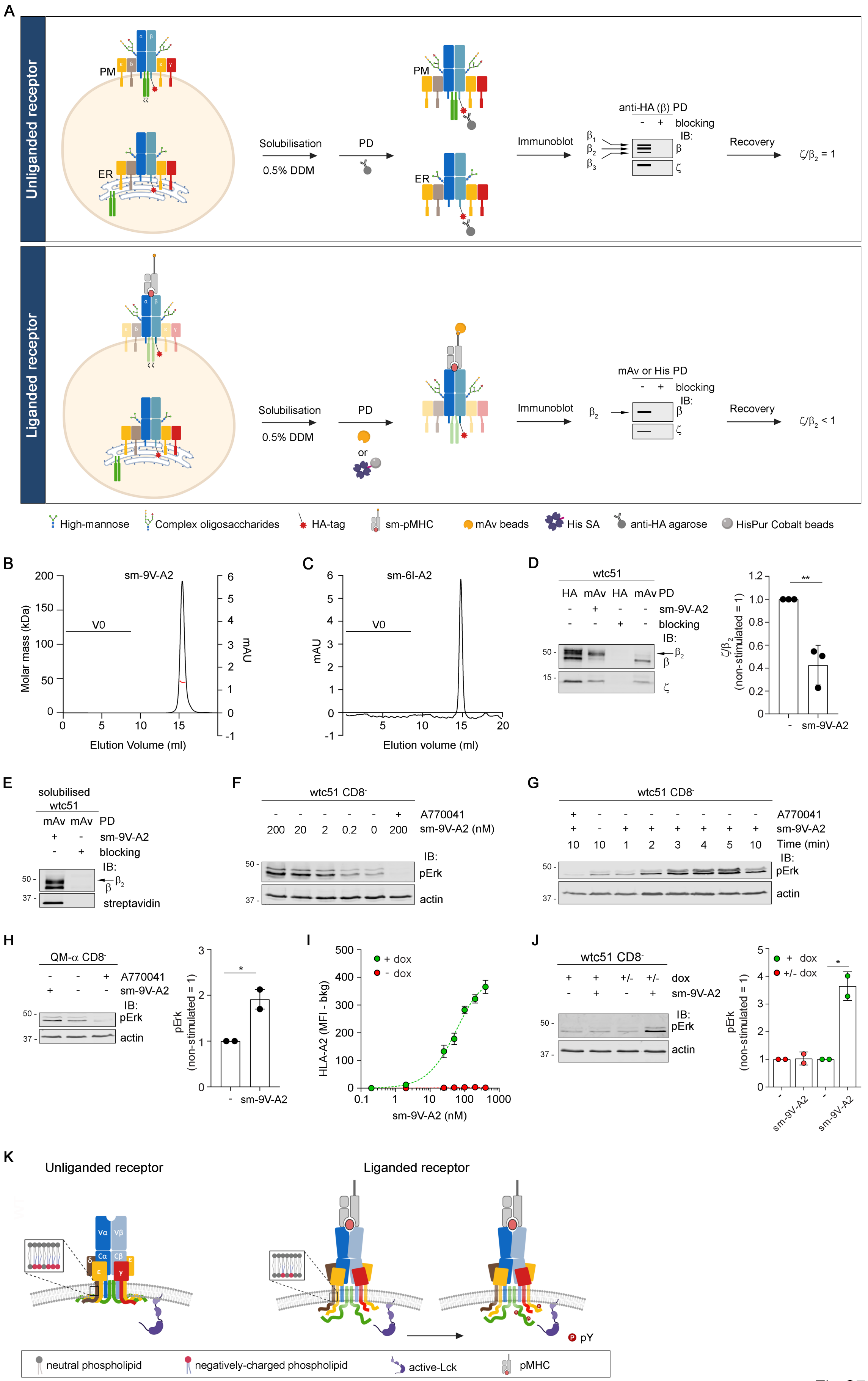
A
Unliganded receptor
Liganded receptor
PM
ER
Solubilisation
0.5% DDM
PD
Immunoblot
anti-HA (β) PD
blocking
IB:
Recovery
ζ/β2 = 1
mAv or His PD
ζ/β2 < 1
High-mannose
Complex oligosaccharides
HA-tag
sm-pMHC
mAv beads
His SA
anti-HA agarose
HisPur Cobalt beads
B
sm-9V-A2
Molar mass (kDa)
mAU
V0
Elution Volume (ml)
C
sm-6I-A2
Elution volume (ml)
D
wtc51
HA mAv HA mAv PD
sm-9V-A2
blocking
pErk
actin
E
solubilised wtc51
streptavidin
F
wtc51 CD8-
A770041
sm-9V-A2 (nM)
G
Time (min)
H
QM-α CD8-
pErk (non-stimulated = 1)
I
+ dox
- dox
HLA-A2 (MFI - bkg)
sm-9V-A2 (nM)
J
dox
K
Unliganded receptor
Liganded receptor
neutral phospholipid
negatively-charged phospholipid
active-Lck
pMHC
pY

Fig S7

**Figure S7. Monovalent pMHC in solution triggers TCR-CD3 untying and intracellular signalling**

**A** Graphical scheme describing the experimental procedure used to compare the cohesion of unliganded receptor (**top**) and soluble, monovalent, mono-dispersed (sm)-pMHC ligated TCR-CD3 (liganded receptor, **bottom**). The unliganded receptor was captured by anti-HA (β-HA) PD described in Fig. **S2B**. To pull down the liganded receptor cells were stimulated with biotinylated sm-pMHC and ligand excess was removed. Cells were then solubilised with 0.5 % DDM and post-nuclear lysate was incubated with His-tagged streptavidin (His SA) followed by pull down with HisPur Cobalt beads or post-nuclear lysate was incubated with monomeric Avidin beads (mAv). The IB schemes on the right show expected bands pattern for β and ζ. Arrows on the left indicate the different isoforms of TCRβ ($\beta_1$, $\beta_2$, $\beta_3$). To evaluate ζ recovery, $\zeta/\beta_2$ ratio was calculated and the value for $\zeta/\beta_2$ ratio from the non-stimulated sample was set equal to one. This value represented the recovery of intact TCR-CD3 complex and was compared to $\zeta/\beta_2$ ratio from sm-pMHC stimulated samples. A ratio < 1 indicates a lower recovery of ζ revealing a reduced cohesion of TCR-CD3 quaternary structure. See STAR Methods for a detailed description of the experimental procedure. **B** Size-exclusion chromatography-multi-angle-light scattering analysis of sm-9V-A2 showing a single, homogeneous peak of 44.6 kDa. The red line indicates the Molar mass (kDa). V0: void volume. **C** Gel-filtration chromatogram of sm-6I-A2 showing a single, homogeneous peak. V0: void volume. **D** J76 wtc51 stimulated or not with sm-9V-A2. **Left**, β-HA (lanes 1, 3) or monomeric Avidin (mAv) (lanes 2, 4) PD and IB for β and ζ (1 of 3 experiments). The arrow indicates $\beta_2$ isoform. **Right**, mean ± SD of $\zeta/\beta_2$, n = 3, unpaired *t*-test $p < 0.01$. **E** CD8-deficient J76 wtc51 were lysed, incubated or not with sm-9V-A2 and subjected to PD by anti-HA or monomeric Avidin (mAv). Monomeric Avidin (mAv) PD and IB for β and streptavidin IRDye 800CW. The arrow indicates $\beta_2$ isoform. **F** CD8-deficient J76 wtc51 ± A770041 were stimulated or not with the indicated concentrations of sm-9V-A2 for 5 minutes. pErk IB representative of 2 experiments. **G** CD8-deficient J76 wtc51 ± A770041 were stimulated or not with sm-9V-A2 for the indicated time points. pErk IB representative of 2 experiments. **H** CD8-deficient J76 QM-α ± A770041 were stimulated or not with sm-9V-A2. **Left**, pErk IB (1 of 2 experiments). **Right**, mean ± SEM of pErk, n = 2, unpaired *t*-test $p = 0.05$. **I** Sm-9V-A2 binding to CD8-deficient J76 wtc51. Cells were induced or not with doxycycline (dox), labelled with CellTrace violet or left untreated, mixed 1:1 and stimulated for 5 min with different doses (0.02 – 400 nM) of sm-9V-A2 at 37 $^{o}$C. Cells were rapidly washed with FACS buffer, stained with anti-HLA-A2 and analysed by FACS. Non-linear regression fit of sm-9V-A2 (nM) vs. HLA-A2 (MFI), n = 3. **J** CD8-deficient J76 wtc51 not doxycycline-induced (- dox) for TCR expression were reacted or not with 200 nM sm-9V-A2 for 5 minutes at 37 $^{o}$C, washed and mixed with equal number of dox-induced CD8-deficient J76 wtc51 (mix ± dox), see STAR Methods for a detailed protocol. Dox-induced (+ dox) CD8-deficient J76 wtc51 stimulated or not with 200 nM sm-9V-A2 for 5 minutes at 37

$^{o}$C served as control. **Left**, pErk IB (1 of 2 experiments). **Right**, mean ± SD of pErk of mixed dox-induced and not dox-induced ± sm-9V-A2 cells (mix ± dox), n = 2 experiments in duplicates, unpaired *t*-test: ns (mix ± dox), $p < 0.05$ (+ dox). **K** Graphical scheme describing the "TCR-CD3 allosteric relaxation" mechanism uncovered in this study. Briefly we suggest that in absence of force, co-receptor, clustering or PTPs exclusion, monovalent pMHC binding to TCR-CD3 allosterically regulates a cascade of conformational changes that relaxes the quaternary structure of TCR-CD3 TMRs. In the proposed model, conformational changes occurring at the pMHC binding site propagate to C$\alpha$C$\beta$ ECDs at the site where they contact the CD3 subunits. These rearrangements are transmitted to the CD3 TMRs, resulting in a reduced cohesion of TCR-CD3 TMRs, ITAMs exposure and their phosphorylation by active Lck. Moreover, we envisage the possibility that pMHC-induced reconfiguration of the octamer's TMRs leads to a local redistribution of negatively-charged lipids which could reduce hydrophobic and electrostatic forces that help holding CD3 tails within the plasma membrane.

### STAR METHODS

### CONTACT FOR REAGENTS AND RESOURCES SHARING

Further information and requests for resources and reagents should be directed to and will be fulfilled by the Lead Contact, Oreste Acuto. There are no restrictions on any data or materials presented in this paper.

### EXPERIMENTAL MODEL AND SUBJECT DETAILS

#### Cell lines

Cell lines were maintained at 37 $^{o}$C with 5 % $CO_2$ in a humidified incubator (Heraeus). Human embryonic kidney epithelial 293T cells HEK293T (ATCC CRL-3216) and Lenti-X293T (Clontech) cell lines were cultured in complete DMEM (Sigma Aldrich) supplemented with 15 % fetal bovine serum (FBS). Jurkat-derived cell line variants 31.13 (Alcover et al., 1990) (abbreviated J31.13) and J76 (Heemskerk et al., 2003), which lack expression of TCRβ and TCRαβ respectively, were cultured in RPMI 1640 supplemented with 10 % FBS. **Key Resources Table** lists all the above cell lines and those that were derived from J31.13 and J76 in this study. Cells were routinely tested and found negative for mycoplasma and are not STR profiled. Jurkat cell lines containing a tetracycline-inducible gene expression systems were maintained in RPMI 1640 supplemented with 10 % tetracycline-negative FBS (Clontech).

#### Primary T cells

Primary CD8 T cells were obtained from HLA-A2$^{-}$ healthy donor blood cones (NHS) by density gradient centrifugation and negative selection using EasySep Human CD8 T cell isolation kit (STEMCELL). Isolated T cells were rested overnight at 2 x $10^{6}$/ml in RPMI medium (Sigma) supplemented with 2 mM Glutamine, 1 mM Sodium Pyruvate, Non-essential AA, 50 μg/ml Kanamycin, 5 % human serum (Sigma).

### METHODS DETAILS

#### DNA constructs and cloning

All plasmid DNA was propagated in *E. coli* strains DH5α or Stbl3.

<u>Generation of 1G4 α and 1G4 β plasmids for transient transfections</u>

1G4 α and β sequences were amplified from total cDNA extracted (Roche cDNA extraction kit) from an established 1G4 expressing Jurkat cell line (Aleksic et al., 2010) with appropriate primers (see primers in **Table S2**) and cloned into pEF3 with a Flag- and HA-tag respectively.

Site directed mutagenesis

Single amino acid mutations in the 1G4 β chain were introduced using a modified QuikChange protocol (Agilent). Briefly, PCR primers were generated using the QuikChange open source software. PCR was carried out with an optimised protocol and the resulting product was purified using QiaQuick Gel purification kit (Qiagen). A fraction of purified PCR was digested with DpnI for 2 h and used to transform Stbl3 *E. coli* (Invitrogen). DNA was extracted from colonies, analysed by gel electrophoresis and sequenced (Source BioScience. UK).

Generation of the self-cleavable 1G4 αβ construct (stable cell lines)

A DNA construct was designed to obtain a single mRNA for β and α giving rise to a single polypeptide separated by the foot-and-mouth disease virus 2A (F2A) self-cleavable sequence (Ryan et al., 1991). This strategy should facilitate expression of similar amounts of β and α proteins. The aa sequence of the entire construct is provided in **Table S1**. See Fig. **S1A** for a schematic of the construct.

Constructs for Tet-inducible expression of 1G4 αβ, 2H5 αβ, ζ WT and mutants, affinity enhanced 1G4 (wtc51 and QM-α), ζTST and constitutive 1G4 αβ and 868 TCR

To generate an inducible expression system for 1G4 αβWT and 1G4 αWT/βA291, the sequence coding for the self- cleavable single polypeptide β-HA-F2A-α-FLAG was cloned into the doxycycline inducible Gateway cloning plasmid pLIX-402 (Addgene) following the manufacturer's guidelines for Gateway cloning. Briefly, the DNA sequence containing the entire αβ sequence was PCR amplified from the constitutive expression plasmid pHR with primers containing the Gateway recombination sites (see primers for Gateway cloning in **Table S2**). Purified PCR products were inserted via the BP recombination reaction into the Gateway entry vector pDONR-221 (Thermo Fisher). After verification of successful recombination by automated sequencing (Source Bioscience, UK), the resulting entry clones were used for a LR recombination reaction to insert the sequence into the pLIX-402 destination vector (Addgene), which carries a tetracycline-inducible promoter for conditional expression of the gene of interest. Both BP and LR reactions were performed with BP and LR Clonase II kits (Thermo Fisher). To clone the high affinity TCRs (wtc51 and QM-α) and the entire panel of 1G4 and 2H5 β and ζ mutants, synthetic DNA fragments coding for either self-cleavable single polypeptide 1G4 αβ (αWT plus βWT or mutants βA291, βL291, βF291) or 2H5 WT and 2H5 βA291 or ζWT and mutants ζA38 and ζA41 were purchased from GeneArt Gene Synthesis (Thermo Fisher). All DNA fragments contained consensus sequences for Gateway Cloning System and the αβ and ζ constructs were introduced into the Gateway plasmid pLIX-402 for inducible expression, following the Gateway cloning protocol as described above. After cloning into the expression vectors, the coding sequences were verified by automated sequencing (Source BioScience, UK). To generate a twin-strep tagged (TST) ζ containing cell-line the sequence of ζ was cloned, after XbaI/EcoRI digestion, into the plasmid pEXP103 IBA (IBA Lifesciences), to be fused with a TS-tag at the C-terminus. The ζTST sequence was subsequently amplified and cloned into pLIX-402 destination vector following the gateway procedure as described above. (See primers for Gateway cloning in **Table S2**). To generate a constitutive expression system for 1G4 αβ and 868 αβ synthetic DNA fragments coding for either self- cleavable single polypeptide 1G4 αβ and 868 αβ were purchased from GeneArt Gene Synthesis and were introduced into the Gateway plasmid pLEX 307 (Addgene) for constitutive expression, following the Gateway cloning protocol as described above.

**CRISPR/Cas9-mediated ζ knock-out**

CRISPR/Cas9 mediated knock-out (KO) of ζ was achieved using the pX458 two component vector (Addgene) and the guide sequence CAGGCACAGTTGCCGATTACAGG. J76 expressing CD8 and the single polypeptide 1G4 αβ were transiently transfected and CD3 negative cells were sorted. Efficient knock-out was verified by transient transfection of the proteins of interest and analysis of rescue by FACS analysis.

**TCR-CD3 expression**

Transient expression of TCR-CD3

J31.13 cells were grown to approximately 0.5 – 0.6 x $10^6$/ml, cells were counted, harvested (2 x $10^6$ cells per electroporation) and centrifuged at 1,200 rpm. The supernatant (conditioned medium) was collected, diluted with equal volume of RPMI containing 10 % FBS and kept at 37 $^o$C in the incubator. Cells were washed twice in RPMI and re-suspended in 400 μl RPMI and placed in sterile Biorad Genepulser Cuvettes (0.4 cm) containing 2 μg of plasmid DNA per $10^6$ cells. Constructs and mutant DNA were routinely purified from *E. coli* using Qiagen mini- or midi-prep kits (Qiagen). Electroporation was performed with a Biorad GenePulser X-cell at 260 V, 1 pulse of 25 ms using the square root method. Cells were flushed out of the cuvette with 200 μl conditioned medium and placed at 37 $^o$C in the incubator until final use. Cells were usually assayed after 24 - 48 h and analysed by flow cytometry for CD3 surface expression or for pErk in response to pMHC tetramer.

Generation of stable cell lines

For lentivirus infection, recipient cells were cultured up to approximately 0.5 – 0.6 x $10^6$/ml. Cells were washed once in RPMI, checked for viability, counted and adjusted to $10^6$/ml in RPMI supplemented with 10 % FBS (tetracycline-free (Tet$^-$) in the case of transduction with doxycycline-inducible plasmids) (Gibco or Clontech) and 5 μg/ml of polybrene (Sigma). Cells were plated in 12-well (1 ml/well) or 6-wells plates (2 ml/well) and one aliquot of lentivirus was added (for preparation of lentiviruses see below). 24 h post-infection, cells were washed and re-suspended in RPMI 10 % FBS (standard or Tet$^-$). After transduction with a plasmid containing an antibiotic selection marker, cells were cultured for an additional 24 h before starting selection with puromycin (1 μg/ml, Gibco). With adequate lentivirus concentration, cell mortality was relatively low.

For tetracycline-inducible cell lines, gene expression could be started after 72 h by adding 0.1 - 1 μg/ml of doxycycline hyclate (Sigma Aldrich). Cells were collected after approximately 16 h and tested for CD3 expression and signalling by flow cytometry.

Primary T cells were plated in 24-well (1 ml/well) at 1 x $10^6$/ml and stimulated with human T-activator CD3/CD28 Dynabeads (Thermo Fisher Scientific) at 2:1 bead to T-cell ratio in complete medium supplemented with 500 U/ml IL-2. After 48 h most of the medium was removed and replaced with fresh medium supplemented with 500 U/ml IL-2, containing 40 μl of concentrated lentiviral stock in presence of 5 μg/ml polybrene (Sigma). After 4 days the dynabeads were removed and 1G4 wtc51 positive cells were sorted after $(9V\text{-}A2)_4$-PE staining. Sorted cells were kept in complete medium supplemented with 500 U/ml IL-2 and re-stimulated with human T-activator CD3/CD28 Dynabeads 12 days after the sorting.

Production of lentiviral particles

Lentiviruses were generated using the packaging cell lines HEK293T or Lenti-X293T. The culture medium was exchanged with RPMI with 10 % FBS just prior to transfection. HEK293T or Lenti-X293T at 80 % confluence were transfected using PEIpro (Polyplus) according to the manufacturer's instructions or by a standard calcium-phosphate precipitation protocol. The packaging plasmids pVSVG and pSPAX2 were mixed with the lentivirus expression vectors containing the gene of interest. For the PEIpro transfection, PEIpro solution was added to the plasmids mix and immediately vortexed, left 15 min at room temperature (RT) and then added drop-wise to the cells by gently swirling the plate. For calcium-phosphate precipitation, cells were left 3 h with DNA-calcium-phosphate precipitate and the media replaced with complete DMEM with 15 % FBS. Independently of the transfection protocol, supernatant containing lentiviral particles was collected after 48 h and filtered through a 0.45 μm sterile filter (Sartorius Stedim). Lentivirus supernatants were either used immediately or concentrated with PEG-*it*™ (SBI) concentration kit according to the manufacturer's instruction. Briefly, lentiviral supernatants were mixed with Virus Precipitation Solution (SBI) to a final concentration of 1X Virus Precipitation Solution and incubated overnight at 4 $^{o}$C followed by a centrifugation at 1,500 x g for 30 min at 4 $^{o}$C. The pellet containing lentivirus particles was re-suspended in 1/100 of the volume of the original cell culture using cold RPMI. Aliquots were immediately frozen in cryogenic vials at - 80 $^{o}$C and stored until use. Aliquots of each lentivirus batch were routinely pre-tested by serial dilution titration. Frozen aliquots were thawed only once and used immediately with minimal loss of virus titer as determined by flow cytometry.

**Preparation of pMHC monomers and tetramers**

pMHC monomers were produced as described elsewhere (Altman and Davis, 2003) with some modifications. Human beta-2 microglobulin (β2m) and the 1 - 278 segment of HLA-A*02:01 heavy chain with the AviTag at the C-terminus were separately expressed in *E. coli*. Both proteins were recovered from inclusion bodies with Bug Buster protein extraction reagent (Millipore) supplemented with lysozyme and benzonase (Millipore). For monomer refolding, HLA heavy chain and β2m were added to refold buffer (100 mM Tris, 400 mM L-arginine hydrochloride, 2 mM EDTA, 5 mM reduced glutathione, 0.5 mM oxidised glutathione, 0.1 mM PMSF) supplemented with 0.5 mM urea and 10 μg/ml synthetic peptide (> 95 % purity by HPLC) of either one the following variants of the NY-ESO-1 antigen (Aleksic et al., 2010) (SLLMWITQC, residues 157-165): 9V (SLLMWITQV, $K_d$ 7.2 ± 0.5 μM), 6V (SLLMWVTQV, $K_d$ 18 μM) or the MART-1 tumour antigen (MART-1 26-35 (ELAGIGILTV) (Circosta et al., 2009) or the 6I (SLYNTIATL, Cambridge Peptides Inc.). The resulting soluble-monovalent-monodisperse pMHC (sm-pMHC) were indicated as: 9V-A2, 6V-A2, MART-1-A2 and 6I-A2.

Peptides were synthesized by standard solid-phase chemistry using F-moc for transient N-terminal protection. All peptides were approximately 98 % pure as determined by analytical HPLC and mass spectrometry. Lyophilized peptides were dissolved in DMSO (Sigma) to 10 mg/ml and aliquoted of 100 μl in low-bind Eppendorf tubes and used immediately or stored at - 80 $^{o}$C. The refold mixture was stirred gently for 40 h at 4 $^{o}$C, concentrated approximately 40-fold to 2.5 ml by centrifugation at 4 $^{o}$C at 3,500 x g in Centricone® Plus-70 centrifugal filter units (30 kD cut-off) (Merck Millipore) and desalted on a disposable PD 10 column (GE Healthcare) by gravity flow eluting in 3.5 ml TBS. The eluent was subject to mono-biotinylation using a BirA kit (Avidity) and fractionated by FPLC using a HiLoad™ 16/600 Superdex™ 200 pg column (GE Lifesciences) and the ÄKTA Pure (GE Healthcare) system. Fractions containing rHLA-A*02:01-β2m dimers were pooled and concentrated in Amicon Ultra 15 ml centrifugal filter units (10 kD cut-off) (Merck Millipore). After addition of glycerol to a final concentration of 5 %, protein concentration was adjusted to 1 mg/ml measuring $OD_{280}$ and aliquots were frozen at - 80 $^{o}$C until use. We rigorously controlled the quality of monomers prior to performing experiments by size-exclusion chromatography (SEC) and we also used multi-angle light scattering with size-exclusion chromatography (SEC-MALS) for sm-9V-A2 to further control for mono-dispersity and unique molecular mass in the peak.

Tetramers were generated by slowly mixing aliquots of biotinylated rHLA-A*02:01-rβ2m with phycoerythrin (PE)-conjugated streptavidin (Sigma) at RT under constant agitation as described earlier (Altman and Davis, 2003). Alternatively, His-streptavidin (Antibodies Online) was used. The resulting tetramers, $(9V\text{-}A2)_4$, $(6V\text{-}A2)_4$, $(MART\text{-}1\text{-}A2)_4$ and $(6I\text{-}A2)_4$, were stored at 4 $^{o}$C until final use.

**Preparation of UCHT1-Fab' fragment**

A Fab' fragment was prepared from the purified mAb UCHT1 (BioXCell). Briefly, UCHT1 was digested with Pepsin (Thermo Fisher) in 100 mM Sodium Acetate (Thermo Fisher) for 24 h at 37 $^{o}$C on a rocking platform. The digest was incubated overnight at 4 $^{o}$C on a rotating shaker with Protein G agarose (Thermo Fisher) to remove the Fc fragment. The resulting $Fab_2$ was reduced with 2-Mercaptoethylamine-HCl (MEA-HCl) (Thermo Fisher) for 90 min at 37 $^{o}$C, according to the manufacturer's instructions, to obtain the Fab' fragment. MEA-HCl excess was removed using a Zeba Spin Desalting column (Thermo Fisher) and the Fab' fragment eluted in Phosphate Buffered Saline (PBS, Sigma) at pH 7.0. The UCHT1-Fab' was conjugated to biotin using EZ-Link Maleimide-PEG2 biotin (Thermo Fisher) or to Alexa-Fluor 647 or 488 using Alexa-Fluor 647 $C_2$ Maleimide and Alexa-Fluor 488 $C_5$ Maleimide respectively (Thermo Fisher). The Fab' was mixed in 1:10 molar ratio with 2 mM EZ-Link Maleimide-PEG2 biotin or with Alexa-Fluor 647 $C_2$ Maleimide for 5 h with slight agitation on a rocking platform at RT. Reagent excess was removed using a Zeba Spin Desalting column. The biotin, Alexa-Fluor 488 or Alexa-Fluor 647-labelled UCHT1-Fab' fragments were analysed on a non-reduced 4 - 15 % Mini-PROTEAN Precast protein gel (Bio-Rad) and stored in PBS at 4 $^{o}$C. The biotin-labelled UCHT1-Fab' was further analysed by FPLC to detect formation of Fab' aggregates. Biotin-labelled UCHT1-Fab' was diluted to a concentration of 0.3 mg/ml in PBS and 40 μl were analysed by size exclusion chromatography. The analysis was carried out using a Superdex™ 200 increase 5/150 GL column (GE Lifesciences) and the ÄKTA Pure system (GE Lifesciences). Fab' was applied onto the column using a 40 μl loop at a flow rate of 100 μl/ml. Isocratic elution with PBS occurred at 300 μl/min with a total elution volume of 3 ml.

**Flow cytometry**

General staining procedures

This section provides a description of the procedures used for antibody (Ab)-mediated cell staining for flow cytometry and data acquisition. Staining was performed in 96-well V-bottom plates (Thermo Fisher) and centrifugations carried out at 670 x g for 1 min in a plate centrifuge (Eppendorf Centrifuge 5810R or Thermo IEC CL30 Centrifuge). Care was taken to gently re-suspend cells after each centrifugation to avoid cell damage and/or death. To optimise the sensitivity of the flow cytometry-based assays, Abs used in this study were systematically titrated for staining of cells expressing or lacking the target antigen and the conditions showing the best signal-to-noise were chosen. Flow cytometry staining was routinely carried out in duplicates or triplicates. For cell surface staining, 0.1 x $10^6$ cells/sample were transferred into a 96-well V-bottom plate, washed once in 150 μl FACS buffer (0.5 % bovine serum albumin (BSA) (Sigma) in PBS). After spinning, supernatants were removed and cell pellets re-suspended in 50 μl staining solution containing fluorescence-conjugated primary Ab (see **Key Resources Table** for a list of the Abs used in this study) diluted in FACS buffer and incubated for 20 min at RT or for 30 min at 4 $^oC$, depending on the Ab used. After removal of the staining solution, samples were washed twice with 150 μl FACS buffer and flow cytometry data acquired immediately or cells were fixed with 150 μl pre-warmed fixation solution (BD Cytofix®, BD Biosciences) for 10 min at 37 $^oC$. For staining of intracellular antigens, fixed samples were washed twice in 150 μl permeabilisation buffer (BD Perm/Wash I, BD Biosciences), re-suspended in 150 μl permeabilisation buffer and incubated at RT for 30 min. Permeabilised cells were stained in 50 μl permeabilisation buffer containing the desired Ab dilution. For fluorescent-conjugated primary Ab staining, samples were incubated at RT for 1 h (except for anti-pY142 CD3ζ and anti-HA, which were incubated at 4 $^oC$ for 2 h). When fluorescent-conjugated secondary Abs were used, they were diluted in 50 μl permeabilisation buffer and added to cells for 20 min at RT. After each staining, cells were washed 3 times with 150 μl permeabilisation buffer and once with 150 μl FACS buffer at the end of the staining procedure. When surface co-staining was included, it was performed prior to fixation as described above. Samples were left in FACS buffer for storage and acquisition and acquired on a CyAn™ ADP analyzer (Beckman Coulter) or BD LSR Fortessa X20 (BD Bioscience) as specified. Raw data was analysed by FlowJo (FlowJo Software part of BD). Counts, percentages or median intensity fluorescence values (MFI) were extracted from FlowJo as excel files. Statistical analysis and non-linear regression were performed with Prism (GraphPad Software).

CellTrace violet labelling

Cells were washed once in PBS and adjusted to a final concentration of $10^6$ cells/ml in pre-warmed PBS at 37 $^oC$. CellTrace violet (Thermo Fisher) or carrier control DMSO (Sigma) was added to reach the indicated staining concentration and cells were incubated at 37 $^oC$ in the dark. After 20 min, samples were diluted 5-fold in complete medium and incubated for an additional 5 min at 37 $^oC$ in the dark. After removal of the diluted staining solution, cells were re-suspended in complete medium, counted and cultured or mixed as indicated.

UCHT1-Fab' fragment binding

J76 1G4 WT or β mutants were induced for TCR expression with doxycycline 1 μg/ml for 96 h. TCR-deficient J76 $CD8^+$ cells were used to evaluate the background. Cells were harvested and stained with AF488-conjugated UCHT1-Fab' at the indicated dilutions in FACS buffer for 30 min at 4 $^oC$. After removing excess of UCHT1-Fab', cells were fixed, permeabilised and intracellular staining was performed with Alexa647-conjugated anti-HA mAb (1:50) as described in the flow cytometry protocol section. Samples were acquired on CyAn™ ADP analyser (Beckman Coulter). Raw data was analysed in FlowJo and MFI of UCHT1-Fab' binding in the TCRβ-HA positive population extracted as excel files. Further analysis and statistical tests were performed in Prism (GraphPad Software). MFI of UCHT1-Fab' binding after background subtraction was plotted against the concentration of UCHT1-Fab' and fitted with non-linear regression (One site - Specific binding).

pMHC-tetramer binding to cells for association and dissociation analysis

To measure the relative association rate of pMHC tetramers to WT and βA291 TCR, J76 $CD8^+$ 1G4 WT or βA291 cells were induced for TCR expression with 1 μg/ml doxycycline for 48 - 72 h. TCR-deficient J76 cells were used to evaluate binding background. Cells were labelled with 200 nM CellTrace violet (Thermo Fisher) or with carrier control DMSO (Sigma), mixed 1:1 and washed once before being re-suspended at 20 x $10^6$/ml in RPMI, pre-warmed at 37 $^oC$. Cells were then distributed at 0.5 x $10^6$ cells, 25 μl/well in a 96-well V-bottom plate (Eppendorf) and equilibrated at 37 $^oC$ for 10 min in a ThermoMixer C (Eppendorf). 1:2 dilution series of 2X $(6V\text{-}A2)_4$-PE were added directly to the wells to a final concentration of 400 to 3.125 nM as indicated in the plot. After 10 min binding, a 3-fold excess of fixation buffer (BD Cytofix) was added and samples fixed for 10 min at 37 $^oC$. Samples were permeabilised and stained intracellularly with anti-HA (1:50) to detect total TCRβ as described in Flow cytometry protocols. To measure the relative dissociation rate of pMHC tetramers from WT and βA291 TCR, the same procedure as for on-rate measurements was used. This included cells, CellTrace violet labelling procedure, binding conditions and staining procedure, except that the assay was performed in 1.5 ml Eppendorf tubes. The concentration of $(6V\text{-}A2)_4$-PE in this assay was 50 nM. After 10 min of pMHC tetramer binding to the cells at 37 $^oC$, samples were washed once in RPMI containing 10 % FBS and re-suspended at $10^6$/ml in RPMI containing 10 % FBS supplemented with 10 μg/ml anti-HLA-A, B, C (BioLegend) to prevent rebinding of dissociated pMHC tetramer. Samples (0.2 x $10^6$ cells in 200 μl) were taken at the indicated times, washed once in FACS buffer and fixed for 2 h at 4 $^oC$ in fixation buffer (BD Cytofix). Cells were permeabilised and stained for TCRβ-HA as for the binding assay. Samples were acquired on a CyAn™ ADP. Raw FACS data was analysed in FlowJo (FlowJo Software) and MFI of $(6V\text{-}A2)_4$-PE binding and TCRβ-HA were extracted as excel files. Further analysis and statistical tests were performed in Prism (GraphPad Software). The MFI of the $(6V\text{-}A2)_4$-PE binding was corrected for background, normalized for maximal binding (max. binding = 100 %), plotted against the concentration of $(6V\text{-}A2)_4$-PE or against the time after initial binding and the curves were fitted by non-linear regression (Specific binding with Hill slope or Dissociation - One phase exponential decay).

TCR-CD3 expression efficiency

To evaluate the efficiency of TCR-CD3 surface expression, J76 $CD8^-$ 1G4 (or 2H5) WT or β mutant cells were induced for TCR expression with 1 μg/ml doxycycline overnight at 0.3 x $10^6$ cells/ml. Cells were incubated with 50 nM CellTrace violet or carrier control DMSO (Sigma), mixed in a 1:1 ratio and stained for surface CD3 and intracellular TCRβ-HA, using UCHT1 AF647-conjugated Fab' and AF488-conjugated anti-HA mAb respectively, as described in Flow cytometry procedures. J76 $CD8^-$ 1G4 (or 2H5) WT or β mutants, non-induced with doxycycline were used to evaluate binding background. Cells were analysed on a BD LSR Fortessa X20 flow cytometer (BD Biosciences) and acquired data was analysed with FlowJo FACS analysis software V10.0 (Tree Star, BD). 2D plots of TCRβ-HA vs. CD3 were obtained and gates for different levels of TCRβ-HA (low, medium and high) were applied. The

MFI ± SD of CD3 and TCRβ-HA was extracted within each gate and normalised to the WT in the corresponding gate. Statistical analysis using unpaired t-test was performed using Prism (GraphPad Software). MFI ± SD of TCRβ-HA for 1G4 WT and β mutants in those bins was not statistically significant (ns). The same procedure was used to evaluate the efficiency of TCR-CD3 surface expression in J76 CD8$^+$ 1G4 ζKO expressing ζWT and ζ mutants (ζA38, ζA41).

pMHC-tetramer stimulation for pErk or pζ-dose-response

To measure proximal signalling in response to stimulation with pMHC, J76 CD8$^+$ or CD8$^-$ 1G4 (or 2H5) WT or mutant cells were induced for TCR expression with doxycycline (dox) overnight at 0.3 x $10^6$ cells/ml. In all pMHC-tetramer stimulations for pErk, J76 CD8$^+$ or CD8$^-$ 1G4 WT (or 2H5 WT) were induced for TCR expression with 0.8 - 0.2 μg/ml of dox whereas β (or ζ) mutants were induced with 1 μg/ml. Different doses of doxycycline for 1G4 WT (or 2H5 WT) and β (or ζ) mutants were used in order to achieve same TCR-CD3 surface expression and to simplify computation of signalling outputs and eliminate potential errors arising from normalising for unequal $(6V\text{-}A2)_4$ binding. Cells were then labelled with 50 nM CellTrace violet (Thermo Fisher) or with carrier control DMSO (Sigma). CellTrace labelling did not interfere with proximal signalling events. To measure TCR surface expression a surface staining with AF647-conjugated UCHT1-Fab' fragment was performed according to the protocol described in Flow cytometry procedures before the stimulation assay. For this, labelled and unlabelled cells were mixed 1:1 and washed once before being re-suspended at 20 x $10^6$/ml in RPMI without FBS pre-warmed at 37 $^o$C. Cells were then distributed at 0.5 x $10^6$ cells/well (25 μl) in a 96-well V-bottom plate (Eppendorf) and equilibrated at 37 $^o$C for 10 min in a ThermoMixer C (Eppendorf). 1:2 dilution series of 2X $(6V\text{-}A2)_4$-PE or $(MART\text{-}1\text{-}A2)_4$-PE were added directly to the wells to a final concentration of 600 to 0.78 nM and 50 to 0.05 nM, respectively. After stimulation (pζ: 60 sec, pErk: 180 sec) an excess of fixation buffer (BD Cytofix) was added and samples fixed for 10 min at 37 $^o$C. Samples were permeabilised and stained intracellularly with AF647-conjugated anti-pErk (1:50) or with AF647-conjugated anti-pY142ζ (1:5) as described in Flow cytometry procedures. Samples were acquired on a CyAn™ ADP (Beckman Coulter) or BD LSR Fortessa X20 (BD Bioscience) as specified. Raw data was analysed by FlowJo (FlowJo software, part of BD) and MFIs were extracted from FlowJo as excel files. Further analysis and statistical tests were performed with Prism (GraphPad Software). For pζ-dose-response the background subtracted MFIs of pζ were plotted against the background subtracted MFI of tetramer $(6V\text{-}A2)_4$-PE binding and fitted by non-linear regression ([Agonist] vs. response - three parameters). The background subtracted values of MFIs of pζ with the highest dose of $(6V\text{-}A2)_4$-PE (400 nM) were normalized to WT or left as pairs and tested for significance with a t-test (paired or unpaired respectively). For pErk-dose-responses, pErk MFIs were background subtracted and plotted against the dose of $(6V\text{-}A2)_4$-PE or $(MART\text{-}1\text{-}A2)_4$-PE and fitted by non-linear regression ([Agonist] vs. response - Variable slope (four parameters) or [Agonist] vs. response - three parameters) or fitted to a minimal model of kinetic proofreading signalling (see Mathematical modelling section). In both non-linear regression fits, no constrains were applied and the expected best-fit values of maximum pErk for WT and mutant were analysed for significance with the F-test. The same procedure was used to evaluate max. pErk response upon $(6V\text{-}A2)_4$-PE titration, in J76 CD8$^+$ 1G4 ζKO expressing ζWT and ζmutants (ζA38, ζA41).

PMA stimulation

To evaluate maximal pErk potential of J76 $CD8^-$ 1G4 WT or βA291, cells were labelled with 50 nM CellTrace violet (Thermo Fisher) or with carrier control DMSO (Sigma), mixed 1:1 and washed once before being re-suspended at 20 x $10^6$/ml in RPMI without FBS pre-warmed at 37 $^0$C. Cells were then distributed at 0.5 x $10^6$ cells/well (25 µl) in a 96-well V-bottom plate (Eppendorf) and equilibrated at 37 $^0$C for 10 min in a ThermoMixer C (Eppendorf). 1:2 dilution of 2X PMA (Phorbol 12-myristate 13-acetate, Sigma) and Ionomycin (Sigma) was added directly to the wells to a final concentration of 66.6 ng/ml and 2 µg/ml, respectively. After stimulation (180 sec) an excess of fixation buffer (BD Cytofix) was added and samples fixed for 10 min at 37 $^0$C. Samples were permeabilised and stained intracellularly with AF647-conjugated anti-pErk (1:50) according to the protocol described above. Raw data was analysed by FlowJo (FlowJo software, part of BD) and MFIs were extracted from FlowJo as excel files. Further analysis and statistical test were performed with Prism (GraphPad Software). The background subtracted values of MFIs of pErk for WT and βA291 were exported and mean values of each experiment, measured in triplicates, were normalised to WT. Statistical significance was tested by unpaired t-test using Prism (GraphPad Software).

Basal ζ-phosphorylation

J76 $CD8^+$ 1G4 WT or βA291 cells were induced for TCR expression with 1 µg/ml doxycycline at 0.3 x $10^6$ cells/ml. Cells were harvested at 24, 48, 72 or 96 h and processed immediately for FACS analysis of CD3 surface expression or intracellular pζ according to the protocol described in Flow cytometry procedures. As staining for CD3 surface expression with an anti-CD3ε induced signalling even when carried out at 4 $^0$C using precooled solutions, samples were split in two and one part was stained for surface CD3 at 4 $^0$C while the other part was fixed at 37 $^0$C, permeabilised and analysed for basal pζ. Samples were acquired on a CyAn™ ADP (Beckman Coulter). Raw data was analysed by FlowJo (FlowJo software, part of BD) and MFIs were extracted from FlowJo as excel files. Further analysis and statistical tests were performed with Prism (GraphPad Software). pζ MFI was normalised to CD3 MFI (pζ/CD3), mean values of each experiment were normalised to WT or left as pairs and significance was tested by t-test (unpaired or paired respectively).

Sm-pMHC binding to cells

To detect non-specific adsorption of sm-pMHC onto J76 cell membrane during the stimulation assay (see soluble-monovalent pMHC stimulation section) CD8-deficient wtc51 J76 were induced or not for TCR expression with 1 µg/ml doxycycline for 48 h, labelled with 50 nM CellTrace violet (Thermo Fisher) or with carrier control DMSO (Sigma), mixed 1:1 and washed once before being re-suspended at 10 x $10^6$ in 125 µl of pre-warmed RPMI at 37 $^0$C. Cells were then distributed in a 96-well V-bottom plate (Eppendorf) and equilibrated at 37 $^0$C for 10 min in a ThermoMixer C (Eppendorf) under constant shaking (500 rpm). Dilution series of 2X sm-9V-A2 were added directly to the wells to a final concentration of 400 to 0.2 nM as indicated in the plot. After 5 min of binding, samples were rapidly centrifuged at 4 $^0$C, washed with 100 µl of ice-cold FACS-buffer and stained with APC-conjugated anti-HLA-A2 (1:100) for 20 min on ice to detect sm-9V-A2 bound to cells as described in Flow cytometry protocols. Samples were acquired on BD LSR Fortessa X20 (BD Bioscience). Raw FACS data for TCR-deficient and TCR-efficient J76 were analysed in FlowJo (FlowJo Software) and MFI of APC-anti-HLA-A2 were extracted as excel files. Further analysis and statistical tests were performed in Prism (GraphPad Software). The MFI of the APC-anti-HLA-A2 binding was corrected for background, plotted against the concentration of sm-9V-A2 and the curves were fitted by non-linear regression.

**Mathematical modelling**

To fit the tetramer binding data, we used an effective 1:1 binding model:

$$C = \mathrm{B_{max}} * \frac{[\mathrm{Tetramer}]}{(K_\mathrm{d} + [\mathrm{Tetramer}])}$$

where $B_{max}$ is the maximum binding and $K_d$ is an effective binding constant. We found that this model was able to fit both the WT and mutant TCR data. Importantly, we found that a single value of $K_d$ was sufficient to fit both dataset or equivalently, we found no evidence to reject the null hypothesis that tetramer binding was identical to both (F-test, $p$ = 0.066).

To fit the pErk data, we coupled the binding model to a minimal model of kinetic proofreading signalling (Dushek et al., 2011; McKeithan, 1995)

$$Y = \frac{[Tetramer]}{K_\mathrm{d} + [\mathrm{Tetramer}]} * \frac{k_\mathrm{p}}{(k_\mathrm{p} + k_\mathrm{off})^N}$$

where the first term determines tetramer occupancy (fraction between 0 and 1) and the second term determines the probability of signalling, $k_p$ the forward rate of proofreading, N the number of steps, and $k_{off}$ the unbinding rate.

Given that we found no evidence for a difference in binding (see above), we asked whether a difference in the proofreading rate ($k$p) can explain the difference in the pErk data. To do this, we simultaneously fit both the WT and mutant TCR pErk data to the mathematical model with a different value of $k_p$ for each TCR. As above, we fit a single value of $K_d$ for both datasets and in this model, we fixed the value of $k_{off}$ to the experimentally determined value (0.85 $s^{-1}$, (Aleksic et al., 2010) and fixed the value of N to the recently reported value (Tischer and Weiner, 2019; Yousefi et al., 2019). This is reasonable since we do not expect the TCR mutation to alter the number of steps.

We found that this model was able to produce an excellent fit to the data. Importantly, we found that a single value of $k_p$ could not explain the pErk data or equivalently, we had sufficient evidence to reject the null hypothesis that a single value of $k_p$ could explain the data (F-test, $p < 0.0001$).

**Biochemical analysis of the TCR-CD3 complex**

SDS-PAGE, immunoblotting and quantitation

SDS-polyacrylamide gel electrophoresis (PAGE) was performed using the protein electrophoresis system from Bio-Rad (Bio-Rad) according to the manufacturer's instructions. Custom-made 15 % polyacrylamide gels were used and proteins separated at 100 V in TGS (Tris Glycine SDS) running buffer (Bio-Rad). Separated proteins were transferred onto nitrocellulose membranes (Trans-blot Turbo Transfer Pack, Bio-Rad) using the Trans-blot turbo transfer system (Bio-Rad). As a routine, the High-MW protocol (25 V, 2.5 A, 10 min) or the Standard protocol (25 V, 1 A, 30 min) were used for the transfer. Membranes were saturated in blocking buffer (TBS, 0.1 % Tween-20, 3 % BSA) for 30 - 60 min at RT with gentle shaking and incubated overnight at 4 $^{o}$C or 1 h at RT with the primary Ab diluted (see **Key Resources Table** for a list of the Abs used in this study) in blocking buffer. After 3 washes of 10 min each with wash buffer (TBS-T), membranes were incubated with IRDye 800 CW or IRDye 680 CW (LI-COR) secondary Ab in blocking buffer for 45 min at RT in the dark. The membrane was then washed twice for 10 min with wash buffer (TBS-T) and once with wash buffer without Tween-20 in order to remove residual detergent and reduce the background during the acquisition. Near-Infrared (NIR) Western Blot Quantitative Detection was performed using the Odyssey CLx system (LI-COR) and the images were quantified using the Image Studio Lite software. The signal of each band was calculated as median - local background, intended as the signal detected in the area surrounding the band analysed. The signal values were exported as excel files for relative quantification.

DDM TCR-CD3 stability assay

Cells (10 x $10^6$/sample) were counted, centrifuged once at 425 x g, transferred to a 1.5 ml tube with RPMI 0 % FBS and washed with cold PBS. The cell pellet was lysed with 150 μl of ice-cold lysis buffer (150 mM NaCl, 20 mM Tris, pH 8.0 containing 0.5 or the indicated % of n-Dodecyl β-D-maltoside (DDM) (Millipore), 1X protease inhibitors (cOmplete, EDTA free, Roche), 1 mM Sodium Orthovanadate (NEB), 10 mM Sodium Fluoride (Sigma) and 25 U/ml Benzonase Nuclease (Millipore) and incubated 30 min on ice. 0.5 % DDM was chosen as it is the highest concentration of detergent compatible with maximal recovery of CD3 ε and ζ associated with αβ, in contrast, increasing concentration of DDM from 1 % to 4 % gradually decreased CD3 recovery, affecting ζ more than ε (Fig. **S2A** and data not shown). These data agreed with the recent cryo-EM structure (Dong et al., 2019) that shows ζζ to be the dimer most loosely associated to the rest of the complex. Lysates were centrifuged 10 min at 16,100 x g at 4 $^{o}$C and 15 μl of the post nuclear supernatant were saved for input control. The rest of the supernatant was transferred in a new tube containing 10 μl of anti-HA-conjugated agarose beads (Sigma) pre-washed 3 times with 1 ml of lysis buffer. One sample was transferred to a tube containing anti-HA-conjugated agarose beads pre-washed and pre-saturated for 2 h at 4 $^{o}$C under constant rotation with 50 μg/ml of HA peptide (Sigma), to assess background. Protein complexes were pulled down by incubating the supernatant and the anti-HA-conjugated agarose beads for 1 h at 4 $^{o}$C under constant rotation. Beads were centrifuged for 30 sec at 2,500 x g and washed 3 times with 1 ml of ice-cold lysis buffer. After the last wash, the supernatant was carefully removed and the beads re-suspended in 20 μl 1X NuPAGE LDS Sample Buffer (Invitrogen) containing or not (see below) 1X NuPAGE Sample Reducing Agent (Invitrogen) and incubated at 70 $^{o}$C for 10 min. After cooling, beads were centrifuged for 30 sec at 2,500 x g and the supernatant was collected in a fresh tube, loaded on a 15 % polyacrylamide gel at 4 $^{o}$C and processed as described above (SDS-PAGE, immunoblotting and quantitation). The results were analysed and presented as follow: for each sample, the background subtracted signals of ε, γ, δ and ζ were normalized to β and the obtained ratios (ε/β, γ/β, δ/β and ζ/β) were normalized to WT. Statistical unpaired t-test analysis was performed using Prism (GraphPad Software). To evaluate γε and δε recovery to αβ, the supernatant from a single pull down was split in two, loaded in duplicates on a 15 % polyacrylamide gel and immunoblotted for either β (HA), γ, ε and ζ or β (HA), δ, ε and ζ. Moreover, to minimise the interference of the IgG light chain of the anti-HA, used to pull down β, with the detection of δ and γ (molecular weight 20 - 25 kDa), at the end of the procedure described above, the beads were re-suspended in 20 μl 1X NuPAGE LDS Sample Buffer (Invitrogen) without adding reducing agent, but rather adding Iodoacetamide (Sigma) to a final concentration of 20 mM, an alkylation agent used to block thiols of proteins. Iodoacetamide was present in all the buffers used in this specific pull down (lysis buffer and sample buffer).

For endoglycosidase H (endo H) treatment, protein complexes were eluted by re-suspending the beads in 22 μl of lysis buffer containing 1X Glycoprotein Denaturing Buffer (NEB) and incubating at 100 $^{o}$C for 10 min. Beads were centrifuged for 30 sec at 2,500 x g, the supernatant was collected, divided in two fresh tubes and 1X buffer 3 (NEB) was added in presence or not of endo H (750 U) for 1 h at 37 $^{o}$C. Samples were then incubated at 70 $^{o}$C for 10 min with 1X NuPAGE LDS Sample Buffer (Invitrogen) containing 1X NuPAGE Sample Reducing Agent (Invitrogen) and loaded on a 15 % polyacrylamide gel at 4 $^{o}$C and processed as described above (SDS-PAGE, immunoblotting and quantitation).

<u>ζTST pull-down</u>

J76 CD8$^{+}$ 1G4 ζKO expressing inducible ζTST were treated with 1 μg/ml doxycycline overnight, counted and washed once with 150 μl of ice-cold PBS. Pellets of 10 x 10$^{6}$ cells were lysed with 150 μl ice-cold lysis buffer (150 mM NaCl, 20 mM Tris, pH 8.0 containing 0.5 % n-Dodecyl β-D-maltoside (DDM) (Millipore), 1X proteases inhibitors (cOmplete, EDTA free, Roche), 1 mM Sodium Orthovanadate (NEB), 10 mM Sodium Fluoride (Sigma) and 25 U/ml Benzonase Nuclease (Millipore) and incubated on ice for 10 min. Lysates were centrifuged for 10 min at 16,100 x g at 4 $^{o}$C and 15 μl of the post nuclear supernatant was collected as a control for total input. The rest of the supernatant was transferred into a fresh tube containing 10 μl Strep-Tactin Sepharose beads (IBA Lifesciences) pre-washed with lysis buffer and incubated for 30 min at 4 $^{o}$C under constant rotation. Alternatively, the supernatant was incubated 30 min at 4 $^{o}$C with anti-HA-conjugated agarose beads (Sigma). One sample was transferred to a tube containing Strep-Tactin beads or anti-HA-conjugated agarose beads pre-washed and pre-saturated for 2 h at 4 $^{o}$C under constant rotation with 50 mM biotin or 50 μg/ml of HA peptide respectively, to assess background. Beads were centrifuged for 30 sec at 2,500 x g and washed 3 times with 1 ml of ice-cold lysis buffer. After the last wash, the supernatant was carefully removed and the beads re-suspended in 20 μl 1X NuPAGE LDS Sample Buffer (Invitrogen) containing 1X NuPAGE Sample Reducing Agent (Invitrogen) and incubated at 70 $^{o}$C for 10 min. After cooling, beads were centrifuged for 30 sec at 2,500 x g and the supernatant was collected in a fresh tube before being separated on a 15 % polyacrylamide gel at 4 $^{o}$C. Quantitative immunoblotting with anti-HA and anti-ζ mAbs were used to identify the β isoform interacting with ζ among the three identified in the total β pull-down performed using anti-HA-conjugated agarose beads.

**Ligand-induced TCR-CD3 quaternary structure changes**

<u>Resting or pMHC-stimulated TCR-CD3</u>

The structural integrity of the TCR-CD3 complex upon $(9V\text{-}A2)_4$ and sm-9V-A2 stimulation of intact cells at physiological temperature or at 0 $^{o}$C was evaluated by the following assay. J76 CD8$^{+}$ or CD8$^{-}$ cells expressing inducible TCR wtc51 were treated with 1 μg/ml doxycycline overnight, counted and washed once with RPMI without FBS. Cells were re-suspended at 10 x 10$^{6}$/150 μl in the same medium and pre-incubated for 10 min at 37 $^{o}$C in a ThermoMixer C (Eppendorf) under constant shaking (500 rpm) or 20 min on ice. $(9V\text{-}A2)_4$ obtained using a His-tagged streptavidin (His-SA) (100 nM), sm-9V-A2 (200 nM) or RPMI alone was added to the cells for 5 min at 37 $^{o}$C or on ice. After ligand binding, the samples were immediately centrifuged 30 sec at 800 x g and rapidly washed once with 150 μl of ice-cold PBS. For Lck inhibition, cells were pre-treated with 5 μM A770041 (Axon) at 37 $^{0}$C for 20 min prior to stimulation and 5 μM A770041 was kept during the binding. Cell pellets were lysed with 150 μl ice-cold lysis buffer (300 mM NaCl, 20 mM Tris, pH 8.0 containing 0.5 % n-Dodecyl β-D-maltoside (DDM) (Millipore), 1X proteases inhibitors (cOmplete, EDTA free, Roche), 1 mM Sodium Orthovanadate (NEB), 10 mM Sodium Fluoride (Sigma), 25 U/ml Benzonase Nuclease (Millipore) and 10 mM Imidazole and incubated on ice for 5 min. Lysates were centrifuged for 5 min at 16,100 x g at 4 $^{0}$C and 15 μl of the post nuclear supernatant was collected as a control for total input. The rest of the supernatant was incubated 5 min on ice with 1 μg of His-tagged streptavidin (sm-9V-A2 engaged samples) or transferred directly into a fresh tube containing 10 μl HisPur Cobalt Resin (Thermo Fisher) ((9V-A2)$_4$ engaged samples) or 10 μl anti-HA-conjugated agarose beads (Sigma) (non-stimulated samples), pre-washed with lysis buffer and incubated for 15 min and 30 min respectively at 4 $^{0}$C under constant rotation. To assess background, one sample was transferred to HisPur Cobalt beads or anti-HA-conjugated agarose beads pre-washed and pre-saturated at 4 $^{0}$C for 2 h under constant rotation with 300 mM Imidazole or 50 μg/ml of HA peptide respectively. Beads were centrifuged for 30 sec at 2,500 x g and washed 3 times with 1 ml of ice-cold lysis buffer. After the last wash, the supernatant was carefully removed and the beads re-suspended in 20 μl 1X NuPAGE LDS Sample Buffer (Invitrogen) without reducing agent and incubated at 80 $^{0}$C for 5 min. After cooling, beads were centrifuged for 30 sec at 2,500 x g and the supernatant was collected in a fresh tube and incubated at 80 $^{0}$C for 5 min with 1X NuPAGE Sample Reducing Agent (Invitrogen). Samples were separated on a 15 % polyacrylamide gel at 4 $^{0}$C and processed as described above (SDS-PAGE, immunoblotting and quantitation). Quantitative immunoblotting with anti-HA and anti-ζ mAbs were used to evaluate the amount of β and ζ pulled-down in each condition. The amount of ζ extracted at steady state was normalised to a specific β isoform ($\beta_2$) that we proved to be the only one able to interact with the ζ chain (for details see in ζTST pull down) and that corresponds to the only detectable isoform in (9V-A2)$_4$-engaged receptor pull downs (for schematic of the procedure see Figs. **S6A** and **S7A**). The value for $\zeta/\beta_2$ ratio from non-stimulated samples was set equal to one. This value represented the recovery of intact TCR-CD3 complex and was compared to $\zeta/\beta_2$ from (9V-A2)$_4$ stimulated samples. Statistical analysis using unpaired t-test was performed using Prism (GraphPad Software).

To isolate sm-9V-A2-engaged wtc51 with monomeric avidin agarose beads (Thermo), cells were incubated with sm-9V-A2 (200 nM) or RPMI alone for 5 min at 37 $^{0}$C. After ligand binding, the samples were lysed as described above and the post-nuclear supernatant was incubated with monomeric avidin beads for 30 min at 4 $^{0}$C under constant rotation.

For primary CD8 T-cells expressing constitutive TCR wtc51, cells were incubated for 5 min at 37 $^{0}$C with (9V-A2)$_4$ obtained using a His-tagged streptavidin (His-SA) (40 nM), 9V-A2 (400 nM) or RPMI alone. Lysis was performed in presence of 30 mM imidazole and this concentration was maintained during the following washes.

Isolation of (6I-A2)$_4$ and sm-6I-A2 engaged receptors in J76 CD8$^+$ cells expressing 868 TCR ($K_d$ = 50 nM) was performed as described above with little variations. Cells were incubated with (6I-A2)$_4$ obtained using a His-tagged streptavidin (His-SA) (40 nM), sm-6I-A2 (400 nM) or RPMI alone for 5 min at 37 $^{0}$C. Lysis was performed in the presence of 30 mM imidazole and this concentration was maintained during the following washes.

To have a comparable amount of β detected by immunoblotting in non-stimulated samples and $(6I\text{-}A2)_4$ and sm-6I-A2 engaged samples, only 1/3 of the elution from non-stimulated samples was loaded on the gel.

Isolation of $(9V\text{-}A2)_4$ engaged receptors in J76 $CD8^+$ cells expressing inducible QM-α TCR ($K_d$ = 140 nM) was performed as described above with little variations. Cells were incubated with $(9V\text{-}A2)_4$ obtained using a His-tagged streptavidin (His-SA) (333 nM), or RPMI alone for 5 min at 37 $^oC$. Lysis was performed in the presence of 30 mM imidazole and this concentration was maintained during the following washes. Elution was performed by incubating the beads at 4 $^oC$ for 15 min with 15 μl of lysis buffer containing 300 mM imidazole. Beads were then centrifuged for 30 sec at 2500 x g and the supernatant was collected in a fresh tube and incubated at 80 $^oC$ for 5 min with 1X NuPAGE LDS Sample Buffer containing 1X NuPAGE Sample Reducing Agent (Invitrogen).

To have a comparable amount of β detected by immunoblotting in non-stimulated samples and $(9V\text{-}A2)_4$ engaged samples, only 1/40 of the elution from non-stimulated samples was loaded on the gel.

Soluble monovalent-pMHC stimulation

Intracellular triggering upon soluble monovalent agonist (sm-pMHC) binding was evaluated by pErk activation, according to the following procedure. J76 $CD8^+$ or $CD8^-$ cells expressing 868 or wtc51 TCRs (the latter induced for TCR-CD3 expression with 1 μg/ml doxycycline for 48 h) were counted and washed once with RPMI without FBS. Cells were re-suspended at 10 x $10^6$/125 μl in the same medium and pre-incubated for 10 min at 37 $^oC$ in a ThermoMixer C (Eppendorf) under constant shaking (500 rpm). For Lck inhibition, cells were pre-treated with 5 μM A770041 (Axon) at 37 $^oC$ for 15 min prior to stimulation and 5 μM A770041 was kept during the binding. Tetramers ($pMHC_4$,25 nM), sm-pMHC (100 nM) or RPMI alone were added to the cells for 5 min at 37 $^oC$. After ligand binding, samples were immediately boiled for 10 min at 95 $^oC$ by adding pre-warmed 2X NuPAGE LDS Sample Buffer (Invitrogen) containing 2X NuPAGE Sample Reducing Agent (Invitrogen) to instantly stop the reaction. Cell lysates were let cool down and 25 U of Benzonase Nuclease (Millipore) were added every 15 min for 4 times to allow a complete DNA/RNA digestion. Lysates were centrifuged for 15 min at 16,100 x g at 4 $^oC$ and supernatant was collected in a fresh tube before being separated on a 15 % polyacrylamide gel. Quantitative immunobloting with anti-pErk and anti-actin was used to evaluate the amount of Erk phosphorylation (for details, see SDS PAGE, immunoblotting and quantitation). The value for pErk (-background)/actin ratio from non-stimulated samples was set equal to one. Statistical analysis using unpaired t-test was performed using Prism (GraphPad Software).

To exclude that the observed signalling (pErk) was the consequence of surface cell-to-cell ligand cross-presentation, we used the above protocol (also used for the evaluation of sm-pMHC binding by FACS, see sm-pMHC binding to cells section) with cells expressing or not TCR. Briefly, CD8-deficient wtc51 J76 not induced for TCR expression were counted, washed once with RPMI, re-suspended at 10 x $10^6$/125 μl in the same medium and pre-incubated for 10 min at 37 $^oC$ in a ThermoMixer C (Eppendorf) under constant shaking (500 rpm). Sm-9V-A2 (200 nM) or RPMI alone was added to the cells for 5 min at 37 $^oC$, rapidly washed with ice-cold RPMI (to minimize unbinding), re-suspended in pre-warmed RPMI at 10 x $10^6$/125 μl and added to an equal amount of dox-induced CD8-deficient wtc51 J76 for 5 min at 37 $^oC$. After ligand binding cells were immediately boiled for 10 min at 95 $^oC$ by adding pre-warmed 2X NuPAGE LDS Sample Buffer (Invitrogen) containing 2X NuPAGE Sample

Reducing Agent (Invitrogen) to instantly stop the reaction and cell lysates were analised as described above.

UCHT1-Fab' or UCHT1 Ab-engaged TCR-CD3

The structural integrity of the TCR-CD3 complex upon agonist stimulation of intact cells at physiological temperature or at 0 $^{o}$C was evaluated by the following assay. The anti-CD3ε mAb UCHT1 was used as a potent agonist of TCR-CD3 and its Fab' as the non-agonist control that binds to the same determinant. J76 $CD8^{+}$ cells stably expressing 1G4 TCR were counted and washed once with RPMI without FBS. Cells were re-suspended at 10 x $10^{6}$/150 μl in the same medium and pre-incubated for 10 min at 37 $^{o}$C in a ThermoMixer C (Eppendorf) under constant shaking (500 rpm) or for 20 min on ice. Mono-biotinylated Fab' of UCHT1 mAb (0.72 μg) or biotinylated UCHT1 intact mAb (2 μg) (BioLegend) was added to the cells for 5 min at 37 $^{o}$C or on ice. These doses of anti-CD3ε ligands that corresponded to approximately the same molarity were proven to pull-down similar amounts of TCR-CD3 complex, as detected by ε immunoblot. One sample was left untreated as negative control. After ligand binding, the samples were immediately centrifuged 30 sec at 800 x g and rapidly washed once with 150 μl of ice-cold PBS. For Lck inhibition, cells were pre-treated with 5 μM A770041 (Axon) at 37 $^{o}$C for 20 min prior to binding of Fab' UCHT1 or UCHT1 mAb and 5 μM A770041 was kept during the binding. Cell pellets were lysed with 150 μl ice-cold lysis buffer (150 mM NaCl, 20 mM Tris, pH 8.0 containing 0.5 % n-Dodecyl β-D-maltoside (DDM) (Millipore), 1X proteases inhibitors (cOmplete, EDTA free, Roche), 1 mM Sodium Orthovanadate (NEB), 10 mM Sodium Fluoride (Sigma) and 25 U/ml Benzonase Nuclease (Millipore) and incubated on ice for 30 min. Lysates were centrifuged for 10 min at 16,100 x g at 4 $^{o}$C and 15 μl of the post nuclear supernatant was collected as a control for total input. The rest of the supernatant was transferred into a fresh tube containing 10 μl Streptavidin Sepharose High Performance beads (GE Healthcare) pre-washed with lysis buffer and incubated for 1 h at 4 $^{o}$C under constant rotation. Beads were centrifuged for 30 sec at 2,500 x g and washed 3 times with 1 ml of ice-cold lysis buffer. After the last wash, the supernatant was carefully removed and the beads re-suspended in 20 μl 1X NuPAGE LDS Sample Buffer (Invitrogen) containing 1X NuPAGE Sample Reducing Agent (Invitrogen) and incubated at 70 $^{o}$C for 10 min. After cooling, beads were centrifuged for 30 sec at 2,500 x g and the supernatant was collected in a fresh tube before being separated on a 15 % polyacrylamide gel. Quantitative immunobloting with anti-ε, anti-HA and anti-ζ mAbs was used to evaluate the amount of ε, β and ζ, respectively, pulled-down after each stimulatory condition (for details, see SDS PAGE, immunoblotting and quantitation). The recovery of ε was assumed to be invariant in both the stimulatory (UCHT1 mAb) and non-stimulatory (UCHT1-Fab') condition and was used for the normalisation of the amounts of the other subunits in each pull-down. Therefore, the values for β/ε and ζ/ε ratios obtained from UCHT1-Fab' engaged TCR-CD3 were set equal to one. These values represented the recovery of intact TCR-CD3 complex and were compared to the same ratios from UCHT1 stimulated samples. Statistical analysis using unpaired t-test was performed using Prism (GraphPad Software).

**Microscopy**

dSTORM imaging and analysis

Prior to dSTORM imaging, cells were plated on glass μ-slide 8-wells chamber (Ibidi) coated with recombinant human ICAM-1 protein (R&D Systems) at 2.5 μg/ml concentration. Cells were incubated for 15 minutes at 37 $^{o}$C followed by fixation for 30 min with 4 % PFA at RT. Cells were blocked with 5 % BSA in PBS for 1 h followed by incubation with an anti-CD3 primary antibody directly conjugated with Alexa-Fluor 647 (BioLegend) for 1 h at RT. Cells were washed three times with PBS and post-fixed with 4 % PFA for 5 min before imaging. Cysteamine based dSTORM imaging buffer was used to perform the single molecule localization experiments with the following composition: 100 mM Cysteamine MEA (Sigma Aldrich), 5 % Glucose (Sigma Aldrich), 1 % Glox (0.5 mg/mL glucose oxidase, 40 mg/mL catalase (Sigma Aldrich) dissolved in 1X PBS. dSTORM imaging of CD3 was performed by using a 150 × 1.45-NA oil-immersion objective in TIRF mode (TIRF; Olympus). Laser light of 642 nm was used to excite the Alexa-Fluor 647 dye and switch it to the dark state. An additional 405 nm laser light was used to reactivate the Alexa-Fluor 647 fluorescence. The emitted light from Alexa-Fluor 647 was collected by the same objective and imaged by an EMCCD camera (Evolve Delta; Photometrics) at a frame rate of 10 ms per frame. A maximum of 5,000 frames per condition were acquired. dSTORM images were analysed and rendered as previously described (Bates et al., 2007; Huang et al., 2008) using a custom-written software (Insight3, provided by B. Huang, University of California, San Francisco). Peaks in single-molecule images were identified based on a threshold and fit to a simple Gaussian to determine the x and y positions. Only localizations with photon count > 400 photons were included, and localizations that appeared within one pixel in five consecutive frames were merged together and fitted as one localization. The final images were rendered by representing the x and y positions of the localizations as a Gaussian with a width that corresponds to the determined localization precision. Sample drift during acquisition was calculated and subtracted by reconstructing dSTORM images from subsets of frames (500 frames) and correlating these images to a reference frame (the initial time segment).

Pair auto-correlation analysis

Pair auto-correlation analysis is independent of the number of localizations and is not susceptible to over-counting artefacts related to fluorescent dye re-blinking (Stone et al., 2017). Auto-correlation analysis of CD3 protein was performed using MATLAB software provided by Sarah Shelby and Sarah Veatch from University of Michigan. Regions containing cells were masked by a region of interest, and the auto-correlation function from the x and y coordinate list from the 642 nm dSTORM channel was computed from these regions using an algorithm described previously (Shelby et al., 2013; Stone et al., 2017; Veatch et al., 2012).

DBSCAN cluster analysis

Quantitative cluster analysis was based on Density-based spatial clustering of applications with noise (DBSCAN) (Ester, 1996). The DBSCAN method detects clusters using a propagative method which links proteins belonging to the same cluster based on the minimum number of neighbours $\varepsilon$ ($\varepsilon = 7$) in the radius r (r = 25 nm). Clus-DoC (Pageon et al., 2016) MATLAB based software with implemented DBSCAN analysis was used. The x and y coordinate list of dSTORM localizations was used and regions containing cells were selected to give the mean±SD value of CD3 cluster size per cell.

Fluorescent Recovery after Photobleaching (FRAP) and analysis

Prior to FRAP experiment, cells were incubated for 20 minutes at 37 $^{0}$C with an anti-CD3 primary antibody directly conjugated with Alexa-Fluor 647 (BioLegend). Cells were washed with cell media and seeded on glass μ-slide 8 well chamber (Ibidi) coated with Poly-L-Lysine for 5 min before imaging. Cysteamine based dSTORM imaging buffer was used to perform the experiments (see dSTORM imaging and analysis). FRAP imaging of CD3 was performed by using a Nikon A1R HD25 confocal system with a 60x oil-immersion objective (Nikon, UK) in humidified 37 $^{0}$C, 5 % $CO_2$ chamber. A circular region was defined on the surface of the cell. This region was bleached using the 405 nm laser with maximum power for 7 sec. Before and after bleaching, the whole region was visualized by 640 nm laser for the duration of 50 s in 4 sec intervals. FRAP curves were exported from NIS-Elements software (Nikon, UK) and the post processing of the extracted curves was performed in Origin software (OriginPro 2017; OriginLab).

**Atomistic Molecular Dynamics Simulations**

The transmembrane region (TMR) of the T-cell receptor (TCR) was extracted from the cryo-EM structure PDB: 6JXR (Dong et al., 2019) and used for atomistic molecular dynamics simulations. The residues of TCRβ TMR are numbered according to that used in our experiments ("n - 1" compared to PDB: 6JXR, i.e. our βY291 is βY292 in 6JXR). The TMR sequences used are shown in **Table S3**. The octameric protein structure was inserted in a complex asymmetric bilayer (lipid concentration is shown in **Table S4**) using CHARMM-GUI (charmm-gui.org) and was solvated with TIP3 water molecules. The lipid types and their concentrations in our simulated bilayer were considered from studies on the T-cell membrane conducted by Zech *et al*. (Zech et al., 2009). The system was then neutralized with a concentration of 150 mM of NaCl ions. The resultant molecular system and molecular dynamics parameters were obtained from CHARMM-GUI and thereafter run using Gromacs 2016.4 (Van Der Spoel et al., 2005) using the CHARMM36 forcefield (Lee et al., 2016). An energy-minimization of the entire system was conducted for 5000 steps using the steepest descent algorithm. The energy-minimized system underwent a 6-step NPT equilibration where position restraints on the protein backbone, side-chain and lipids were gradually released. The LINCS algorithm (Hess et al., 1997) was used to apply constraints on bond lengths. The fully equilibrated system was used to generate starting points for 3 production simulation repeats with different initial velocities. This was done for the wild-type complex, ζA38, ζA41 and βA291 mutants which started from the same initial configuration. Each production simulation was run for 1250 nanoseconds employing a 2 femtoseconds time-step and the co-ordinates were saved every 40 picoseconds. The Nose-Hoover thermostat (Hoover, 1985) was used with a reference temperature of 323 K and coupling constant of 1 picosecond. The Parrinello-Rahman barostat (Parrinello and Rahman, 1981) was used with a semi-isotropic pressure coupling type, reference pressure of 1 atmosphere, compressibility value of 4.5 x $10^{-5}$ $bar^{-1}$ and coupling constant of 5 picoseconds. The Particle mesh Ewald algorithm (Essmann et al., 1995) with a 12 Å distance cut-off was used to define non-bonded van der Waals (Lennard-Jones) and coulombic interactions. Visualisation was carried out using VMD (Humphrey et al., 1996). Protein-protein interaction profiles and spatial distribution plots are a result of merged data from all the 3 repeats of all-atom simulations. To calculate the spatial distribution of ζζ relative to αβ, the positions of Cα atoms of αβ were fixed throughout the simulation time. The spatial distributions of Cα atoms of ζζ and αβ are viewed from the extracellular region.

**Modelling of the 1G4 TCR affinity mutants**

The 1G4-A2-SLL structure (from PDB: 2BNR) (Chen et al., 2005) was used to model the 1G4 QM-α-A2-SLL and 1G4 wtc51-A2-SLL tri-molecular complex structures. Sequences were adjusted with COOT (Emsley and Cowtan, 2004) and graphical representations and analysis were prepared with PYMOL (DeLano WL. The PyMOL Molecular Graphics System. Schrödinger LLC. 2002; Version 1.: http://www.pymol.org. Available: http://www.pymol.org).

#### QUANTIFICATIONS AND STATISTICAL ANALYSIS

##### Flow cytometry

All experiments were replicated at least two - three times with technical and/ or biological replicates each time. Raw data was processed in FlowJo (FlowJo software, part of BD) and MFIs, counts or percentages were extracted from FlowJo as excel files. Statistical analysis and non-linear regression fitting were performed with Prism (GraphPad Software). Significance was assessed by two-tailed paired or unpaired t-test. Non-linear regressions were tested for Goodness of Fit ($R^2$), for normality (D'Agostino & Pearson omnibus normality test) and with the replicates test. Non-linear regression fits are specified in the methods paragraph describing the experimental procedure. $R^2$ and *p*-value for the fits, significance tests and significance are indicated in the figure or figure legend.

##### Pull-down and Immunoblotting

Experiments were replicated at least two - three times with technical and/ or biological replicates each time as indicated at the representing figure. Raw data was extracted from Image studio (LI-COR Biosciences) and processed in Excel and Prism (GraphPad Software). Statistical analysis (two-tailed unpaired t-test) was performed and method and significance are indicated in the figure or figure legend.

##### Microscopy

Samples were tested for normality with a Kolmogorov–Smirnov test. The statistical significance of differences between two datasets was assessed by a two-tailed t-test assuming unequal variance. All statistical analysis was performed using Origin software (OriginPro 2017; OriginLab).

##### Simulations

For the atomistic simulations multiple repeat simulations were performed to ensure reproducibility of our results. Analyses of the simulations were conducted using Gromacs, VMD, and locally written code.

##### Modelling

Graphics and analysis were prepared with PYMOL (DeLano WL. The PyMOL Molecular Graphics System. Schrödinger LLC 2002; Version 1.: http://www.pymol.org. Available: http://www.pymol.org).

**Significance**

* $p \leq 0.05$ /** $p \leq 0.01$/*** $p \leq 0.001$/**** $p \leq 0.0001$/ ns = not significant.

##### Table S1. 1G4 amino acid sequence

| 1G4 WT αβ self-cleavable single polypeptide (TCRβ-spacer-HA-spacer-F2A-L1-TCRα-L2-Flag) |
|---|
| MSIGLLCCAALSLLWAGPVNAGVTQTPKFQVLKTGQSMTLQCAQDMNHEYMSWYRQDPGMGLRLIHYSVGAGITDQ<br>GEVPNGYNVSRSTTEDFPLRLLSAAPSQTSVYFCASSYVGNTGELFFGEGSRLTVLEDLKNVFPPEVAVFEPSEAEISHTQKA<br>TLVCLATGFYPDHVELSWWVNGKEVHSGVSTDPQPLKEQPALNDSRYCLSSRLRVSATFWQNPRNHFRCQVQFYGLSEN<br>DEWTQDRAKPVTQIVSAEAWGRADCGFTSESYQQGVLSATILYEILLGKATLYAVLVSALVLMAMVKRKDFSRGGSGGGS<br>GGGSGGGSGGGSGASYPYDVPDYAGAGGSGGGSGGGSGGGSGGGSGVKQTLNFDLLKLAGDVESNPGPEFMETLLGL<br>LILWLQLQWVSSKQEVTQIPAALSVPEGENLVLNCSFTDSAIYNLQWFRQDPGKGLTSLLLIQSSQREQTSGRLNASLDKSS<br>GRSTLYIAASQPGDSATYLCAVRPTSGGSYIPTFGRGTSLIVHPYIQNPDPAVYQLRDSKSSDKSVCLFTDFDSQTNVSQSKD<br>SDVYITDKTVLDMRSMDFKSNSAVAWSNKSDFACANAFNNSIIPEDTFFPSPESSCDVKLVEKSFETDTNLNFQNLSVIGFR<br>ILLLKVAGFNLLMTLRLWSSSGDYKDDDDK |

#### Table S2. gRNA and Primers

| | |
|---|---|
| CRISPR gRNA | |
| CD3ζ | CAGGCACAGTTGCCGATTACAGG |
| Gateway cloning | |
| 1G4 β-HA-F2A-α-FLAG FW | GGGGACAAGTTTGTACAAAAAAGCAGGCTTAATGAGCATCGGCCTCCTGTG CTGTGCAGCC |
| 1G4 β-HA-F2A-α-FLAG REV | GGGGACCACTTTGTACAAGAAAGCTGGGTTTTATTTGTCGTCGTC GTCTTTGTAGTCTCC |
| CD3ζ-TST FW | ACAAGTTTGTACAAAAAAGCAGGCTTCACCATGAAGTGGAAGGCGCTTTTC AC |
| CD3ζ-TST REV | ACCACTTTGTACAAGAAAGCTGGGTTTTATTTTTCGAACTGCGGGTGGCTC |
| Amplification of 1G4 WT from cDNA | |
| TCRα FW | GGATCCATGGAGACCCTCTTGGGCCTGCTT |
| TCRα REV | TCTAGAGCTGGACCACAGCCGCAGCGT |
| TCRβ FW | GGATCCATGAGCATCGGCCTCCTGTGCTGT |
| TCRβ REV | GGATCCATGAGCATCGGCCTCCTGTGCTGT |

##### Table S3. Protein sequences used in the all atom simulations

| Chain | Residue range | TMR sequences used |
|---|---|---|
| CD3-$\zeta_1$ | 27-57 | LDPKLCYLLDGILFIYGVILTALFLRVKFSR |
| CD3-$\zeta_2$ | 27-55 | LDPKLCYLLDGILFIYGVILTALFLRVKF |
| TCR-$\alpha$ | 239-273 | DTNLNFQNLSVIGFRILLLKVAGFNLLMTLRLWSS |
| TCR-$\beta$ | 266-307 | TSESYQQGVLSATILYEILLGKATLYAVLVSALVLMAMVKRK |
| CD3-$\delta$ | 98-129 | ELDPATVAGIIVTDVIATLLLALGVFCFAGHE |
| CD3-$\varepsilon$ ($\delta\varepsilon$ dimer) | 125-155 | MDVMSVATIVIVDICITGGLLLLVYYWSKNR |
| CD3-$\gamma$ | 109-138 | ELNAATISGFLFAEIVSIFVLAVGVYFIAG |
| CD3-$\varepsilon$ ($\gamma\varepsilon$ dimer) | 125-156 | MDVMSVATIVIVDICITGGLLLLVYYWSKNRK |

##### Table S4. Bilayer lipid concentrations used in the all atom simulations

| Lipid concentration (%) | POPC | POPS | POPE | SM | CHOL | $PIP_2$ | $PIP_3$ |
|---|---|---|---|---|---|---|---|
| Inner leaflet | 10 | 20 | 40 | - | 20 | 8 | 2 |
| Outer leaflet | 50 | - | 10 | 20 | 20 | - | - |

#### KEY RESOURCES TABLE

| REAGENT or RESOURCE | SOURCE | IDENTIFIER |
|---|---|---|
| Antibodies | | |
| Mouse anti-human AF647 CD3ζ pY142 mAb | BD Bioscience | Cat # 558489; RRID: |
| Mouse anti-human PE CD3ζ pY142 mAb | BD Bioscience | Cat # 558488; RRID: |
| Mouse anti-human AF647 CD3ε mAb | BioLegend | Cat # 300422; RRID: |
| Mouse anti-human BV421 CD3ε mAb | BioLegend | Cat # 300434; RRID: |
| Mouse anti-human PE CD3ε mAb | BioLegend | Cat # 300456; RRID: |
| Mouse biotinylated anti-human CD3ε mAb | BioLegend | Cat # 300404; RRID: |
| Rabbit anti-human AF647 pErk mAb | Cell Signaling Technology | Cat # 4375; RRID: |
| Rabbit anti-pTyr416 Src polyclonal Ab | Cell Signaling Technology | Cat # 2101; RRID: |
| Rabbit anti-human ERK 1/2 polyclonal Ab | Cell Signaling Technology | Cat # 9107; RRID: |
| Rabbit anti-HA mAb | Cell Signaling Technology | Cat # 3724; RRID: |
| Rabbit anti-human CD3 γ mAb | Abcam | Cat # 134096; RRID: |
| Rabbit anti-human CD3 δ polyclonal Ab | GeneTex | Cat # 105811; RRID: |
| Mouse AF488 anti-HA mAb | Cell Signaling Technology | Cat # 2350; RRID: |
| Mouse AF647 anti-HA mAb | Cell Signaling Technology | Cat # 3444; RRID: |
| Rat anti-human CD3ε mAb | Cell Signaling Technology | Cat # 4443; RRID: |
| Mouse anti-CD3ζ mAb | Santa Cruz Biotechnology | Cat # sc-1239; RRID: |
| Rabbit anti-CD3ζ pY142 mAb | Abcam | Cat # ab68235; RRID: |
| Mouse anti-Actin mAb | Merck Millipore | Cat # MAB1501; RRID: |
| Mouse anti-pTyr mAb | Merck Millipore | Cat # 05-321; RRID: |
| Mouse anti-human CD3ε mAb | BioXCell | Cat # BE0231; RRID: |
| Mouse anti-HLA-A, B, C mAb | BioLegend | Cat # 311423; RRID: |
| Mouse anti-human APC HLA-A2 mAb | BioLegend | Cat # 343308; RRID: |
| Rabbit anti-CD3ζ polyclonal Ab | See citation Ref. (San Jose et al., 1998) | N/A |
| Bacterial and Virus Strains | | |
| *E.coli* DH5α Competent Cells | Thermo Fisher | Cat # 18265-017; RRID: |
| *E.coli* Stbl3 Competent Cells | Invitrogen | Cat # C7373-03; RRID: |
| Chemicals, Peptides, and Recombinant Proteins | | |
| 10X Tris/Glycine/SDS | Biorad | Cat # 161-0772; RRID: |
| 16 % Formaldehyde (w/v), Methanol-free | Thermo Fisher | Cat #: 28908; RRID: |
| 2-Mercaptoethylamine-*HCl* | Thermo Fisher | Cat # 20408; RRID: |
| Annexin-V-AF647 | BioLegend | Cat # 640912; RRID: |
| Annexin-V-PE | BioLegend | Cat # 640908; RRID: |
| Alexa-Fluor 488 $C_5$ Maleimide | Thermo Fisher | Cat # A10254; RRID: |
| Alexa-Fluor 647 $C_2$ Maleimide | Thermo Fisher | Cat # A20347; RRID: |
| Amicon Ultra 15 ml centrifugal filter units | Millipore | Cat # UFC901008; RRID: |
| Anti-HA-conjugated agarose beads | Sigma Aldrich | Cat # AL2095; RRID: |
| A770041 Lck inhibitor | Axon Medchem | Cat # 1698; RRID: |
| BamH1 | New England Biolabs | Cat # R0136; RRID: |
| BD Cytofix buffer | BD Biosciences | Cat # 554655; RRID: |
| BD PhosFlow Perm/Wash buffer 1 | BD Biosciences | Cat # 55785; RRID: |
| Benzonase Nuclease | Millipore | Cat # 71206-25KUN; RRID: |
| BirA kit | Avidity | Cat # BirA500; RRID: |
| Bovine serum albumin (BSA) | Sigma Aldrich | Cat # A4503-100G; RRID: |
| Bug Buster protein extraction reagent | Millipore | Cat # 70584-4; RRID |
| Catalase | Sigma Aldrich | Cat # C100; RRID: |
| CellTrace ™ Violet | Thermo Fisher | Cat # C34557; RRID: |

| | | |
|---|---|---|
| Centricon® Plus-70 centrifugal filter units | Millipore | Cat # UFC703008; RRID: |
| Cysteamine MEA | Sigma Aldrich | Cat # 30070; RRID |
| Disposable PD 10 desalting column | G&E Healthcare | Cat # 17-0851-01; RRID: |
| DMEM | Sigma Aldrich | Cat # D6429; RRID: |
| DMSO | Sigma Aldrich | Cat # D8418-100ML; RRID: |
| Doxycycline hyclade | Sigma Aldrich | Cat # D9891-10G; RRID: |
| DpnI | New England Biolabs | Cat # R0176; RRID: |
| EasySep Human CD8 T cell isolation kit | STEMCELL | Cat # 17953; RRID: |
| EcoR1 | New England Biolabs | Cat # R0101; RRID: |
| Endoglycosidase H (endo H) | New England Biolabs | Cat # P0702S; RRID |
| EZ-Link Maleimide-PEG2 biotin | Thermo Fisher | Cat # 21901BID; RRID: |
| Fetal bovine serum (FBS) | Gibco | Cat # 10500-064; RRID: |
| Glucose | Sigma Aldrich | Cat # G8270; RRID: |
| Glucose Oxidase | Sigma Aldrich | Cat # G2133; RRID: |
| Glutamine | Gibco | Cat # A2916801; RRID: |
| HA peptide | Sigma Aldrich | Cat # I2149; RRID: |
| His-Pur Cobalt Resin | Thermo Fisher | Cat # 89964; RRID: |
| His-Streptavidin (AA 25-183) protein (His tag) | Antibodies Online | Cat # ABIN666648; RRID: |
| HLA-A2-SLYNTIATL (6I-A2) monomer | See citation Ref. (Cole et al., 2017) | N/A |
| Human T-activator CD3/CD28 Dynabeads | Thermo Fisher Scientific | Cat # 11161D; RRID: |
| Human serum | Sigma Aldrich | Cat # H4522; RRID: |
| ICAM-1 protein, recombinant human | R&D Systems | Cat # 720-IC-050; RRID: |
| Imidazole | Sigma Aldrich | Cat # I202-500G; RRID: |
| Iodoacetamide | Sigma Aldrich | Cat # I6125; RRID: |
| Ionomycin | Sigma Aldrich | Cat # I0634; RRID: |
| Lysonase (Lysozyme + Benzonase) | Millipore | Cat # 71230-3; RRID: |
| MART-1 peptide (> 95 % purity by HPLC) | Cambridge Peptides Inc. | N/A |
| n-Dodecyl β-D-maltoside (DDM) | Millipore | Cat # 324355-5GM; RRID: |
| Non-essential amino acids solution | Gibco | Cat # 11140050; RRID: |
| Not 1 | New England Biolabs | Cat # R0189; RRID: |
| NuPAGE LDS Sample Buffer | Invitrogen | Cat # NP0007; RRID: |
| NuPAGE Sample Reducing Agent | Invitrogen | Cat # NP009; RRID: |
| *NY*-ESO-1 peptides (> 95 % purity by HPLC) | Cambridge Peptides Inc. | N/A |
| PEG-*it* ™ | SBI | Cat # LV810A-1; RRID: |
| PEIpro | Polyplus | Cat # 115-010; RRID: |
| Pepsin | Thermo Fisher | Cat # 20343; RRID: |
| Phosphate Buffer Saline (PBS) | Sigma Aldrich | Cat # P4417; RRID: |
| Phorbol 12-myristate 13-acetate (PMA) | Sigma Aldrich | Cat # P1585; RRID |
| Phycoerythrin-conjugated streptavidin | Sigma Aldrich | Cat # S4762-5MG; RRID: |
| Pme1 | New England Biolabs | Cat # R0560; RRID: |
| Polybrene | Sigma Aldrich | Cat # 28728554; RRID: |
| Proteases inhibitors (cOmplete, EDTA free) | Roche | Cat # 11873580001; RRID: |
| Protein G agarose | Thermo Fisher | Cat # 15920; RRID: |
| Puromycin | Gibco | Cat # A11138-03; RRID: |
| RPMI 1640 | Sigma Aldrich | Cat # R8758; RRID: |
| Sodium Acetate | Thermo Fisher | Cat # 10000500; RRID: |
| Sodium Fluoride | Sigma Aldrich | Cat # S1504-500G; RRID: |

| | | |
|---|---|---|
| Sodium Orthovanadate | New England Biolabs | Cat # 11873580001; RRID: |
| Sodium Pyruvate | Gibco | Cat # 11360070; RRID: |
| Strep-Tactin Sepharose 50% suspension | IBA Lifesciences | Cat # 2-1201-010; RRID: |
| Streptavidin Sepharose High Performance beads | GE Healthcare | Cat # 17-5113-01; RRID: |
| Tetracycline-free FBS | Clontech | Cat # 631106; RRID: |
| Trans-Blot Turbo Midi Nitrocellulose Transfer Packs | Biorad | Cat # 170-4159; RRID: |
| Trans-Blot Turbo Mini Nitrocellulose Transfer Packs | Biorad | Cat # 170-4158; RRID: |
| Trizma base | Sigma | Cat # T6066 ; RRID: |
| Xba1 | New England Biolabs | Cat # R0145; RRID: |
| Zeba Spin Desalting | Thermo Fisher | Cat # 89889; RRID: |
| 0.45 µm sterile filters | Sartorius Stedim | Cat # 1655-K ; RRID: |
| 8 well chamber Glass µ-slide | Ibidi | Cat # 80821; RRID: |
| 96-well V-bottom plates | Thermo Fisher | Cat # 634-0009; RRID: |
| Critical Commercial Assays | | |
| QIAprep Spin Midiprep Kit | Qiagen | Cat # 27106; RRID: |
| QIAprep Spin Miniprep Kit | Qiagen | Cat # 12243; RRID: |
| BP Clonase II kit | Thermo Fisher | Cat #:11791020; RRID |
| LR Clonase II kit | Thermo Fisher | Cat #:11789020; RRID |
| Experimental Models: Cell Lines | | |
| Human: TCR$\beta^-$ CD8$^+$ Jurkat 31.13; (J13.31*) | See citation Ref.(Alcover et al., 1990) | N/A |
| Human: TCR$\alpha^-\beta^-$/CD8$^-$ Jurkat J76; (CD8-deficient J76*) | See citation Ref.(Heemskerk et al., 2003) | N/A |
| Human: TCR$\alpha^-\beta^-$/CD8$^+$ Jurkat J76; (J76*) | This study | N/A |
| Human: 1G4$\alpha^{WT}\beta^{WT}$ J76 CD8$^-$; (CD8-deficient J76 1G4 WT*) | This study | N/A |
| Human: 1G4$\alpha^{WT}\beta^{WT}$ J76 CD8$^+$; (J76 1G4 WT*) | This study | N/A |
| Human: 1G4$\alpha^{WT}\beta^{A291}$ J76 CD8$^-$; (CD8-deficient J76 1G4 $\beta$A291*) | This study | N/A |
| Human: 1G4$\alpha^{WT}\beta^{A291}$ J76 CD8$^+$; (J76 1G4 $\beta$A291*) | This study | N/A |
| Human: 1G4$\alpha^{WT}\beta^{F291}$ J76 CD8$^-$; (CD8-deficient J76 1G4 $\beta$F291*) | This study | N/A |
| Human: 1G4$\alpha^{WT}\beta^{L291}$ J76 CD8$^-$; (CD8-deficient J76 1G4 $\beta$L291*) | This study | N/A |
| Human: 1G4$\alpha^{WT}\beta^{A293}$ J76 CD8$^-$; (CD8-deficient J76 1G4 $\beta$A293*) | This study | N/A |
| Human: 1G4$\alpha^{WT}\beta^{A303}$ J76 CD8$^-$; (CD8-deficient J76 1G4 $\beta$A303*) | This study | N/A |
| Human: 2H5$\alpha^{WT}\beta^{WT}$ J76 CD8$^-$; (J76 CD8$^-$ 2H5 WT*) | This study | N/A |
| Human: 2H5$\alpha^{WT}\beta^{A291}$ J76 CD8$^-$; (J76 CD8$^-$ 2H5 $\beta$A291*) | This study | N/A |
| Human: 1G4$\alpha^{WT}\beta^{WT}$ CD3$\zeta^{-/-}$ J76 CD8$^+$; (J76-1G4WT-$\zeta$KO*) | This study | N/A |
| Human: 1G4 $\alpha^{WT}\beta^{WT}$ CD3$\zeta^{-/-}$ J76 CD8$^+$ expressing CD3$\zeta$WT; (J76-1G4WT-$\zeta$KO expressing $\zeta$WT*) | This study | N/A |

| | | |
|---|---|---|
| Human: 1G4 $\alpha^{WT}\beta^{WT}$ CD3$\zeta^{-/-}$ J76 CD8$^{+}$ expressing $\zeta$I38A; (J76-1G4WT-$\zeta$KO expressing $\zeta$I38A*) | This study | N/A |
| Human: 1G4 $\alpha^{WT}\beta^{WT}$ CD3-$\zeta^{-/-}$ J76 CD8$^{+}$ expressing $\zeta$I41A; (J76-1G4WT-$\zeta$KO expressing $\zeta$I41A*) | This study | N/A |
| Human: 1G4 wtc51 J76 CD8$^{+}$; (J76 wtc51*) | This study | N/A |
| Human: 1G4 wtc51 J76 CD8$^{-}$; (CD8-deficient J76 wtc51*) | This study | N/A |
| Human: 1G4 QM-$\alpha$ J76 CD8$^{+}$; (QM-$\alpha$*) | This study | N/A |
| Human: 868 J76 CD8$^{+}$; (J76 868*) | This study | N/A |
| Human: 868 J76 CD8$^{-}$; (CD8-deficient J76 868*) | This study | N/A |
| Human CD8$^{+}$ primary T cell 1G4 wtc51 | This study | N/A |
| Human: 1G4 $\alpha^{WT}\beta^{WT}$ CD3$\zeta^{-/-}$ expressing CD3$\zeta$-TST (J76 1G4 $\zeta$-TST*) | This study | N/A |
| Human: HEK293T | ATCC | Acc. #: CRL-3216 |
| Human: Lenti-X293T | Clontech | Cat. #: 632180; RRID: |
| *: indicates nomenclature used within the text | | |
| Experimental Models: Organisms/Strains | | |
| *E. coli* Stbl3 Competent Cells | Invitrogen | Cat. #: C7373-03 |
| *E. coli* DH5$\alpha$ Competent Cells | TermoFisher Scientific | Cat. #: 18265-017 |
| Oligonucleotides | | |
| See Table S2 | | |
| Recombinant DNA | | |
| pDONR-221 | Thermo Fisher scientific | Cat # 12536017; RRID |
| pEF3 FLAG | K. Nika, University of Oxford | N/A |
| pEF3 HA | K. Nika, University of Oxford | N/A |
| pEXPR-IBA103 | IBA Lifesciences | Cat # 2-3503-000; RRID |
| pHR-CD8$\alpha\alpha$ | A. Van Der Merwe, University of Oxford | N/A |
| pHR-SIN-BX-IRES-Emerald | V. Cerundolo, University of Oxford | N/A |
| pLIX-402 | Addgene | Cat # 41394; RRID |
| pLEX_307 | Addgene | Cat # 41392; RRID |
| psPAX2 | Addgene | Cat # 12260; RRID |
| pVSV-G | Addgene | Cat # 14888; RRID |
| Software and Algorithms | | |
| Flow Jo | BD | N/A |
| GROMACS | http://www.gromacs.org/ | N/A |
| Image Studio | Li-COR Bioscience | N/A |
| Insight3 | B. Huang, University of California, San Francisco | N/A |
| Jalview | See citation Ref. (Waterhouse et al., 2009) | N/A |
| MATLAB software | S. Shelby and S. Veatch, University of Michigan | N/A |
| OriginPro 2017 | OriginLab | N/A |
| Prism | GraphPad Software | N/A |
| PyMOL | Schrödinger LLC | N/A |
| SnapGene | GSL Biotech LLC | N/A |
| Other | | |
| Hi-Load 16/600 Superdex 200 pg Column | GE Healthcare | Cat # 28-9893-35; RRID: |

| | | |
|---|---|---|
| Superdex-200 Increase 5/150 GL Column | GE Healthcare | Cat # 28-9909-44; RRID: |
